# *Bacteroides* expand the functional versatility of a universal transcription factor and transcribed DNA to program capsule diversity

**DOI:** 10.1101/2024.06.21.599965

**Authors:** Jason Saba, Katia Flores, Bailey Marshall, Michael D. Engstrom, Yikai Peng, Atharv S. Garje, Laurie Comstock, Robert Landick

## Abstract

Human gut *Bacteroides* species encode numerous (eight or more) tightly regulated capsular polysaccharides (CPS). Specialized paralogs of the universal transcription elongation factor NusG, called UpxY (Y), and an anti-Y UpxZ (Z) are encoded by the first two genes of each CPS operon. The Y-Z regulators combine with promoter inversions to limit CPS transcription to a single operon in most cells. Y enhances transcript elongation whereas Z inhibits noncognate Ys. How Y distinguishes among cognate CPS operons and how Z inhibits only noncognate Ys are unknown. Using in-vivo nascent-RNA sequencing and <u>p</u>romoter-less in <u>v</u>itr<u>o</u> transcription (PIVoT), we establish that Y recognizes a paused RNA polymerase via sequences in both the exposed non-template DNA and the upstream duplex DNA. Y association is aided by novel ‘pause-then-escape’ nascent RNA hairpins. Z binds non-cognate Ys to directly inhibit Y association. This Y-Z hierarchical regulatory program allows *Bacteroides* to create CPS subpopulations for optimal fitness.

## INTRODUCTION

*Bacteroides* are abundant and crucial members of the modern human gut microbiota. A key evolved feature of these bacteria is the ability of each strain to produce numerous (eight or more) distinct capsular polysaccharides (CPS)^1,2^ that are tightly regulated so that only one CPS is typically produced per bacterial cell. This bet-hedging strategy generates *Bacteroides* populations with great surface variability that protect from phage^3–5^ while also mediating other processes such as immune modulation, biofilm formation, and affecting antibiotic resistance and inflammation^6–11^.

CPS diversity is achieved by regulating both transcription initiation and elongation of CPS biosynthesis operons. *Bacteroides fragilis* (*Bfr*) has eight distinct CPS operons, producing PSA-PSH. All but PSC use invertible promoters and all encode *upxY* (Y_X_) and *upxZ* (Z_X_) paralogs as the first genes in each operon^12,13^. The fraction of each promoter oriented ON versus OFF varies with environmental conditions^14^. CPS promoter inversions are stochastic and multiple CPS promoters are oriented ON in most cells simultaneously^15–17^. *Bacteroides* prioritize expression of one promoter-ON CPS operon over others by regulating RNA polymerase (RNAP) elongation via the operon-specific Y_X_ elongation activator and Z_X_ inhibitor of non-cognate Y_X_. Z_X_ inhibits a subset of non-cognate Y_X_ possibly via direct binding (e.g., Z<u>_A_</u> from PS<u>A</u> may inhibit Y<u>_E_</u> from PS<u>E</u>). *Bfr* Y_X_ paralogs must distinguish among eight target CPS loci to enable operon-specific regulation, but how this discrimination is accomplished is unknown.

Y_X_ family proteins are specialized (i.e., locus-specific) paralogs of NusG/Spt5, the only universal transcription factor found in archaea, eukaryotes, and bacteria^18^. NusG-family regulators bind RNAPs during transcript elongation and modulate RNAP activity through interactions with the RNAP and the surface-exposed nontemplate DNA strand^19–21^. Globally acting *E. coli* NusG and its single specialized paralog RfaH increase elongation rate and decrease pausing^22–25^. In contrast, *B. subtilis*, *M. tuberculosis*, and *T. thermophilus* NusGs enhance both pausing and intrinsic termination^26–30^. Pausing during transcript elongation is a universal regulatory feature of RNAPs that allows site-specific recruitment of transcription factors (TFs)^31^ and guides RNA synthesis.

Among the many known NusG_SP_ families, RfaH of Proteobacteria is the best understood. RfaH targets operons that contain a DNA element called *ops* (<u>o</u>peron <u>p</u>olarity <u>s</u>uppressor) in their leader regions (DNA between the transcription start site and the translation start codon of the first gene). RNAP pauses at the 12-nucleotide *ops*, allowing RfaH to associate via sequence-specific interactions with a non-template strand DNA hairpin (ntDNAhp) exposed by the paused RNAP. Other NusG_SP_ include LoaP in Firmicutes/Bacillota^21^, TaA in Myxococcota^32^, and plasmid-encoded ActX in Pseudomonadota^33^.

The CPS operon leader regions are required for Y-mediated regulation^12^, consistent with sequence-specific Y_X_ recruitment to RNAP paused in this region (Fig. 1A). In principle, Y_X_ could recognize ntDNA (like RfaH), nascent RNA (like LoaP), or both to discriminate among multiple, similar CPS operon targets. We used both in vivo and in vitro analyses to identify pauses in CPS operon leader regions, establish that these pause sites function as recruitment sites for Y, and discover novel NusG_SP_–DNA interactions and mechanisms that mediate Y–CPS operon specificity. We found that Z directly binds noncognate Ys to block Y action and that differential Y_X_–Z_X_ affinities enable CPS hierarchical control of transcript elongation. These results define mechanisms that explain the exquisite specificity of multiple NusG_SP_ and that allow *Bacteroides* to program CPS diversity in the highly dynamic human gut environment.

**Figure 1.**
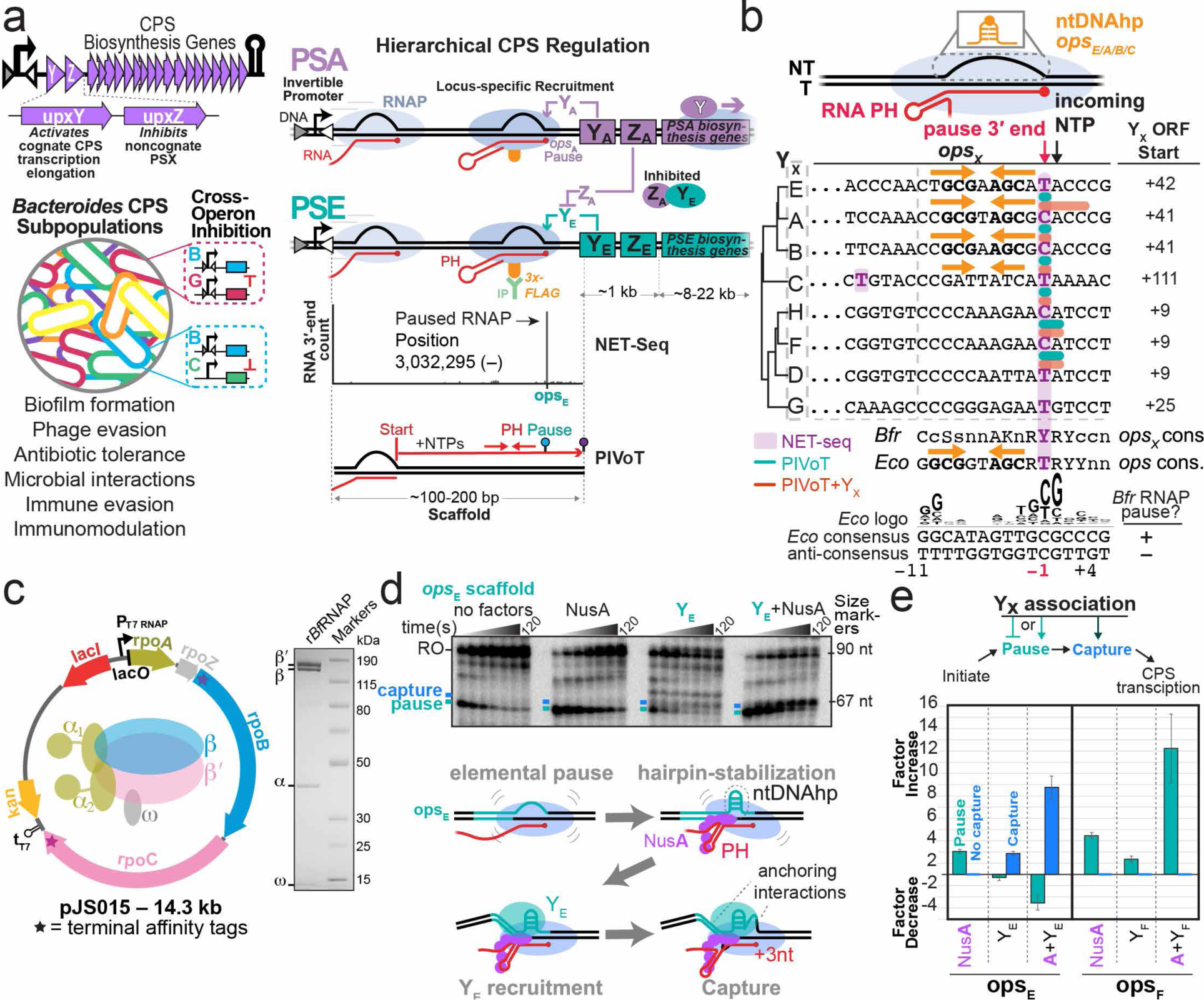
*Bacteroides fragilis* RNAP pauses in CPS operon leader regions in vivo and in vitro at candidate Y_X_ recruitment sites called *ops*_X_. **a)** Representative CPS operon diagram highlighting Y_X_ and Z_X_, the first two genes in *B. fragilis* PSX operons. Horizontal triangles mark the inverted repeats recognized by Mpi recombinase for promoter inversion^17^. Proposed roles for CPS diversity in *B. fragilis* subpopulations (colored coats) are listed^3,6,8–10,99^. The schematics illustrate the proposed roles of Y_X_ activation and Z_X_ inhibition of noncognate Y_X_ in generating subpopulation CPS diversity^12,13^. Y_X_ is recruited to RNAP paused at cognate but not non-cognate *ops*_X_ sites that encode a pause hairpin (PH). Z_X_ directly binds Y_X_ from heterologous operons and inhibits its recruitment. In vivo (NET-seq) and in vitro (PIVoT) methods for identifying RNAP pause sites (*ops*_X_) are illustrated. **b)** Transcriptional pauses in CPS leader regions identified in this study are shown in comparison to the RfaH *ops* pause and the *E. coli* consensus elemental pause sequences^35^. T=template strand; NT=non-template strand. Fully conserved nucleotides are capitalized; largely conserved nucleotides are lowercase. **c)** r*Bfr*RNAP overexpression plasmid and final purified RNAP separated by SDS-PAGE and Coomassie-stained. Stars indicate terminal affinity tags. **d)** PIVoT assay of PSE promoter-distal leader regions. Assays included 1 µM NusA or 150 nM Y_E_ added concomitantly with NTPs as indicated. RNAs from a reaction time course were separated by 8% Urea-PAGE. **e)** NusA and Y_X_ synergistic activities at cognate *ops_X_* sites. Y_X_ association manifests as pause inhibition or pause enhancement (aqua bars), or capture (blue bars). Error bars represent standard deviations from triplicate assays.

## RESULTS

### *Bacteroides fragilis* RNAP pauses in CPS operon leader regions in vivo and in vitro at candidate Y_X_-recruitment sites (*ops*_X_)

Specific Y_X_ recruitment sites likely exist in CPS leader regions because these leader sequences are variable and are required for Y_X_ activity^12^. Since *Eco*RfaH is recruited to RNAP at leader region *ops* pause sites, we first asked if *Bfr*RNAP pauses in the leader regions of CPS operons. To identify candidate Y_X_-recruiting pause sites directly in vivo, we used <u>n</u>ascent <u>e</u>longating transcript <u>seq</u>uencing (NET-seq) (Extended Data Fig. 1a; Fig. 1a,b). NET-seq allows genome-scale identification of precise nascent RNA 3′ ends, which are enriched at pause sites^34,35^.

Remarkably, NET-seq revealed single prominent pause sites in most CPS operon leader regions (Fig. 1a,b; Extended Data Fig. 1b)^35^. Eight CPS leader pauses exhibited an obvious consensus sequence typical of strong *Eco* pauses and appear to be type-1 pauses,^36^ meaning pauses stimulated by nascent RNA pause hairpins (PHs) that allosterically inhibit RNAP activity^37,38^. Pausing in the PSC leader region (the only *Bfr* CPS operon with a non-invertible, constitutively ON promoter)^39^ occurred at multiple sites; weak pausing occurred at a site resembling the other seven in sequence and location (Fig. 1b and Extended Data Fig. 1b). We designated the CPS leader pause sites *ops*_X_ (‘X’ designates the CPS operon) based on analogy to the RfaH *ops* site.

To test whether the *ops*_X_ pause recruits Y_X_, we generated recombinant *Bacteroides fragilis* RNAP (r*Bfr*RNAP) and assayed CPS leader regions using <u>p</u>romoter-less in <u>v</u>itr<u>o</u> transcription (PIVoT)^40,41^. PIVoT bypasses the need for σ^A^-dependent initiation (Fig. 1a,c; Extended Data Fig. 2a). We first asked if r*Bfr*RNAP recognizes the consensus elemental pause signal defined for *Eco*RNAP (Fig. 1b)^35^. Signals resembling this consensus direct pausing by a wide variety of RNAPs from bacteria to human^35,42,43^. Bacterial pause sequences are reported to differ in some species^44,45^ and have not been tested for Bacteroidota. We found that r*Bfr*RNAP pauses strongly at the consensus sequence but not anti-consensus sequence (Extended Data Fig. 2a,b,c), suggesting its pause signals resemble those of *Eco*RNAP and most other tested RNAPs.

We next assayed pausing in a representative subset of CPS leader regions. Strikingly, the PSA, B, E, F and H leader segments encoded single prominent pause sites that corresponded exactly to the sites found by NET-seq (Fig. 1b,d; Extended Data Figs. 2 and 3, Supplementary Fig. 1). Pausing was less prominent but detectable at *ops*_C_, consistent with the heterogeneous pausing observed the NET-seq. We conclude that CPS operon leader regions encode strong pause sites for RNAP with similar but not identical sequences, as might be expected for Y_X_ recruitment sites that must distinguish among Y_X_ paralogs.

To ask if the CPS leader pauses function as targets for Y_X_ recruitment, we purified Y_A_, Y_B_, Y_C_, Y_E_, Y_F_, and Y_H_ (Methods) and tested their effects on pausing using PIVoT. In *Eco* and *Bsu*, NusA stimulates pausing in part via contacts to PHs^35,37,41,46–48^. Thus, we also purified *Bfr*NusA and tested for NusA synergies with Y_X_. Intriguingly, Y_A,B,E_ inhibited the cognate leader pause, whereas Y_C,F,H_ enhanced the cognate leader pause. Y_E_ additionally trapped a fraction of RNAP just downstream from the pause site, as seen previously with *Eco*RfaH (Fig. 1d,e, labeled ‘capture’) (Extended Data Fig. 3). Thus, Y_X_ association with paused elongation complexes (PECs) may manifest as either pro-pausing or anti-pausing activity.

All six CPS leader pauses were greatly enhanced by NusA additively with the Y_X_ effects (Extended Data Fig. 3). Importantly, the effects of Y_X_ were specific to the NET-seq identified leader pauses, consistent with *ops*_X_ sites functioning as specific Y_X_-recruitment sites. We conclude that the NET-seq-identified leader pauses are bona fide target sites for Y_X_ association with *Bfr*RNAP. Notably, *ops*_A,B,E_ encode putative ntDNAhps at [–11 to +1] that resemble the *ops* ntDNAhp known to recruit *Eco*RfaH (5′-<u>GCG</u>–<u>AGC</u> stems; Fig 1b, Extended Data Fig. 1c). The *Bfr ops*_X_ ntDNAhp sequences differ, consistent with specific recruitment of cognate Y_X_. However, *ops*_F,H_ are identical in the ntDNAhp region, suggesting that some other element contributes to specificity.

RNAP capture by Y_X_–*ops*_X_ interaction, which is evident for *ops*_E_ but not *ops*_A_ or *ops*_B_ by accumulation of RNAs a few nucleotides longer than the primary *ops*_E_ pause RNA (Extended Data Fig. 3), suggests some but not all *ops*_X_ sites exhibit pause cycling^31,49,50^. Pause cycling occurs when the ntDNA is captured by a regulator that also contacts RNAP (e.g., *Eco*σ^70^ or RfaH), anchoring the paused elongation complex (PEC) and hindering extension beyond 2-3 nt^51,52^. Trapped PECs can be rescued by RNA cleavage factors GreA,B^50^, creating a cycle that repeats until ntDNA contacts rearrange to allow normal elongation^49^.

Importantly, even in the presence of globally acting *Bfr*NusG, Y_F_ still enhances *ops*_F_ pausing (Extended Data Fig. 4). Thus, Y_X_ appears to outcompete *Bfr*NusG even though both NusG and its specialized paralog Y_X_ use the same primary binding site on RNAP.

### Z_X_ inhibits Y_X_ at *ops*_X_ through direct Z_X_–Y_X_ interaction

We next sought to confirm that Y_X_ associates with *ops*_X_ PECs and to test whether Y_X_ binding requires sequence upstream of the putative ntDNAhp region using in vitro binding, in silico interaction, and in vivo gene expression assays. Y_E_ is predicted to be inhibited by Z_A_ but not by Z_E_ or Z_C_ in a strain with only the PSA, PSE, and PSC promoters oriented ON (expression hierarchy PSA>E>C)^13,17^. We call this strain [AE]_ON_ for simplicity because the PSC promoter is constitutive^13^. To test our prediction, we measured Z_A_–Y_E_ and Z_E_–Y_E_ binding constants by biolayer interferometry (BLI) (Fig. 2a,b). Z_A_ but not Z_E_ bound tightly to Y_E_ (*K*_D_ ∼0.9 nM vs ∼88 nM). We conclude that Z_X_ acts through direct Y_X_ binding.

**Figure 2.**
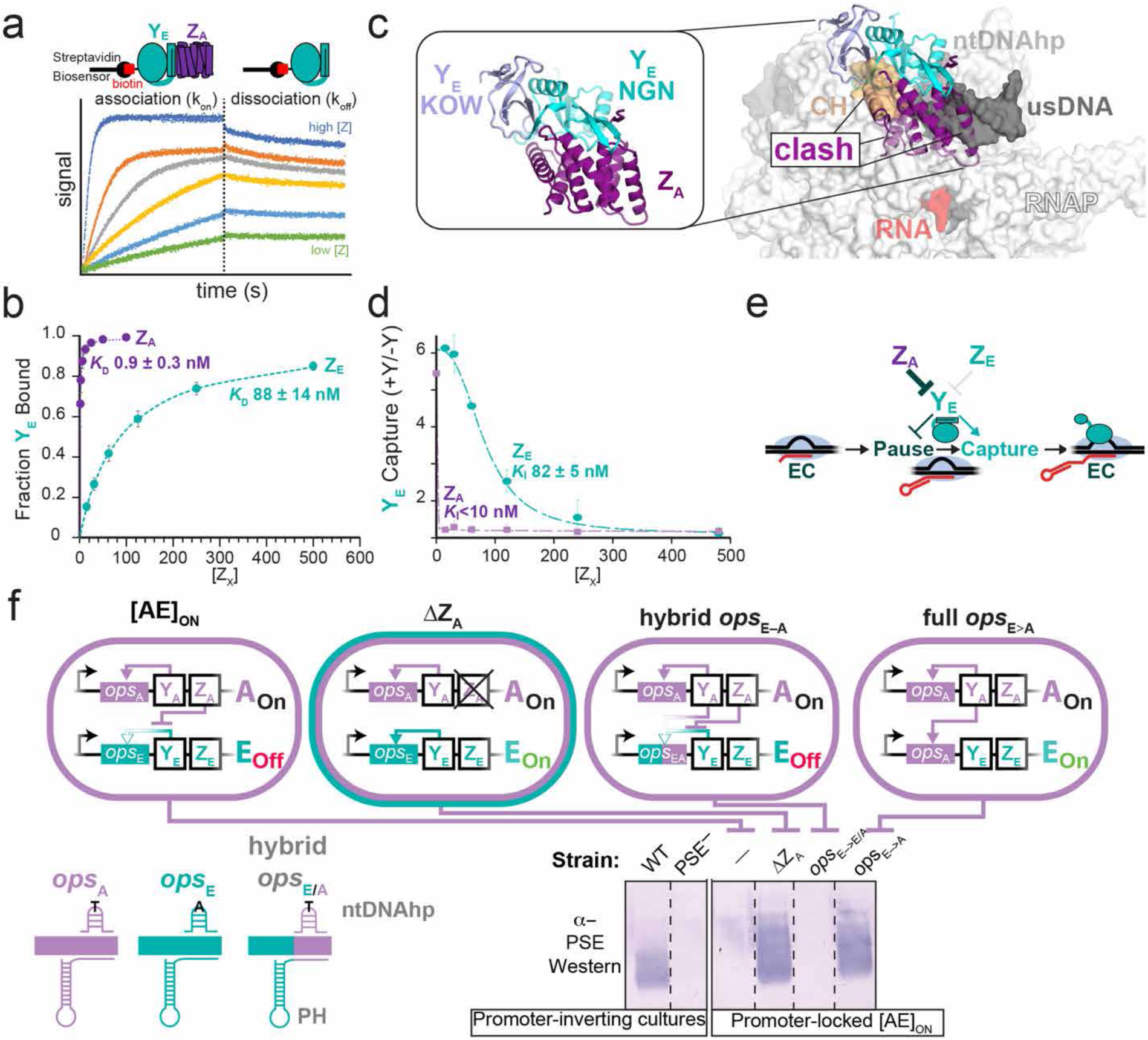
*ops*_X_ pause sites are recruitment sites in vivo that enable Y_X_-locus specificity, CPS hierarchical control, and can be re-wired to bypass direct inhibition by Z_X_. **a)** Z_X_ directly binds Y_X_ as revealed by biolayer interferometry (BLI^100^) over a range of Z_X_ concentrations yielding biotin-Y_E_–Z_X_ on and off rates: Y_E_–Z_A_ k_on_ = 1.1×10^6^ ± 2.9×10^5^ M^−1^s^−1^, k_off_ = 9.0×10^−4^ ± 2.4×10^−4^ s^−1^; Y_E_–Z_E_ k_on_ = 1.1×10^5^ ± 2.5×10^4^ M^−1^s^−1^, k_off_ = 9.6×10^−3^ ± 7.3×10^−4^ s^−1^. Assays were performed in triplicate and globally fit to a 1:1 binding model (see Methods). Reported *K*_D_s are averages from three independent global fits and errors represent standard deviations. **b)** Y_E_–Z_A_ and Y_E_–Z_E_ binding curves calculated from BLI results. **c)** AlphaFold3^53^ model of Y_E_–Z_A_ and steric clash evident when Y_E_-Z_A_ is aligned to an RfaH-bound PEC^23^. **d)** Fold change in Y_E_ capture as a function of cognate Z_E_ or non-cognate Z_A_ concentration in PIVoT assays (Methods). Subsets of NusA, Y_E_, variable [Z_X_], and NTPs were added to initiate pause assays (50 nM Y_E_, 15-480 nM Z_X_, 1 µM NusA, final). **e)** Model for Z_X_ inhibition of Y_X_ recruitment. **f)** Strain background used in *ops*_X_ replacement experiments are depicted (Δ*mpi* M44 in each strain ensured only the PSA, PSC, and PSE promoters are oriented ON). In WT, promoter orientations are variable in single cells, but some cells express PSE. PSE^−^ is an insertion mutant that abrogates PSE expression. In promoter-locked [AE_ON_] strains, only YA activated genes are expressed because of cross-operon inhibition of Y_E_ and Y_C_ by Z_A_. Strains with partial [-10:-1]_E_ or full [-38:-1]_E_ segments of *ops*_E_ were replaced with their *ops*_A_ counterparts ([-10:-1]_A_ and [-43:-1]_A_) and assayed for their ability to rescue PSE expression by Western blot.

To understand how Z_A_ might interact with Y_E_, we predicted their association using AlphaFold 3^53^ (Fig 2c, Extended Data Fig. 5). The Z_A_–Y_E_ complex, which was predicted with high confidence, placed Z_X_ on the RNAP-binding interface of Y_E_. When modeled into an *Eco*RNAP-RfaH-*ops*-PEC (PDB 8PHK)^50^ by alignment of the Y_E_ NGN domain with the RfaH NGN, Z_A_ clashed with two major PEC features: (*i*) the RNAP clamp helices (CH), which provide the primary RNAP binding site for of all NusG-family regulators (Fig. 2c, orange); and (*ii*) the proximal upstream DNA duplex (usDNA). Thus, Z_X_ likely inhibits Y_X_ by preventing its recruitment to RNAP at *ops_X_* pause sites.

We next used PIVoT to test whether Z_A_ or Z_E_ blocked Y_E_ inhibition of pausing at the candidate *ops*_E_ pause site as predicted by the AlphaFold model. Z_E_ blocked Y_E_ action only at high concentrations (*K*_I_ approximating the *K*_D_ measured by BLI; Fig. 2d). In contrast, Z_A_ inhibited Y_E_ at all tested concentrations. We conclude that differential Y_X_–Z_X_ affinities enable CPS hierarchical control of transcript elongation (Fig. 2e).

### Y_X_ targets extended *ops*_X_ sites in vivo

Using these insights into Z_X_–Y_X_ interaction, we tested whether *ops*_X_ pause sites function as Y_X_ recruitment sites in vivo and which sequences govern cognate Y_X_ function. Using a constitutive [AE]_ON_ strain^17^, we replaced *ops*_E_ segments with the corresponding *ops*_A_ segments. We predicted that the *ops*_E_–*ops*_A_ swapped strain should activate PSE expression because Y_A_ should bind *ops*_A_ in PSE. To ask if the PH-encoding region of *ops*_X_ is required for Y_X_ recruitment, we also constructed a hybrid *ops*_E–A_ strain in which only the ntDNAhp region corresponding to the RfaH *ops* but not the PH-encoding region of *ops*_E_ was replaced with *ops*_A_ sequence (Fig. 2f). Using antibodies confirmed to detect PSE in a WT strain but not in a PSE^−^ mutant, we tested for PSE expression in [AE]_ON_ and derivative strains: ΔZ_A_, hybrid *ops*_E–A_, and full *ops*_E→A_ (Fig. 2f). PSE was (*i*) not expressed in [AE]_ON_; (*ii*) expressed in ΔZ_A_; (*iii*) not expressed in the hybrid *ops*_E–A_ strain; and expressed in the full *ops*_E→A_ swapped strain.

To confirm that the upstream PH-encoding region is required for Y_X_ action, we also tested Y_A_ and Y_E_ effects similarly using PIVoT (Extended Data Fig. 6a,b). Neither Y_A_ nor Y_E_ modulated pausing or PEC capture at WT levels unless the full cognate *ops*_X_ including the upstream PH-encoding region was present. Thus, both in vivo and in vitro, the cognate upstream PH-encoding region is required for full Y_X_ activity.

We conclude that *ops_X_* is comprised of both the ntDNAhp region and the upstream PH-encoding region. These regions are necessary and sufficient to program Y_X_ recruitment and enhancement of CPS-operon transcription. The inactivity of Y_X_ at hybrid sites establishes that the ∼40 bp *Bacteroides* CPS *ops_X_* sequences differ fundamentally from the RfaH *ops* that requires only a 12-bp ntDNAhp sequence. Additional recognition of the upstream PH-encoding region likely aids Y_X_ discrimination among target sites. However, determining whether these upstream sequences contact Y_X_ as a nascent RNA hairpin, as proposed for LoaP^54^, or as duplex DNA required further experimentation.

### Y_X_–*ops*_X_ pairs can be divided into distinct classes

To ask if the variability in *ops*_X_ sequences could be related to variability in Y_X_ paralogs, we compared their apparent evolutionary relationships to sequence and structural alignments of Y_X_, RfaH, and NusGs (Fig 3a, Extended Data Fig. 7). Strikingly, both Y_X_ protein and *ops*_X_ DNA sequences clustered into two distinct classes with two outliers (anti-pausing Class-1, PSA,B,E; pro-pausing Class-2, PSD,F,H; Outliers PSG,C) (Fig. 3b; Extended Data Fig. 7). We use the *ops*_X_ pause site defined as position –1 as a reference in this analysis. Class-1 DNA–RNA sequences exhibited several key features: (*i*) an apparent ntDNAhp (orange arrows); (*ii*) an apparent PH that extends to –12 to –9 (red arrows; relative to –1 pause RNA 3′ nucleotide position); and (*iii*) the Y_X_ gene start codon is at +41,+42. Class-1 Y_X_ protein sequences (Fig. 3a) exhibited (*i*) an identical β2–β3 hairpin sequence in the NGN domain (LPTQFVIRQLYKRR[R/K]RVEVP); (*ii*) variable sequences (pink) in NGN α1 and α2 that contact the *ops* ntDNAhp (yellow), RNAP protrusion, and RNAP gate loop; and (*iii*) variability in the C-terminal KOW domain (Fig 1a, Extended Data Fig. 7). The variable Y_X_ sequences in contacts to the ntDNAhp, protrusion, and gate loop are consistent with Y_X_ recognition and potential effects on pausing^27,55,56^.

**Figure 3.**
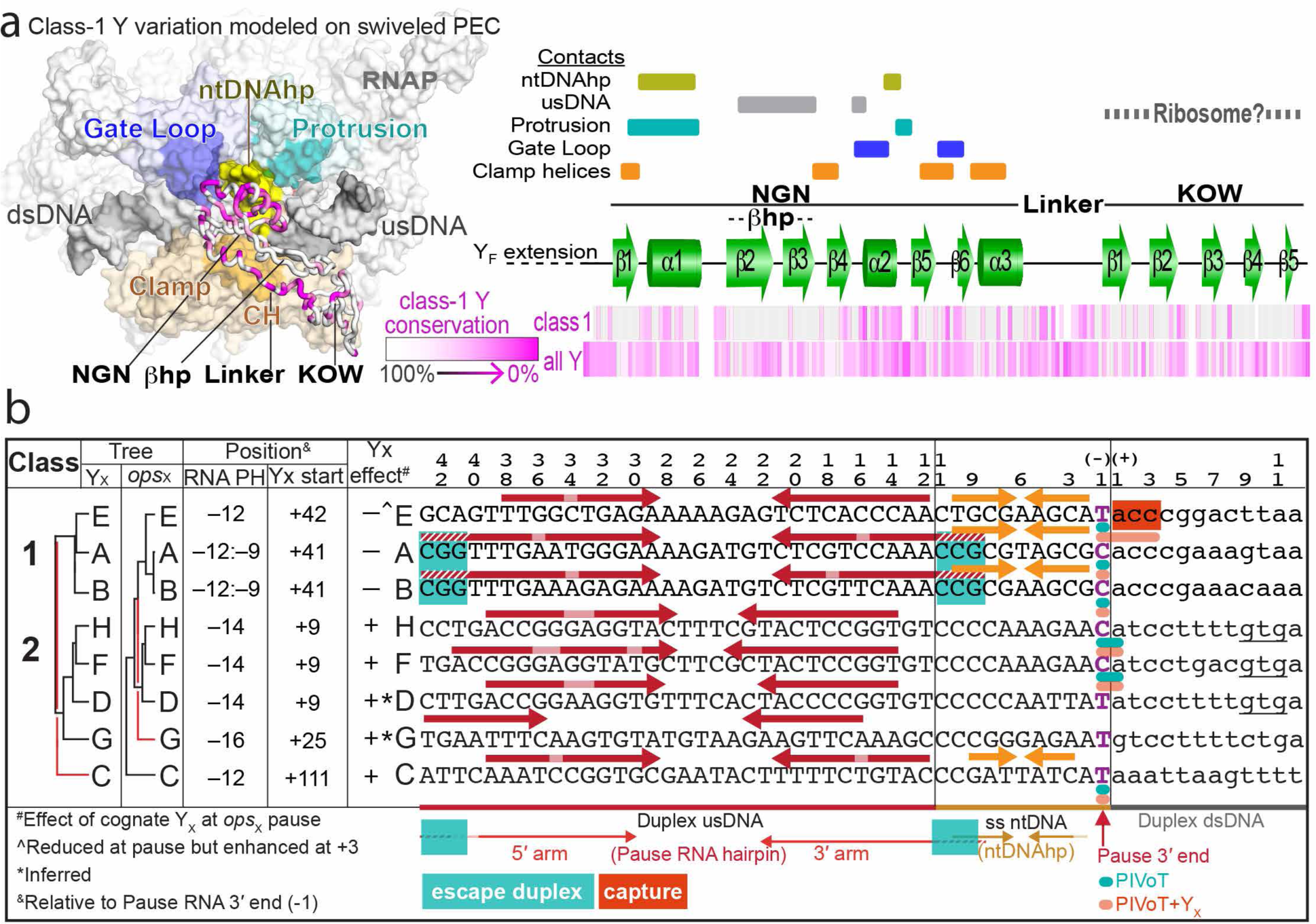
Y_X_ can be divided into distinct classes. **a)** Analysis of sequence conservation and solvent accessibility of CPS operon Y_X_s from *B. fragilis* (Genbank accession NC_003228.3; strain NCTC 9343). Known contacts to RNAP modules or DNA are based on structures of *E. coli* RNAP in complex with NusG (PDB 6C6U)^23^ or RfaH (PDB 6C6S)^23^ (bars on right). The structural model, based on PDB 8PHK^50^, depicts key NusG_SP_-interacting modules of a PEC (gate loop, protrusion, clamp, ntDNAhp, upstream and downstream DNA (usDNA and dsDNA) and features of NusG_SP_ (NGN, KOW, hairpin). The extent of sequence conservation of among Class-1Y_X_ is shown on a magenta color scale mapped to RfaH in the 8PHK model and also compared linearly to conservation among all Y_X_ proteins. (**b)** Sequence comparisons among *ops*_X_ annotated with features relevant to pausing and Y_X_ action compared to phylograms^101^ of Y_X_ and *ops*_X_ shown on right. The red lines indicate the alternative clustering of *ops*_C_ and *ops*_G_ versus Y_C_ and Y_G_ relative to the uniform clustering of other Y_X_ and *ops*_X_ sequences into Class 1 and Class 2.

The Class-1 PSA,B PHs have greater potential to extend towards the pause RNA 3′ end (teal highlight) relative to the PSE PH. Extension of PHs past –10 is thought to destabilize PECs at intrinsic terminators^57^, but we do not observe termination at these sites. An alternative role of PHs extending past –10 could be to aid PEC escape from pause cycles if auxiliary factors like GreA,B are insufficient. Thus, we postulated that base-pairing of the PSA,B PHs at –11,–10, and –9 could explain why Y_A_ and Y_B_ (but not Y_E_) did not capture PECs in pause cycles (Extended Data. Fig 3, Fig. 3b red highlight) (see next section). Based on an apparent ability to prevent PEC capture by Y_X_, we call this PH extension the escape duplex (ED).

Pro-pausing Class-2 (PSD,F,H) sequences exhibited features that differed from Class-1 (Fig 3b; Extended Data Fig. 7). For Class-2 DNA-RNA: (*i*) *ops*_X_ lacks an obvious ntDNAhp; (*ii*) the apparent PH extends only to –14; and (*iii*) the Y_X_ gene start codon is at +9 relative to *ops_X_*. For Class-2 Y_X_: (*i*) the β2–β3 hairpin sequence is variable with pattern of basic residues distinct from Class-1; (*ii*) NGN α1 and α2 also are variable but distinct from Class-1 and thus consistent with differential recognition and different effects on pausing; and (*iii*) the KOW domain exhibits greatly increased positive charge relative to Class-1 (Extended Data Fig. 7).

PSC,G were outliers whose Y_X_ and *ops*_X_ clustered differently relative to Class-1,2. Their apparent PHs extended to –12 or –16, respectively. The Y_X_ start codons were at +111,+25 and both Y_X_ sequences were relatively divergent compared to Class-1,2. Y_C_ enhanced rather than inhibited the *ops*_C_ pause (Extended Data Fig. 3). Class-2 Y_X_ and PSC Y_C_ exhibited charge similarity to the LoaP KOW proposed to bind RNA hairpins (Extended Data Fig. 7).

We conclude that Y_X_ regulators diverged during evolution to form at least two distinct classes within which the interactions that determine Y_X_–*ops_X_* specificity and pro- vs. anti-pausing action appear to have followed different trajectories.

### *ops_X_* PHs stabilize PECs but also can aid escape of PECs captured by Y_X_-DNA contacts

We next sought to assess the function of the putative *ops*_X_ PHs (Fig. 3b). We focused on Class-1 *ops*_X_ to investigate the impact of PH and ED (Fig. 3b, Supplementary Fig. 2). The strong effect of NusA on Class-1 pauses (Fig 1e, Extended Data Fig. 3) made it likely the PHs stimulate pausing^37,41,46–48,58^. Further, removal of the PH-encoding region from an *ops_E_* scaffold eliminated NusA-stimulation of pausing (Extended Data Fig. 8a). To probe the functions of the conventional *ops*_E_ PH and the unconventional *ops*_B_ PH+ED, we used complementary antisense oligonucleotides (asDNAs or asRNAs) to progressively disrupt the 5′ arm of the PSE,B PHs (Fig. 4a,c).

**Figure 4.**
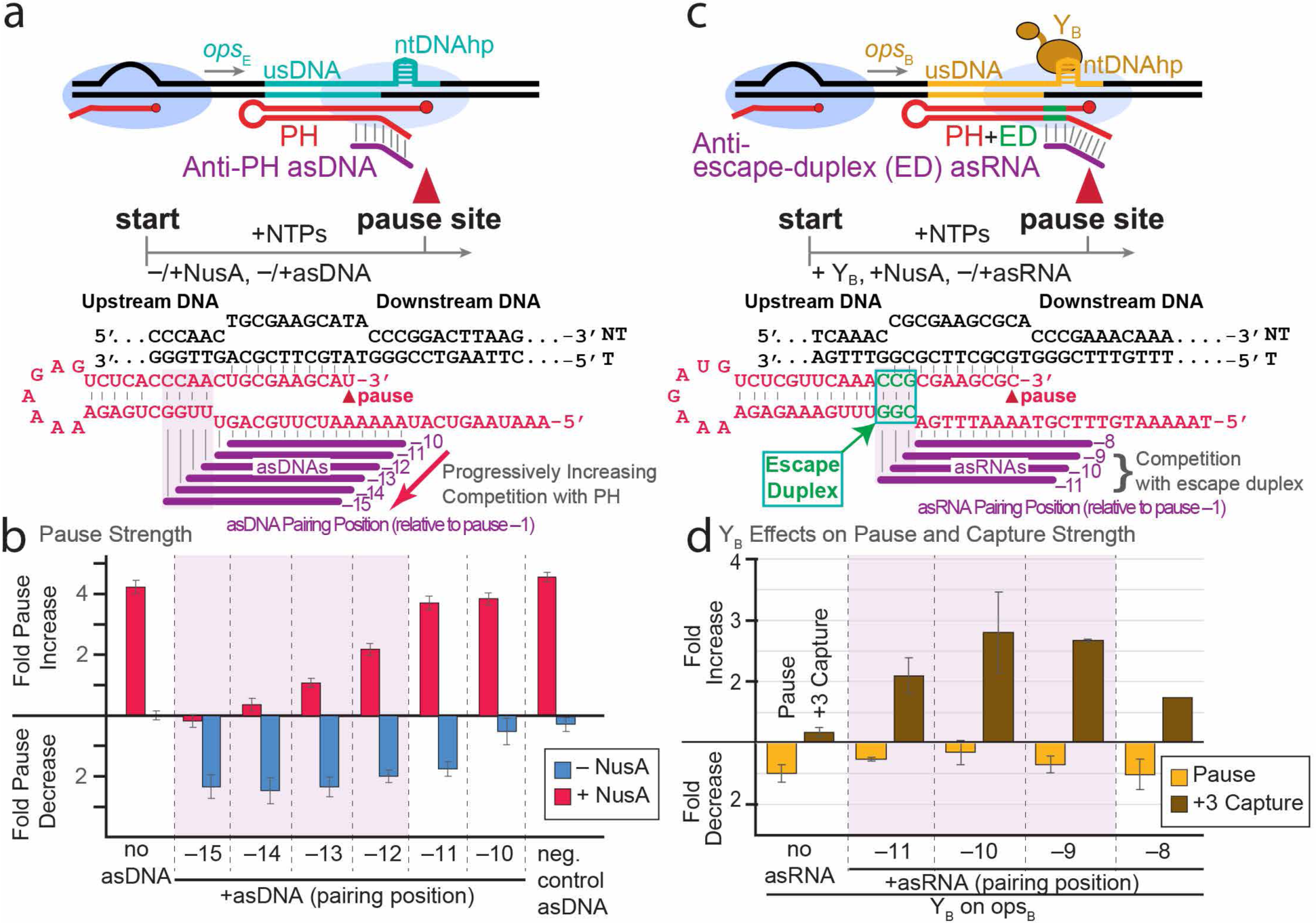
Nascent RNA hairpins promote pausing or pausing-then-escape at *ops*_X_. a) Experimental scheme. r*Bfr*RNAP was reconstituted upstream of the *ops*_E_ pause, enabling PH formation upon RNA extension. **b)** Antisense DNA (asDNA; 10 µM final) effects on NusA enhancement of PH-stimulated *ops_E_* pausing, where different asDNAs disrupt PH formation to different extents. asDNA oligonucleotides were added concomitantly with NusA (or storage buffer) and NTPs (1µM and 100 µM each NTP, final). Error bars are standard deviations from three experiments. **c)** Experimental scheme. r*Bfr*RNAP was reconstituted upstream of *ops*_B_ pause, enabling PH formation. **d)** Antisense RNAs (asRNAs; 0.5 µM final) pairing with an escape duplex (ED, green; the EC is unique to PSA and PSB). asRNAs inhibited PEC escape, leading to accumulation of a captured RNA (*ops*_B_+3 nt; e.g., see Extended Data Fig. 2e). Assays were performed in the presence of 1 µM NusA and 150 nM Y_B_. Error bars are the range of result from duplicate assays of amount RNA paused or captured 45 s after addition of NTPs. Fold changes are relative to plus NusA, no Y_B_, no asRNA).

asDNAs that disrupt the PSE PH by pairing with the 5′ arm but not those that pair just upstream reduced pausing both in the absence and presence of NusA (Fig. 4b). Thus, the PH alone stimulates pausing at *ops*_X_ and *Bfr*NusA significantly stimulates pausing in a PH-dependent manner. We conclude that *ops_X_* sites are type-1 pauses that encode NusA-stabilized PHs, in notable contrast to the type-2 RfaH *ops* that lacks a PH^36^.

To test the idea that the apparent escape duplex (ED) could aid escape of PECs, we measured the effect on capture of antisense RNAs (asRNAs) that disrupt the ED by pairing to the distal bases of 5′ arm of the ops_B_ PH. *ops*_B_ but not *ops*_E_ encodes an ED, and Y_B_ does not cause PEC capture in contrast to Y_E_ (Fig 4cd, Extended Data Fig 3). Addition of asRNAs that progressively disrupted the ED caused Y_B_ to capture PECs in pause cycles. Thus, *ops*_B_, and by analogy *ops*_A_, PHs not only stimulate *ops*_X_ pausing synergistically with NusA to allow time for Y_X_ recruitment, but also use an ED to drive forward translocation at the pause. The ED breaks extensive contacts by Y_X_ necessary for its initial recruitment but problematic for subsequent EC escape.

### Y_X_ distinguishes PECs via multipartite NGN interactions with exposed ntDNA and upstream duplex DNA

We next sought to determine how Class-1 Y_X_ proteins distinguish cognate vs. non-cognate *ops*_X_ sites via the PH-encoding region (Fig. 2). Since the ntDNA of *ops*_E_ and *ops*_B_ are most similar, particularly at the key –6 ntDNAhp position (Fig 3b, Extended Data Fig. 8b), we reasoned that the contribution of sequences upstream from the ntDNAhp might be most apparent by swapping regions between *ops*_E_ and *ops*_B_. We used PIVoT to measure Y_X_ effects on NusA-stimulated pausing and capture using templates with *ops*_E–B_ swapped sequences or Y_E_–Y_B_ hybrid proteins that separate potential NGN vs. KOW contributions (Fig. 5a).

**Figure 5.**
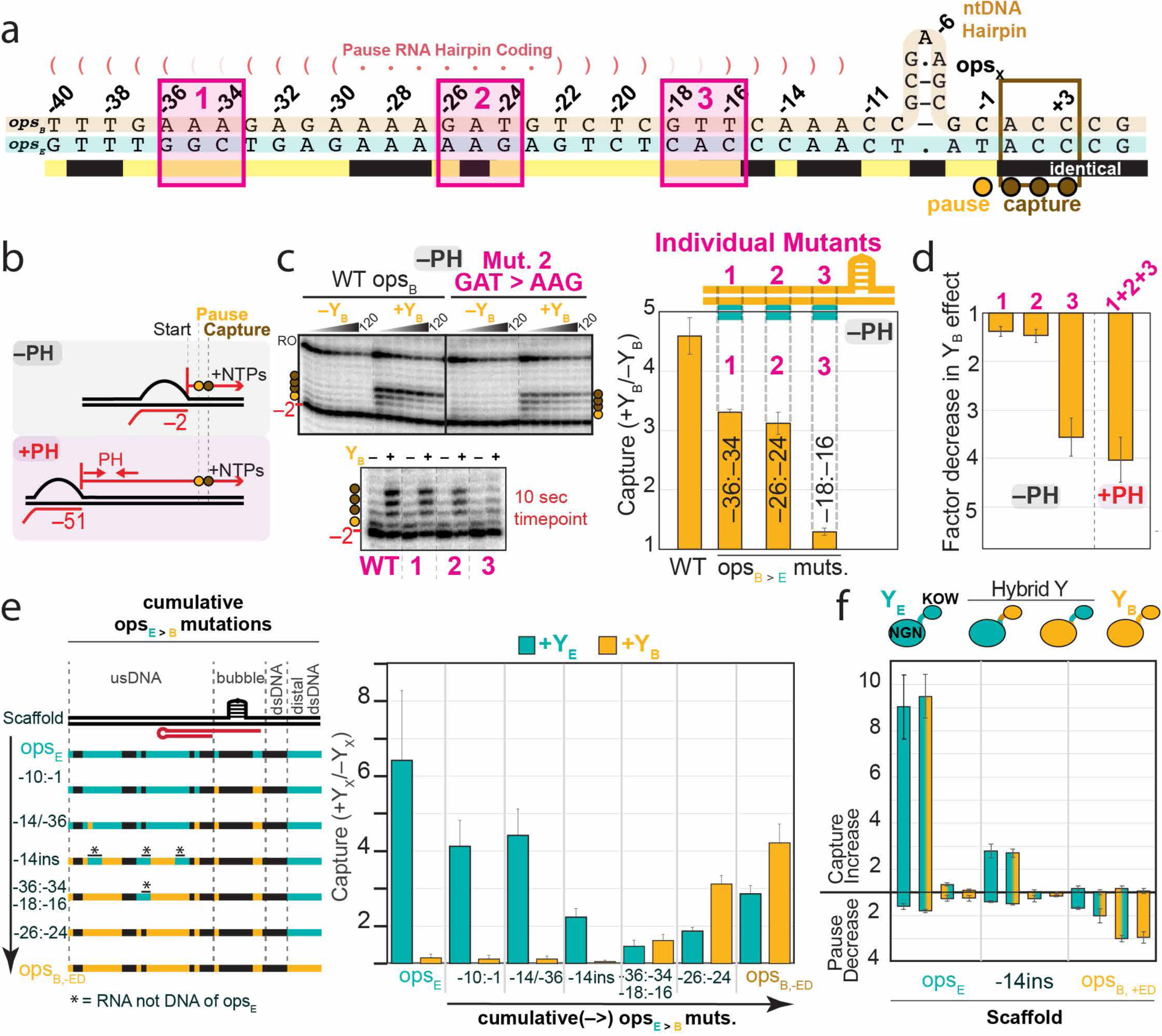
Y_X_ distinguish *ops_X_* binding sites through variations in the non-template DNA and upstream duplex DNA. **a)** Diagram comparing Class I *ops*_E_ and *ops*_B_ sequences. Regions varied in experiments shown in panels b-d are highlighted in magenta. **b)** Experimental scheme to assay effects of the usDNA and PH. **c)** PIVoT assays (1 µM NusA, 100 µM each NTP, 150 nM Y_X_ when added) comparing Y_B_ fold effects on capture on WT versus mutant scaffolds. The gel panels depict (top) time-courses of pausing on WT *ops_B_* vs a mutant and (bottom) a representative single time-point (10 s after NTP addition) comparison of Y_B_ effects across multiple mutant scaffolds. Error bars are SD from three replicates. **d)** Comparison of Y_B_ effects in the absence or presence of a PH. Error bars are SD from three replicates. **e)** Y_X_ fold effect on pausing at the 45 s time point on WT *ops*_E_ or hybrid sequences progressively mutated from *ops*_E_ towards *ops*_B_ with the escape duplex disrupted (see Supplementary Fig. 3 for scaffold sequence). PIVoT assays were performed in at least triplicate at a single timepoint (45s). Error bars are SD from ≥3 replicates. **f)** PIVoT assay of pause and capture for WT vs Hybrid NGN–KOW Y_X_ (150 nM each) on WT vs hybrid *ops_X_* sequences. Error bars are SD from three replicates.

To ask if Y_X_ recognizes the upstream DNA or the PH RNA encoded by it, we first tested whether the PH-encoding DNA sequences affected Y_X_ action in the absence of a PH (Fig. 5b). With the PH removed, Y_B_ stimulated RNAP capture at the *ops*_B_ pause site by a factor of ∼4.5 (Fig 5c). When 3-bp segments of the *ops*_B_ usDNA were replaced with *ops*_E_ sequence, Y_B_ capture of RNAP decreased either modestly (substitutions 1 and 2) or nearly completely (substitution 3). However, when we assayed Y_B_ capture of RNAP on a +PH scaffold, we observed a Y_B_ effect indistinguishable from the effect of substitution 3 alone on the –PH template (Fig. 5d). We conclude that Y_B_ recognition of the extended *ops*_B_ site depends on the usDNA and not on the PH RNA.

We next investigated the contributions of the upstream sequences in progressively interconverted *ops*_E_ and *ops*_B_ to PEC capture by Y_X_ (Fig. 5e, Supplementary Fig. 3). To simplify this comparison, we used a variant of *ops_B_* in which capture was activated by removing the ED (Supplementary Fig. 2, 3). Strikingly, Y_E_ continued to function even when the *ops*_E_ ntDNAhp was changed to the *ops*_B_ ntDNAhp. However, the Y_E_ effect was mostly lost and Y_B_ capture progressively increased as the usDNA was increasingly converted to *ops*_B_ sequence (Fig. 5e). Thus, multiple segments of usDNA contribute to Y_B_ recognition of *ops*_B_. Consistent with our in vivo experiments (Fig. 2f), we conclude that *ops*_X_ sequences are multipartite ntDNA and usDNA signals of ∼40 nucleotides whose constituent parts variably contribute to Y_X_ recruitment in different CPS operons.

We next asked if the NGN alone recognizes *ops*_X_ as it does for RfaH–*ops* interaction^23,59^ or if the KOW domain might also participate, as proposed for LoaP^54^. Attempts to purify a Class-1 NGN alone yielded only insoluble protein. Instead, we compared NGN–KOW Y_E–B_ hybrids to Y_E_ and Y_B_ on *ops*_E_, *ops*_B_, and an *ops*_E-B_ hybrid scaffold (Fig. 5f). For both Y_E_ and Y_B_, the effect on capture or pausing was determined completely by the NGN domains. We conclude that recognition of *ops*_X_ by at least Class-1 Y_X_ is mediated by the NGN and not the KOW domain.

### Class-1 Y_X_ protects upstream DNA from exonucleolytic cleavage

For the Y_X_ NGN to contact upstream duplex DNA, the DNA must distort from a canonical B-form trajectory departing the PEC (Fig. 6a). Although protein interactions can easily bend duplex DNA^60^, we sought direct physical evidence for usDNA–Y_X_-NGN interaction. Exonuclease III (ExoIII) has been used extensively to detect PEC boundaries on DNA^61–63^. Since Y_E,B_ variably depend on distal usDNA in our activity assays, we assayed *ops*_E,B_ with cognate Y_X_.

**Figure 6.**
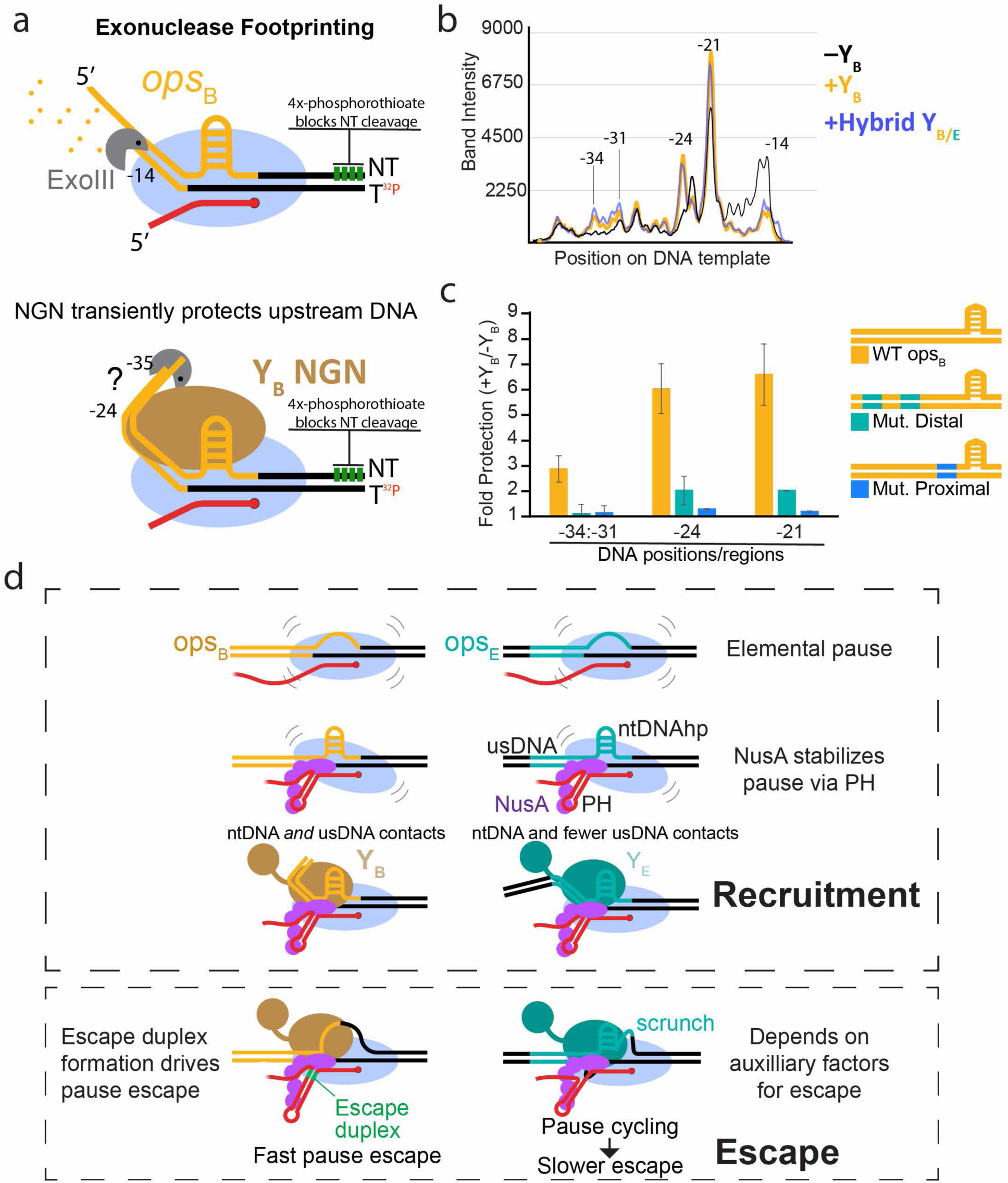
Y_X_ contacts sequences in the upstream duplex DNA and ntDNAhp to recognize cognate *ops*_X_ via a capture-then-escape mechanism. **a)** Y_X_ protects the distal upstream DNA from exonucleolytic cleavage. Exonuclease III (ExoIII) cleaves in the 3′-to-5′ direction but temporarily halts when encountering obstacles such as DNA-bound proteins. Protection was assayed in the absence or presence of Y_X_ or hybrid NGN-KOW at various timepoints. b) Pseudodensitometry traces of template DNA cleavage products separated by 8% Urea-PAGE. Band intensities reflect relative levels of cleavage products after 5 seconds of exonucleolytic cleavage. Traces are representative of experiments performed in at least duplicate (Extended Data Fig. 9). **c)** Quantification of upstream DNA protection from exonucleolytic cleavage at various regions on WT or mutant *ops*_B_ scaffolds (n=2; error bars are the range of duplicates). **d)** Model for Y_X_-specific recruitment. *Bfr*RNAP pauses in the 5′ leader of CPS operons to provide time for Y_X_ recruitment. These pauses arise initially through RNA–DNA contacts to RNAP (elemental pause), then are stabilized synergistically by a PH and NusA. Y_X_ is recruited with high fidelity to cognate operons by multipartite ∼40-bp *ops*_X_ elements, with variable influence of constituent elements depending on the CPS operon. Y_X_s that interact extensively with cognate *ops*_X_ (Y_A_, Y_B_) are associated with novel escape duplex–encoding PHs, which provides force in the form of base-pairing to drive forward translocation and inhibit backtracking. Differences among escape mechanisms (e.g., those with or without EDs) may aid differential regulation.

Over the full time course, Y_E,B_ strongly stabilized a –21 footprint, 6-7 base pairs upstream of RNAP (Extended Data Fig. 9). However, Y_B_ but not Y_E_ also slowed ExoIII digestion at –24, and –31 to –34. Further, these same upstream protections were caused by a Y_B,E_ NGN–KOW hybrid (Fig 6b, Extended Data Fig. 9). We conclude that Y_B_ NGN likely contacts usDNA at least near –21 to –24, and –31 to –34.

As an additional test of the upstream Y_B_ contacts, we performed ExoIII assays on scaffolds containing *ops*_B_ to *ops*_E_ sequence changes to distal usDNA (–36 to –34 and –26 to –24) and proximal usDNA (–18 to –16). These substitutions strongly reduced upstream protection from ExoIII (Fig. 6c, Supplementary Fig. 4). Together, our results suggest a set of Y_X_ specificity determinants reflected in both physical contacts detected with ExoIII and sequence effects on Y_X_ activity.

To understand these contacts in a structural context, we modeled Y_E_ and Y_B_ into an RfaH-*ops*-PEC structure (PDB 8PHK)^50^. Both Y_E_ and Y_B_ are predicted to have a much larger positively charged surface approximately in the path of the usDNA (Extended Data Fig. 10). This charge is created largely by basic residues in the beta hairpin mini-domain of Y_E,B_ and could position the usDNA for sequence-specific readout by NGN.

## DISCUSSION

Human gut *Bacteroides* strains synthesize numerous surface capsular polysaccharides that are highly regulated to create subpopulations in which primarily a single PS locus is transcribed, providing phenotypic plasticity to environmental challenges. To coordinate CPS gene expression in a manner that maximizes CPS diversity, *Bacteroides* have developed a complex hierarchy involving locus-specific cognate Y_X_ activation and noncognate Z_X_ inhibition.

We have elucidated the biochemical mechanisms of *Bacteroides* CPS hierarchical control (Fig 6d): (*i*) *Bfr*RNAP pauses prominently at single CPS leader-region pause sites (*ops*_X_); (*ii*) *ops*_X_ programs NusA-enhanced, RNA hairpin-stabilized transcriptional pauses that create time windows for Y_X_ recruitment; (*iii*) Z_X_ inhibits non-cognate Y_X_ directly via differential binding affinities, forming a heterodimer that precludes Y_X_ recruitment by steric clash of Z_X_ with RNAP and *ops*_X_; (*iv*) Y_X_ locus–specific recruitment depends on multipartite interactions of the Y_X_ NGN domain with the exposed *ops*_X_ ntDNA and upstream duplex DNA; (*v*) Y_X_s evolved into functionally distinct classes; and (*vi*) Y_X_-bound PECs use different mechanisms to escape *ops*_X_. This combination of multiple functions at a single pause site has little precedent and may reflect the strong evolutionary pressure associated with the challenges of discriminating among multiple similar NusG_SP_s.

*Bacteroides* belong to the greater phylum Bacteroidota, evolutionarily distant from the commonly studied model organisms *E. coli* (Pseudomonadota) and *B. subtilis* (Bacillota). Despite the importance of these bacteria to human health, there is a limited understanding of *Bacteroides* transcription regulation. Our recombinant *Bfr*RNAP overexpression system enables facile production and genetic manipulation of *Bfr*RNAP. Multiple questions can now be addressed, including the roles of novel RNAP sequence insertions^64^, the molecular interactions of RNAP with TFs (e.g., σ^A^) and small molecules (e.g., ppGpp), and sequence-dependent effects on transcriptional activities (e.g., backtracking, translocation, etc.). Recombinant RNAPs enable studies of both lineage-specific transcription mechanisms and evolutionary comparisons. r*Bfr*RNAP will enhance mechanistic understanding in the entire field of transcription, as demonstrated by numerous recent studies in *M. tuberculosis*, *C. difficile*, and *B. subtilis*^26–28,65,66^.

We found that *ops*_X_ recruitment sites for Y_X_ are ∼40 bp multipartite DNA elements with both upstream duplex and transcription bubble ntDNA components, in striking contrast to the 12-nucleotide ntDNAhp (*ops*) necessary for RfaH-recruitment and the proposed nascent RNA hairpin necessary for LoaP recruitment^54,59^. The ntDNAhps between *ops* and *ops_X_* differ in apparent structure and position relative to the pause site. All eight *ops* sites in *E. coli* targeted by the single RfaH encode the same ntDNAhp sequence: 5′-<u>GCG</u>GT<u>AGC</u>^67^, having conserved and variable elements compared to the longer *Bacteroides* 5′-<u>YGCG</u>N<u>AGCR</u> ntDNAhps. These key mechanistic differences highlight how *Bacteroides* evolved to manage numerous NusG_SP_. Extensive Y_X_–*ops_X_* interactions may also accelerate *Bacteroides* adaptation by expanding the sequence space available for functional bifurcation following gene duplication.

Z_X_ inhibits Y_X_ recruitment primarily by blocking Y_X_ interaction with the conserved ′ clamp helices (CH) and the *ops*_X_ usDNA. Z_X_ could also tune heterologous operon PSX expression or limit self-expression through negative feedback. Ultimately, Y_X_-Z_X_ interactions define the cell surface architecture of *Bacteroides.* Our findings provide a foundation for understanding them.

The closer start-codon proximity to *ops*_X_ (9 bp) suggests Class-2 Y_X_ may play a stronger role in ribosome association for coupled transcription–translation of the Y_X_ gene. Translation is not well studied in *Bacteroides*^68–71^, but both the similarity of anti-pausing by *Bfr*NusG to *Eco*NusG (Extended Data Fig. 4) and the location of stop codons relative to intrinsic terminators^72^ suggests transcription and translation may be coupled in *Bacteroides* – like *E. coli* but unlike *B. subtilis*^72–82^. RfaH is thought to recruit ribosomes for coupled translation in *E. coli*^83,84^. Start codon GUG is thought to initiate ribosomes 5–10 times more weakly than AUG in *E. coli*^85^. Taken together, these differences are consistent with evolution of Class-2 Y_X_-*ops*_X_ pairs for tight linkage of Y_X_ and ribosome recruitment at *ops_X_* sites immediately adjacent to the translation start site. Both these potential distinctions (relative to Class 1) in Class-2 Y_X_-*ops*_X_ function and interesting differences evident for Y_C_-*ops*_C_ and Y_G_-*ops*_G_ require future experimental investigation.

We also discovered a novel regulatory RNA element – the *ops_X_* PH escape duplex (ED) – involved in the regulation of PSA and PSB. The conserved role of PHs at *ops*_X_ is to enhance pausing with NusA. The ops_A,B_ ED provides a driving force to propel RNAP out of pause-cycling traps created by extensive interactions that occur at these sites. Possibly, *ops*_E_ does not encode an ED because Y_E_ interacts with less sequence (Extended Data Fig. 9) and Gre factor may be sufficient for its escape as it is for RfaH^50^. Alternatively, the strong kinetic difference in escape mechanisms could be exploited by *Bacteroides* in CPS expression control. We propose that the ED evolved in response to evolutionary pressure to expand Y_X_ specificity.

Our results provide new mechanistic insights into transcriptional regulation by a large class of NusG_SP_, Y_X_ (UpxY). We find that determinants of transcriptional pausing in the phylum Bacteroidota resemble those found for other bacteria, but that recruitment sites for these NusG_SP_s differ notably both in being multipartite and much more extensive (∼40 bp) than found for *E. coli* RfaH (∼12 bp). Two novel aspects of the Y_X_ recruitment mechanisms provide precedent for new types of transcriptional regulation: (1) the upstream DNA is a sequence-specific platform for PEC regulation, and (2) pause hairpins can include escape duplexes that can drive escape from regulator-stabilized pauses. These discoveries highlight the importance of studying transcriptional mechanisms in diverse bacteria.

## METHODS

Plasmids, oligonucleotides, and strains used in this study are listed in Supplementary Tables S1-4. Nucleic acid scaffolds used in PIVoT assays are organized by figure in Supplementary Notes.

### *E. coli* strain construction

*E. coli* strain RL3569 was created by P1 transduction of RL1674 with donor strain RL3570^86^ harboring the rifampicin-resistance mutation S522F in *rpoB*. Briefly, 5 mL of donor strain RL3570 was grown to saturation (overnight) in LB + 5 mM CaCl_2_. The next day, 50 µl of the donor strain was mixed with 100 µl of a 10^−5^ dilution (in LB + 5 mM CaCl_2_) of a freshly made P1 stock, then incubated at 37 °C for 20 minutes without shaking. 2.5 mL of 45c-equilibrated R top agar (0.8 % agar, 1% tryptone, 0.8% NaCl, 0.1% yeast extract, supplementing to a final concentration of 2 mM CaCl_2_ and 0.1% glucose after autoclaving) was added to the bacteria-phage mixture, flicked to mix, then poured evenly onto a thick, moist, freshly-made R plate (1.2% agar, 1% tryptone, 0.8% NaCl, and 0.1% yeast extract, supplementing to a final concentration 2 mM CaCl_2_ and 0.2% glucose after autoclaving). The plates were incubated at 37 °C overnight in a plastic bag with wet paper towels. The next day, the plate was transferred to a 4c room and overlayed with 5 mL of MC solution (10 mM MgSO_4_ + 5 mM CaCl_2_). After a 5 hour incubation at 4 °C, the overlayed solution containing fresh P1 lysate was collected, 0.2 µm filter-sterilized, then stored in the dark at 4c until use. The recipient strain (RL1674) was grown to saturation (overnight) in LB + 5 mM CaCl_2_ + 20 µg chloramphenicol/mL. The next day, 100 µl of donor P1 phage serial dilutions were separately mixed with 100 µl of recipient strain overnight culture, then incubated at 37 °C for twenty minutes with no shaking. The mixture was plated on LB agar + 20 µg chloramphenicol/mL + 100 µg rifampicin/mL. Candidates were sequence-verified.

### *B. fragilis* strain construction

#### Bacterial growth

*B. fragilis* NCTC 9343 (ATCC25285; Genbank assembly ASM2598v1) strains were grown in basal medium^87^ or on BHI plates supplemented with 5 mg hemin/liter and 2.5 µg vitamin K_1_/L. Mutants Δ*mpi*M44 ^17^, Δ*mpi*M44Δ*upaZ* ^13^ and ΩPSE ^39^ were previously constructed. For selection of cointegrants, gentamycin (200µg/ml) and erythromycin (5 µg/ml) were added to the plates when indicated.

#### Construction of mutant PSE *ops* and HP-*ops* regions in 9343Δ*mpi*M44

Two different alterations to the PSE 5’ UTR were made in the Δ*mpi*M44 strain. In the first mutant, the *ops* sequence of the PSE locus (CTGCGAAGCATA) was replaced with the *ops* sequence of the PSA locus (ccgcgtagcgca). In the second mutant, a larger replacement was made and included the hairpin region adjacent to the *ops* sequence. The sequence from the PSE 5’ UTR (ttggctgagaaaaagagtctcacccaaCTGCGAAGCATA) was replaced with the sequence from the PSA 5’UTR (cggtttgaatgggaaaagatgtctcgtccaaaccgcgtagcgca). The recombinant plasmids were created by PCR amplifying two (*ops*) or three (HP-*ops*) DNA segments using Phusion polymerase (NEB) with Δ*mpi*M44 as template with the primers listed in Table S2. These segments were cloned into BamHI-digested pLGB13 ^88^ using NEBuilder (NEB). Plasmids were sequenced to confirm the correct assembly of the segments. Plasmids were conjugally transferred from *E. coli* S17 λpir to Δ*mpi*M44 and after overnight co-incubation, were plated on BHIS with gentamycin and erythromycin. The resulting cointegrants were passaged in basal medium for several hours and plated on BHIS with 50 ng anhydrotetracycline to select for double cross-over recombinants. These strains were tested by PCR for replacement of the PSE sequences with the respective PSA sequences and the genomes of these two strains were sequenced to confirm the correct replacements.

### Western immunoblot analysis

Bacterial strains were grown overnight to an apparent OD_600_ of ∼1.2. Bacteria were pelleted and resuspended in 1X LDS loading buffer (Invitrogen) and boiled for 5 minutes. Cell lysates (equivalent to 3.5 µl of the original culture) were loaded onto 4-12% NuPAGE (Invitrogen) and run with MES buffer until the 17 kDa molecular weight standard had run to the bottom of the gel to allow for migration of the high molecular weight PSE further into the gel. The contents of the gel were transferred to PVDF and blocked with 5% skim milk in TBS with 0.5% tween (TBST). The blot was probed with a mouse monoclonal antibody specific to PSE, washed with TBST, and probed with alkaline phosphatase conjugated goat-anti mouse IgG (Pierce). After washing with TBST, the blot was developed with BCIP/NBT (KPL).

### NET-seq

*B. fragilis* NCTC 9343 rpoC-3xFLAG was streaked onto BHIS plates and incubated at 37 °C anaerobically for 2 days. A swab from a dense area on the plate was used to inoculate overnight cultures. The next day, 10 mL of the overnight culture was used to inoculate 500 mL SBM (starting apparent OD_600_ 0.04 as measured by a Denville® CO8000 Personal Cell Density Meter). When the apparent OD_600_ measured 0.65, cultures were removed from the anaerobic chamber and 300 mL was used for subsequent steps.

To harvest nascent transcripts for the NET-seq workflow, cultures were filtered between two vacuum filtration systems using a 0.45 µm pore nitrocellulose filter (GVS Micron Sep, 1215305). Cells were scraped off each filter using a spatula and plunged immediately into liquid nitrogen (i.e., cells from the same culture were combined into the same 50mL conical tube containing ∼25 mL liquid nitrogen). Collected cells were cryo-lysed using a RETSCH mixer mill (MM 400) as previously described^35^, with the exception that 50mL stainless steel canisters and a 25mm stainless steel ball were used to perform the cryomilling.

To isolate nascent transcripts, we performed a modified 3xFLAG-IP protocol with previously described buffers^35^. Specifically, the thawed grindate volume was scaled to 5.5 mL with lysis buffer (1x lysis stock [20mM Tris, pH 8.0, 0.4% Triton X-100, and 0.1% NP-40 substitute], 100mM NH4Cl, 1x EDTA-free cOmplete Mini protease inhibitor cocktail [Roche Diagnostics GmbH, 11836170001], 10mM MnCl2, and 50U/mL RNasin [Promega, N211B], and 0.4 mg/mL puromycin), DNA was partially digested for 20 minutes with RQ1 DNase (0.054 U/mL [0.02 U/mL for the *E. coli*-only NET-seq pilot experiment])[Promega, M6101], and digestion reactions were stopped by addition of EDTA to 28mM (final concentration). RNAP-nascent transcript complexes were directly immunoprecipitated using Anti-FLAG M2 affinity gel (Sigma, A2220) (i.e., without buffer exchange), and the precipitated RNAP-nascent transcript complexes were subsequently washed four times (1x lysis stock, 100mM NH4Cl, 300mM KCl, 1mM EDTA, and 50U/mL RNasin)[Promega, N2515]. RNAP-nascent transcript complexes were eluted twice with 3xFLAG peptide (Sigma, F4799) (1x lysis stock, 100mM NH4Cl, 2mg/mL 3xFLAG peptide, 1mM EDTA, and 50U/mL RNasin). Nascent transcripts were purified using a miRNeasy kit [Qiagen, 217084] as previously described (Larson REF). However, to reduce phenol and chaotropic salt contamination, nascent transcripts were subjected to an additional overnight isopropanol-GlycoBlue (Invitrogen, AM9516) precipitation at −20 °C.

For nascent transcript library generation, we followed a modification of a previous NET-seq workflow^34,35^. Specifically, our workflow included using custom adaptors compatible with an Illumina NovaSeq X instrument. Likewise, the DNA adapter used for nascent transcript 3’ end ligation was adenylated using components from a NEB 5′ DNA Adenylation kit (E2610; 6µM DNA linker [RL15032], 80µM ATP, 6 µM Mth RNA ligase, and 1X Adenylation Reaction Buffer). The adenylation reaction was incubated for 4 hrs incubation at 65°C, inactivated at 85°C for 5mins, and precipitated overnight at −20°C with isopropanol and GlycoBlue. The precipitated, adenylated DNA linker was ligated to 750 ng of precipitated nascent transcripts, in duplicate, using components of a NEB T4 RNA Ligase 2, truncated (T4 Rnl2tr) kit (M0242; 10% DMSO, 22% PEG8000, 3 µM adenylated DNA linker, T4 Rnl2tr [14.7U/µL], RNasin [2U/µL], and 1x T4 RNA Ligase Reaction Buffer). These ligation reactions were incubated at 37 °C for 4 h. After this incubation, T4 Rnl2tr was inactivated by incubation with Proteinase K (0.04U/µL) (NEB, P8107) at 37 °C for 1 h. RNAs were fragmented, resolved, gel extracted, and precipitated as previously described^34,35^, with the exception that the gel extraction incubation at 70 °C was increased to 25 min. cDNAs were synthesized using a custom adapter (RL14637) and a previously described protocol^34,35^, with the exception that the reaction time was increased to 1 hr. Circularization of gel extracted and precipitated cDNAs was performed using a protocol previously described ^34,35^, with the exception that the circularization reaction incubation period was increased to 3 h and the gel extraction incubation period was increased as above. After circularization, cDNA libraries were PCR amplified using minimal cycles and custom adapters, gel extracted, and precipitated as previously described^34,35^. Library concentration and amplified product size distribution were determined using an Agilent TapeStation 4150. NET-seq libraries were sequenced by the University of Wisconsin-Madison Biotechnology Center on an Illumina NovaSeq X instrument.NET-seq data were processed using a combination of custom scripts and standard tools. Briefly, adapters, linker, and control oligos potentially contaminating each sample were trimmed from raw reads using cutadapt (v3.4). Reads with a minimum length of 14 nts were mapped to the *B. fragilis* genome (NC_003228.3) using Bowtie (v1.3.0) allowing both one mismatch and random assignment of reads mapping to multiple loci based on alignment stratum (Bowtie options --best -a -M 1 -v 1). Alignments were converted to BAM and BED files using samtools (v1.16.1) and bedtools (v2.30.0). The specific 3’ end counts for each genome position were determined using bedtools (options -d -strand - -5 [plus strand] or -d -strand + -5 [minus strand]).

### r*Bfr*RNAP cloning and purification

*B. fragilis* RNAP coding regions were codon-optimized using Gene Designer from DNA2.0 (now ATUM) using *E. coli* codon frequencies^89^ and amplified from synthetic DNA (IDT) of *B. fragilis* NCTC 9343, then cloned into a pRM756 backbone^90^, incorporating a His10-ppx tag at the C-terminus of ′ and a Strep tag at the N-terminus of β. RBS sites were optimized using denovodna.com^91,92^. This plasmid enables T7 overexpression of all subunits under IPTG control.

r*Bfr*RNAP was purified essentially as described previously for *E. coli* RNAP^93^, with some changes. Following transformation of RL3569 with pJS015, a colony was picked and inoculated into a 3 mL LB + 25 µg kanamycin/mL + 20 µg chloramphenicol/mL. 2 mL of overnight culture was used to inoculate 2 L LB + 25 µg kanamycin/ml + 10 drops Sigma Antifoam Y-30 Emulsion in baffled Fernbach flasks and incubated at 37 °C. When the apparent OD_600_ reached 0.4, the temperature was dropped to 16 °C, overexpression was induced by addition of 200 µM IPTG, and incubation was continued with shaking at 200 RPM overnight (∼18 h). Cell cultures were placed on ice for 20 min, then pelleted by centrifugation at 3000 x *g* for 15 min at 4 °C.

Moving forward, all steps were performed at 4 °C or on ice, and all buffers were filtered through 0.2 µm filters. Pellets were resuspended in 30 mL lysis buffer (50 mM Tris-HCl pH 8.0, 5% glycerol, 100 mM NaCl, 2 mM EDTA, 10 mM BME, 10 mM DTT, 0.1 mg/mL phenylmethylsulfonyl fluoride (PMSF), with one dissolved tablet of Roche cOmplete^TM^ ULTRA EDTA-Free Protease Inhibitor Cocktail). The resuspended cell solution was sonicated for 20 min total (alternating sonication on/off times of 5 min) with settings Power 8, Duty Cycle 20%. The lysate was then transferred to round-bottom polycarbonate tubes and spun at 27,000 x *g* for 15 min. The supernatant was transferred to a 100 mL beaker with stir bar, then 6.5% PEI was slowly added to a final concentration of 0.6% while stirring. The solution was stirred for one hour, then transferred to open-top, round-bottom polycarbonate tubes and spun at 11,000xg for 15 min. After decanting supernatant, a tissue homogenizer was used to resuspend the pellet in 25 mL of TGEDZ (10 mM Tris–HCl pH 8.0, 5 % glycerol, 0.1 mM EDTA, 5 µM ZnCl_2_, 1 mM dithiothreitol) with added 0.3 M NaCl. The solution was spun at 11,000 x *g* for 15 min. After decanting supernatant, a tissue homogenizer was used to resuspend the pellet in 25 mL of TGEDZ with added 1 M NaCl. The solution was spun at 11,000 x *g* for 15 min. The supernatant was transferred into a 100 mL beaker with stir bar, then finely-ground AmSO_4_ was added to the stirring solution to a final concentration of ∼0.37 g/mL and precipitated overnight. The solution was transferred to Oak Ridge round-bottom tubes and spun at 27,000 x *g* for 15 min.

The pellet was dissolved in 35 mL of HisTrap Binding Buffer (20 mM Tris-HCl pH 8.0, 500 mM NaCl, 5 mM imidazole, 5 mM beta-mercaptoethanol (BME), then spun at 27,000 x *g* for 15 min in the same Oak Ridge round-bottom tube. The supernatant was filtered through 0.2 µm filters and applied at 1 mL/min to a HisTrap HP 5 mL column, pre-equilibrated with HisTrap Binding Buffer. The column was washed with HisTrap Binding Buffer at 5 mL/min until A280 reached baseline, then washed at 5 mL/min with 2% HisTrap Elution Buffer (20 mM Tris-HCl pH 8.0, 500 mM NaCl, 1 M imidazole, 5 mM beta-mercaptoethanol [BME]) until A_280_ reached baseline. rBfRNAP was eluted at 5 mL/min with a 2-50% gradient of HisTrap Elution Buffer (translating to a 20-500 mM imidazole gradient). 3 mL elution fractions containing rBfRNAP were pooled, filtered through 0.2 µm filters, then the NaCl concentration was reduced to 150 mM for the following purification step by dilution with TGEDZ buffer.

HisTrap elution fractions were pooled then diluted with 100 mM Tris-HCl, pH 8.0, 1 mM EDTA, 10 mM DTT to adjust the salt concentration to 150 mM NaCl. The sample was then applied to a 5 mL Strep-Tactin® XT High Capacity column pre-equilibrated with 2 CV Buffer W (100 mM Tris-HCl, pH 8.0, 150 mM NaCl, 1 mM EDTA, 10 mM DTT) at 2 mL/min. The flow-through was reapplied to the column at 0.037 mL/min. The column was then washed with 5 CV of Buffer W. rBfRNAP was eluted with Buffer BXT (100 mM Tris-HCl, pH 8.0, 150 mM NaCl, 1 mM EDTA, 10 mM DTT, 50 mM D+Biotin (Acros Organics)).

Pooled fractions from the previous step were applied at 1.5 mL/min to a HiTrap HP column pre-equilibrated with TGEDZ + 200 mM NaCl. The column was then washed with TGEDZ + 200 mM NaCl until A280 reached baseline, then rBfRNAP was eluted with TGEDZ + 500 mM NaCl at 2.5 mL/min.

Pooled fractions from the previous step were dialyzed overnight in RNAP storage buffer (10 mM Tris-HCl, pH 8.0, 25% glycerol, 100 mM NaCl, 100 µM EDTA, 1 mM MgCl_2_, 20 µM ZnCl_2_, 10 mM DTT) using a 10 kDa MWCO cassette, then concentrated using Ultra-4, MWCO 100 kDa (Sigma-Aldrich Z648043-24EA) to a final concentration of 8 µM. The solution was then aliquoted, flash-frozen, and stored at –80 °C.

### Cloning and purification of transcription factors

All transcription factors (NusG, NusA, Y_A_, Y_B_, Y_C_, Y_E_, Y_F_, Y_H_, Y_B_(NGN)–Y_B_(KOW), Y_E_(NGN)– Y_B_(KOW)) were cloned into a pTYB2 backbone (Addgene catalog N6702S) after PCR amplification from *Bacteroides fragilis* ATCC 25285 (NCTC 9343) genomic DNA by NEB HiFi DNA assembly (Gibson Assembly). This vector enables IPTG-inducible over-expression of proteins fused at the C-terminus to the *Saccharomyces cerevisiae* VMA intein and chitin-binding domain. Importantly, to ensure efficient self-cleavage via the intein, an Ala residue was incorporated at the C-terminus of all transcription factor coding sequences.

After plasmid sequence verification, RL1674 (*E. coli* BL21 Rosetta^TM^ (DE3)) was transformed by electroporation with pTYB2-derived constructs, then plated on LB agar with 100 µg ampicillin/mL and 20 µg chloramphenicol/mL (for retention of pRARE2 plasmid). For each expression construct, a single colony was picked and used to inoculate a 3 mL overnight LB culture grown at 37°C containing the same concentration of antibiotics. The next day, 1 mL of overnight culture was used to inoculate a 200 mL LB culture containing antibiotics (3% ethanol was added for all Y_X_ constructs) and grown at 37°C. When the OD reached 0.2-0.3, the incubation temperature was dropped to 16 °C and shaking continued for 30 minutes. Subsequently, a final concentration of 200 µM IPTG was added and incubation continued overnight (16-18 hours). The next day, cultures were placed on ice for 20 min, then pelleted at 3000xg for 15 min at 4°C.

Pellets were resuspended in 40 mL of Chitin Wash Buffer (CWB; 30 mM Tris-HCl, pH 7.5-8.0 depending on protein pI, 0.5 M NaCl, 1 mM EDTA, 0.05% Tween^®^ 20) plus one dissolved tablet of Roche cOmplete^TM^ ULTRA EDTA-Free Protease Inhibitor Cocktail. The cell suspension was sonicated 10 min at 20% duty cycle, Power 8. The lysate was pelleted at 30,000xg for 30 min at 4°C, then the supernatant was passed through 0.2 µm filters.

The subsequent steps were performed at room temperature closely following manufacturer’s instructions. Briefly, 3 mL of a homogenous suspension of NEB Chitin Resin (Catalog S6651L) were loaded into a 25 mL Poly-Prep Gravity Chromatography Column (Biorad), washed with 5 mL of mQH_2_O, then equilibrated by washing 3 times each with 10 mL of CWB. The lysate was subsequently loaded onto the column, then washed three times each with 10 mL of CWB. Cleavage Buffer (CB) was made by adding 500 µl of 1 M DTT (prepared fresh from solid reagent) to 10 mL of CWB, then a quick flush was performed by adding 3 mL of CB. SDS-PAGE revealed no premature elution in the quick flush fraction. Immediately after dripping stopped, the bottom and top of the column were capped, parafilmed, and the column was incubated at room temperature overnight (16-18 hours) to allow sufficient time for cleavage. The next day, cleaved protein was eluted by addition of 1.5 mL CWB + 10 mM DTT, then dialyzed overnight in 10 mM Tris-HCl, pH 7.5-8.0 depending on pI, 2% glycerol, 100 mM NaCl, 100 µM EDTA, 10 mM DTT using a 10K MWCO cassette. After removal from the dialysis cassette, additional glycerol was added to a final concentration of 25%. The solution was aliquoted, flash-frozen, then stored at −80°C until use.

### PIVoT assays

A direct reconstitution approach was used to assemble elongation complexes (ECs). Briefly, RNA and template DNA oligonucleotides were mixed at a ratio of 1:1.2 (5 µM: 6 µM) in transcription buffer (TB; 20 mM Tris-OAc, pH 7.7, 40 mM KOAc, 5 mM Mg(OAc)_2_, 1 mM DTT), then annealed by slow cooling in a thermocycler. To assemble 10X ECs, first the annealed RNA:tDNA scaffold and RNAP were mixed in TB and incubated for 15 min at 37°C. Then, non-template DNA oligonucleotide was added and incubation continued for an additional 15 min at 37°C. The solution was diluted with TB to prepare 2X EC (subtracting volume of further additions) and incubated for 1 min at 37°C. Then, 5 µCi of [α-^32^P]NTP (depending on the scaffold) was added and incubated for 3 min at 37°C. Additional GTP was added such that the final concentration of GTP in the solution was 10 µM, and incubation continued for 3 min at 37°C.

2X ECs were aliquoted and all comparisons made were therefore performed with identically formed ECs. The assay was performed at 37°C: transcription was restarted by addition of 2X NTPs minus/plus transcription factors or storage buffer. For Fig 5c, Y_B_ was pre-incubated with halted ECs following reconstitution at –3 and incorporation labeling to –2 prior to restarting transcription. Timepoints were taken by mixing 5 µl reaction aliquots with 5 µl of 2X Stop Buffer (25 mM EDTA, 8 M Urea, 1X TBE, 0.1% bromophenol blue, 0.1% xylene cyanol). The ratio and concentrations of EC components in the 1X EC solution was 1:1.2:1.4:1.6 (R:T:RNAP:NT; 50 nM, 60 nM, 70 nM, 80 nM). The final reaction concentrations of transcription factors are indicated in each figure legend. Unless otherwise indicated, NTPs are added to a final reaction concentration of 100 µM. RNAs were resolved by 8% or 15% Urea– PAGE with 0.5X TBE running buffer until the leading dye ran off the gel. Gels were exposed to PhosphorImager screens and scanned using a Typhoon Phosphorimager. To quantify effects in ImageQuant, boxes were drawn around the pause band *ops*_X_, the capture band(s) (if applicable), and beyond. After subtracting background, the fractions of RNA at *ops*_X_ or at capture positions were averaged and errors reflect standard deviation from at least three replicates (unless indicated otherwise).

For the Z-titration assay in Figure 2d, data were fit in Kaleidagraph to a sigmoidal function of the form y = a+(b-a) / (1+(x/c)^d) where a= ymin, b is ymax, c is the Z_X_ concentration at mid-point, and d is slope at mid-point; and weighted by standard deviation (error bars) from three assays.

### Biolayer interferometry

Preparation of biotinylated-Y_E_: pJS060 was cloned similarly to other pTYB2-derived constructs (see above), with the exception that two oligos were included in the Gibson assembly to introduce the 16 codon Avi-tag^TM^ onto the N-terminus of upeY. Expression, cell harvesting, and lysis conditions are as described above. Avi-Y_E_ was biotinylated on a gravity column as described below:

The subsequent steps were performed at room temperature closely following NEB instructions. Briefly, 3 mL of a homogenous suspension of NEB Chitin Resin (Catalog S6651L) were loaded into a 25 mL Poly-Prep Gravity Chromatography Column (Biorad), washed with 5 mL of mQH_2_O, then equilibrated by washing 3 times each with 10 mL of CWB. The lysate was subsequently loaded onto the column, then washed three times each with 10 mL of CWB. The column was then washed with three times each with Avi Chitin Wash Buffer (AviCWB = 10 mM Tris 8.0, 0.5 M KGlu, 0.1% Tween20). Components from Avidity BirA500 Kit were used in the subsequent biotinylation reaction: a biotinylating solution (500 µL AviCWB, 70 µL of BiomixA, 70 µL Biomix B, 10 µL of 1 mg/mL BirA) was added to the column and the reaction was allowed to continue for 2.5 hours. The column was subsequently washed three times each with 10 mL of CWB. Cleavage Buffer (CB) was made by adding 500 µl of 1 M DTT (prepared fresh from solid reagent) to 10 mL of CWB, then a quick flush was performed by adding 3 mL of CB. Immediately after dripping stopped, the bottom then the top of the column were capped, parafilmed, and the column was incubated at room temperature overnight (16-18 hours) to allow sufficient time for cleavage. The next day, cleaved protein was eluted by addition of 1.5 mL CWB + 10 mM DTT, then dialyzed overnight in 10 mM Tris-HCl pH 7.5, 2% glycerol, 100 mM NaCl, 100 µM EDTA, 1 mM DTT using a 10K MWCO cassette. After removal from the dialysis cassette, additional glycerol was added to a final concentration of 20%. The solution was aliquoted, flash-frozen, then stored at −80°C until use. Importantly, Biotin-Y_E_ retained activity *in vitro*.

For each titration, 1 mL of 0.3 µM biotinylated-Y_E_ was prepared in Octet Binding Buffer 4.1 (OBB4.1 = PBS + 400 mM NaCl + 0.01% Triton X-100 + 0.25% BSA). Z_A_ solution was prepared at 100 nM in OBB4.1 with 2-fold serial dilutions down to 1.56 nM. Z_E_ solution was prepared at 500 nM in OBB4.1 with serial dilutions down to 31.3 nM. Plates were prepared for binding assays: in plate 1, 200 µL of OBB4.1 was placed in each well of column 1 containing a biosensor (up to 8 biosensors per experiment); plate 2 (containing ‘half-area’ wells permitting 100 µL volumes) column 1 contained 100 µL/well of OBB4.1, column 2 contained 100 µL/well of 0.3 µM biotinylated-Y_E_, and column 3 contained 100 µl/well of Z_X_ serial dilutions or buffer (as a blank/reference) prepared above.

A basic kinetics assay was performed using standard acquisition rates at 30c on a ForteBio Octet RED96 system. Octet® Streptavidin (SA) Biosensors were pre-equilibrated for 10 min at 30c. Step times: Baseline (Plate 2 Column 1 (P2C1)) = 60 sec; Loading (P2C2= 320 sec (or until 2 nm loading density reached); Baseline (P2C1) = 60 sec; Association (P2C3) = ≥300 sec; Dissociation (P2C1) = ≥ 300 sec.

Data were processed using Octet Data Analysis Software. The reference biosensor curve (bio-Y_E_ + buffer in place of Z_X_) was subtracted from all binding curves. Traces were subsequently aligned along the Y axis at pre-association baseline with interstep correction performed at the dissociation step. Noise Filtering (Savitsky-GolayFiltering, smoothingfunction) was performed. Data from each experiment were independently globally fit. For each binding pair tested, two out of three global fits have R^2^ values around 0.95 or greater and chi-squared values less than 3 as recommended by ForteBio. Given the two orders of magnitude difference in binding constants, limited conclusions we are making, and parsimonious agreement of these constants among replicates and with our PIVoT assays, we deemed the fits overall acceptable. The average and standard deviation of the kinetic parameters from the global fits are reported. Equilibrium constants are calculated from models. The value ‘Req/Rmax’ is reported as fraction Y_E_ bound.

### Exonuclease footprinting

Nucleic acid scaffolds used in exonuclease footprinting assays were each comprised of: i) a ^32^P-labeled template DNA oligo, ii) a non-template DNA oligo with four consecutive phosphorothioate bonds at the 3′ end, and iii) an RNA oligo with 3′ end at the position of pausing in *ops*_X_ and having noncomplementary bases upstream of the RNA-DNA hybrid to prohibit backtracking.

Template DNA oligo (20 μM) was labeled in a T4 PNK reaction with 1 μCi of [γ-^32^P]ATP and allowed to proceed for 15 minutes at 37 °C. ATP (1 μL of 1 mM) was subsequently added to the reaction and allowed to proceed for 30 minutes at 37°C. Reactions were stopped by heating at 65 °C for 20 min and oligos were subsequently purified using G-50 columns pre-equilibrated with TE and following the manufacturer’s instructions.

TECs were reconstituted essentially as described in *in vitro* transcription assays, except that the molar ratio of T:R:Pol:NT was 1:2:3:5 (50 nM ^32^P-T: 100 nM R: 150 nM RNAP: 250 nM NT). TECs were subsequently split into 35 μL aliquots and incubated with either storage buffer or Y_X_ variants for 3 minutes at 37 °C. Tubes were shifted to 30 °C and allowed to incubate for 3 minutes before removing a 5 μl aliquot (time 0) and mixing with equal volume 2X Stop Buffer. Exonuclease reactions were initiated by adding 100 U of exonuclease III, and aliquots were removed from reactions and mixed with stop buffer at times indicated in figures.

To quantify both transient and stable protection from exonucleolytic cleavage, pseudodensitometry traces were generated for the first timepoint lane. Regions of interest were identified by comparison to a sequencing ladder (Supplementary Fig. 4). Areas under the peaks of these regions were determined by manual integration in Microsoft Excel, then divided by the sum of the areas under all peaks to the right of it. These values were determined in the absence or presence of Y_B_, and their ratio is reported as fold change (+Y_B_/-Y_B_) for each sequence variant.

### Structural Models

A model of Y_B_ was made using Modeller^94^_,95_ and fitted to 8PHK^50^. Additional upstream and downstream DNA were modeled using Pymol. The Y_E_-Z_A_ complex structure was predicted using AlphaFold 3^53^, yielding an interface predicted template modeling (iPTM) score of 0.89 and predicted template modeling (pTM) score of 0.9 (values above 0.8 represent confident high-quality predictions). Additional confidence metrics are illustrated in Extended Data Fig. 14. RNA secondary structures were predicted using RNAFold^96^.

The *Bfr*RNA polymerase PEC model was generated using Modeller^94,95^, the *M. tuberculosis* PEC formed on the *B. subtilis trpL* pause sequence (8E74)^27^, NusA and NusG NGN models from Swiss model^97^, and *Porphymonas gingevalis* RNAP (8DKC)^98^.

## Acknowledgements

We thank members of the Landick and Comstock labs for helpful discussions and comments on the manuscript. This work was supported by NIH R01 GM038660 and USDA Hatch WIS05004 to R.L, NIH R01 AI093771 to L.C., the Duchossois Family Institute, and the DOE Office of Science, Biological and Environmental Research Program Great Lakes Bioenergy Research Center (DE-SC0018409). A.G. was supported by the NIH Predoctoral Training Program in Genetics (T32 GM007133). J.S. was supported by the NIH Biotechnology Training Grant (T32 GM135066 and T32 GM008349), an NIH F31 Graduate Fellowship (F31 GM142153), and a SciMed Graduate Research Scholars Fellowship from the UW–Madison Graduate School and Wisconsin Alumni Research Foundation.

## Author Contributions

R.L. and J.S. conceived of the study. J.S. conceived and developed assays, cloned most plasmids, purified all proteins, performed most experiments, and analyzed data. K.F. constructed plasmids for *B. fragilis* genetic manipulation, created *Bacteroides* strains and performed Western blots. M.E., B.M., and J.S. wrote custom scripts. J.S. and R.L. interpreted data. M.E., Y.P, and A.G. performed experiments. R.L. and J.S. constructed structural models. J.S. and R.L. wrote the original manuscript and designed figures. J.S., R.L., and L.C. revised the manuscript. R.L, L.C., and J.S., secured funding. R.L. and L.C. supervised the study.

## Abbreviations

NusG_SP_: specialized paralog of NusG
PSX: Capsular Polysaccharide Operon X (X = A–H)
Y_X_: UpxY
Z_X_: UpxZ
RNAP: RNA polymerase
r*Bfr*RNAP: recombinant *Bacteroides fragilis* RNA polymerase
PEC: Paused elongation complex
*Eco*RNAP: *E. coli* RNA polymerase
ntDNA: non-template DNA
tDNA: template DNA
usDNA: upstream DNA
*ops_X_*: operon polarity suppressor of PSX operon
asDNA: antisense DNA
asRNA: antisense RNA
ED: Escape duplex
PH: Pause hairpin
KOW: Kyprides Ouzounis Woese Domain
NGN: NusG-like N-terminal domain

---

**Extended Data Fig. 1.**
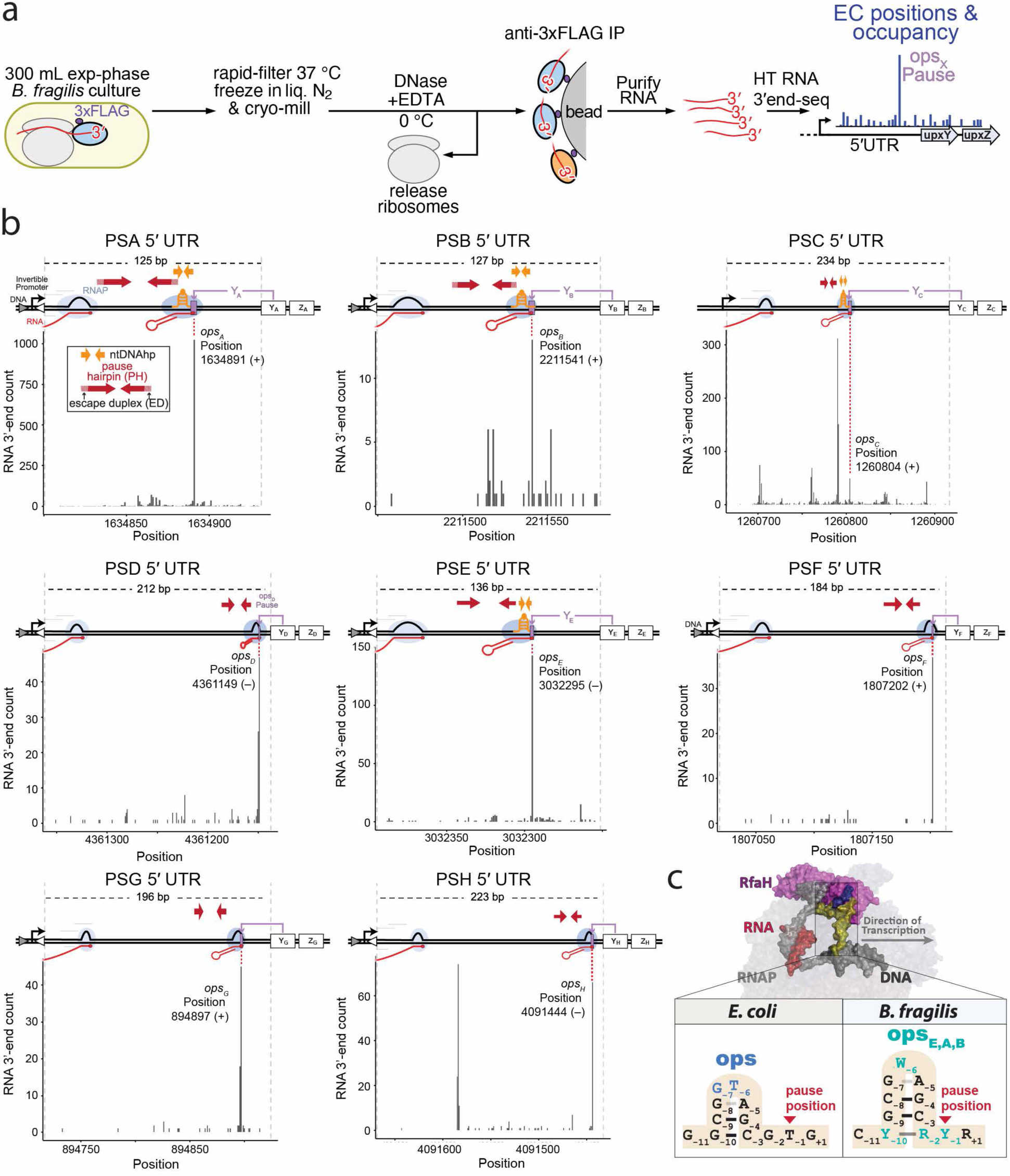
*ops*_X_ pauses found by NET-seq. **a)** Schematic depiction of NET-seq. **b)** CPS operon leaders aligned with mapped NET-seq reads (Genbank accession NC_003228.3). Genome coordinates are PSA 1634806:1634931(+); PSB 2211455:2211581 (+); PSC 1260676:1260910 (+); PSD 4361353:4361141(–); PSE 3032389:3032254(–); PSF 1807026:1807210(+); PSG 894725:894921(+); PSH 4091659:4091436(–). Red arrows indicate PH stems. Orange arrows indicate ntDNAhp stems. **(c)** Comparison of ntDNAhps in some CPS leader *ops*_X_ sites to the RfaH *ops* ntDNAhp^50^. Base lettering follows IUPAC nomenclature. Blue colored nucleotides in *ops* make base-specific contacts to RfaH^23,50,59^.

**Extended Data Fig. 2.**
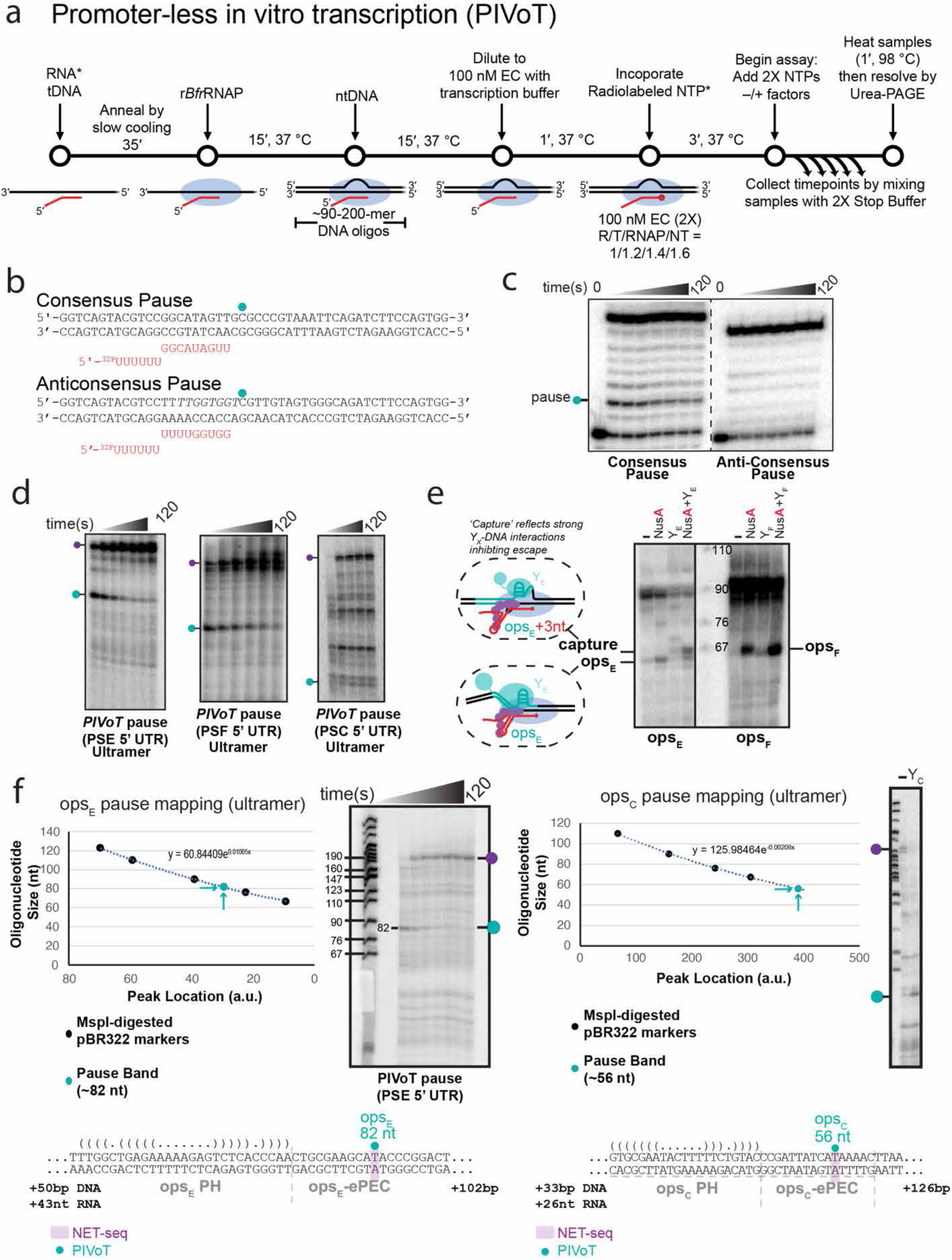
*ops_X_* pause mapping by PIVoT. **a)** Schematic of PIVoT assays. *, source radiolabel (either 5′-^32^P-labeled RNA or incorporation of [α-^32^P]NMP at RNA 3′ end, depending on the assay; see Methods). **b)** Scaffolds used in panel c to assay r*Bfr*RNAP pausing propensity. **c)** r*Bfr*RNAP pauses on consensus but not anti-consensus pause sequences. **d)** Representative transcriptional pauses from distinct CPS operon leader regions mapped in vitro. PIVoT assay of relevant regions from CPS operon leader regions. **e)** One of three replicates of effects of Y_X_ and NusA on pausing and capture assayed by PIVoT using a single timepoint (45 s or 8 min after NTP addition for Y_E_–*ops*_E_ and Y_F_–*ops*_F_, respectively) for results shown in Fig. 1F (see also Extended Data Fig. 3). Both *ops*_E_ and *ops*_F_ could be seen in proximity to 67 nt marker on short scaffolds. **f)** *ops*_E_ and *ops*_C_ pause RNAs mapped by quantitative comparison to markers of known sizes. Data generated by quantitation of pseudodensitometry traces drawn for assay and marker lanes. The *ops*_C_ pause band (15 s timepoint) is weak in the absence of Y_C_ (full time course in Extended Data Fig. 3, bottom gel), so Y_C_ was added in a separate assay and run alongside to aid pause RNA identification.

**Extended Data Fig. 3.**
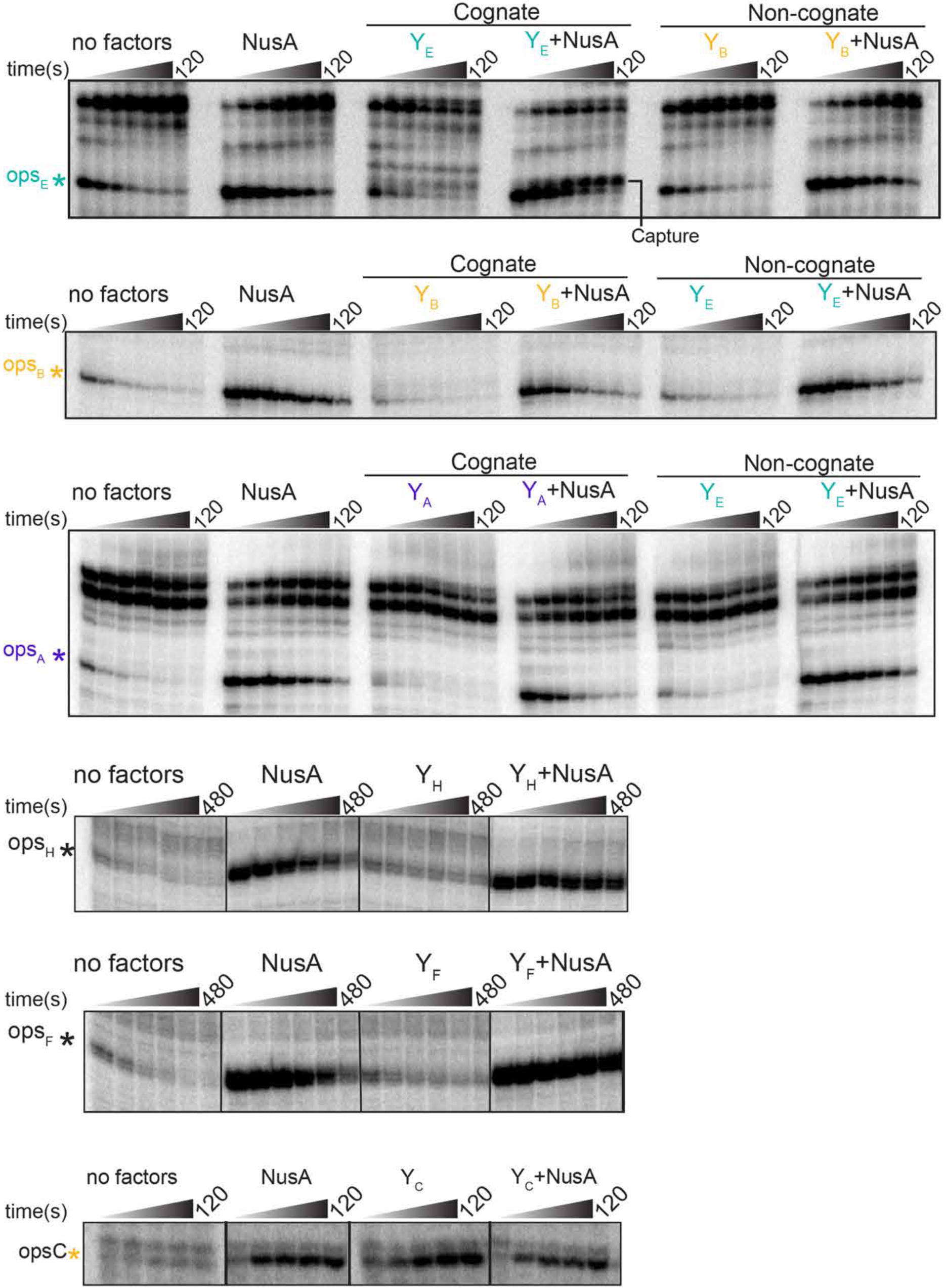
Y_X_ and NusA effects at 6 different *ops_X_* sites. PIVoT assays for *ops*_E_, *ops*_A_, and *ops*_B_ (150 nM Y_X_, 1 µM NusA, and 200 µM NTPs added concomitantly where indicated). PIVoT assays for *ops*_H_ and *ops*_F_ (1 µM Y_X_, 1 µM NusA, and 500 µM NTPs added concomitantly where indicated). PIVoT assay for *ops*_C_ (0.5 µM Y_C_, 0.5 µM NusA, and 200 µM NTPs added concomitantly where indicated).

**Extended Data Fig. 4.**
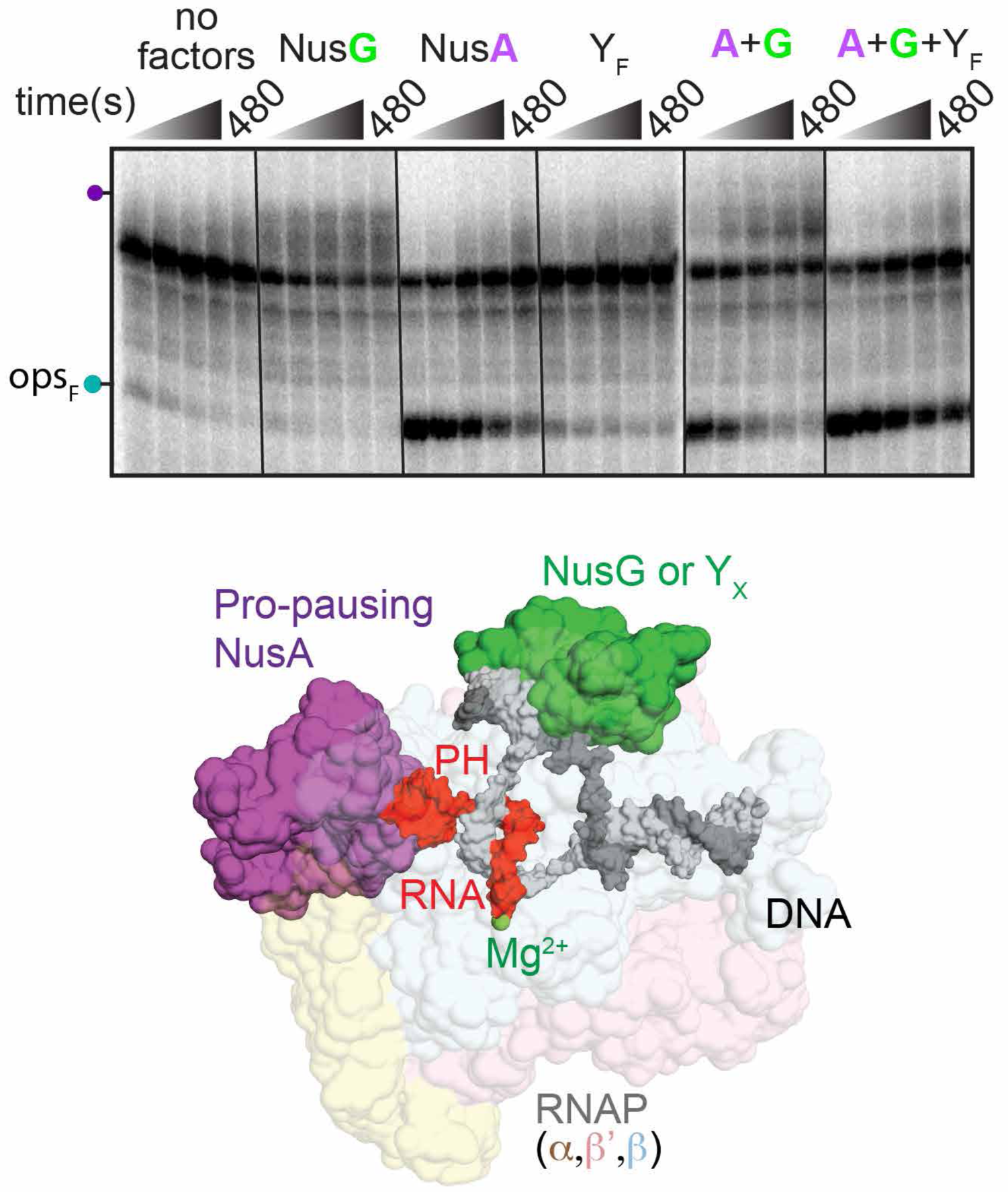
Y_F_ outcompetes NusG at *ops*_F_. PIVoT assays performed with 1 µM Y_F_, 1 µM NusG, 0.5 µM NusA, and 0.5 mM NTPs added concomitantly where indicated. Homology model of a *Bfr*RNAP PEC bound by NusG NGN, and NusA (see Methods).

**Extended Data Fig. 5.**
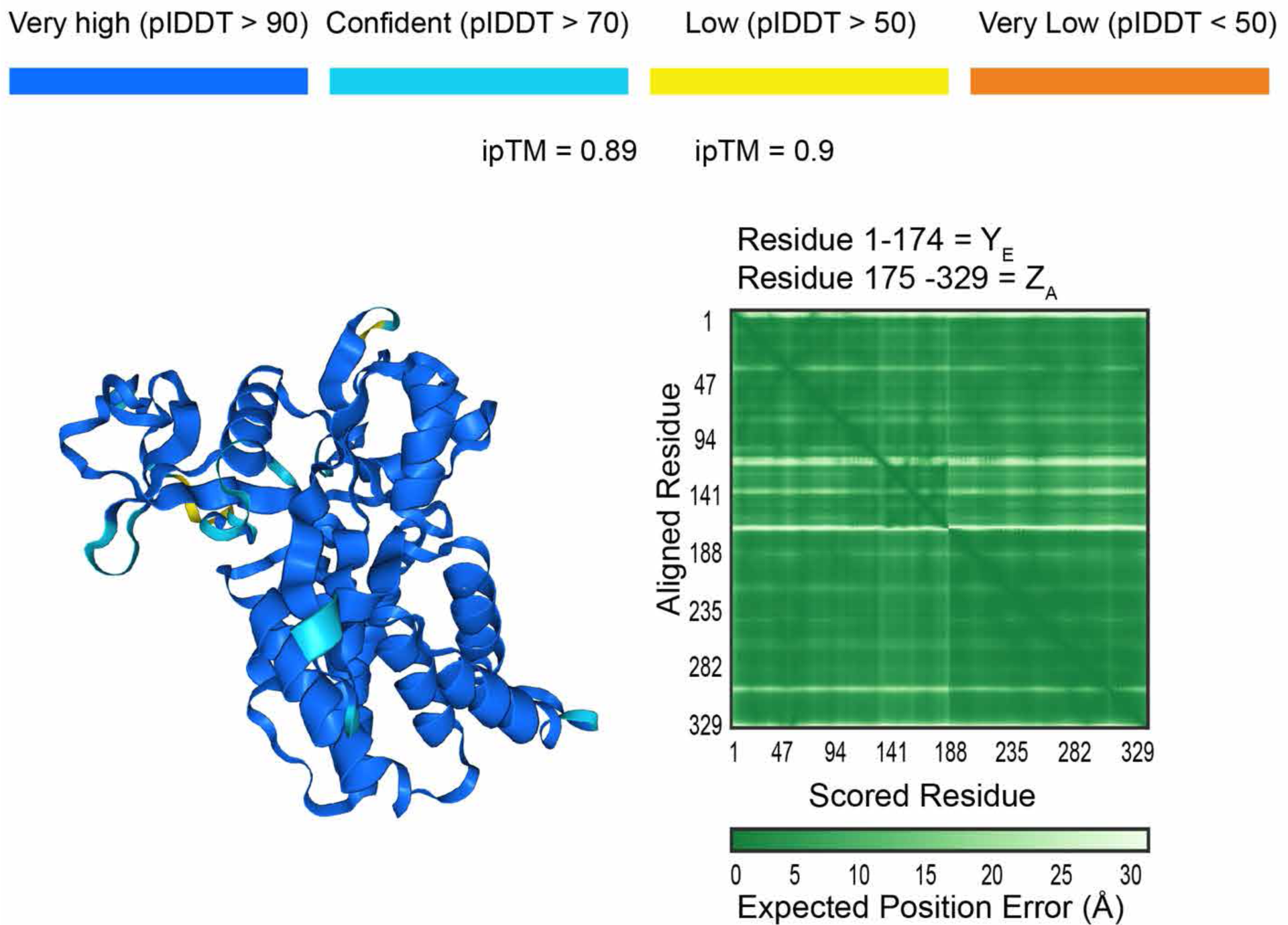
Confidence metrics from AlphaFold 3^53^ Y_E_-Z_A_ complex structural prediction. The interface predicted template modeling (iPTM) score of 0.89 and predicted template modeling (pTM) score of 0.9 represent confident high-quality predictions (values greater than 0.8).

**Extended Data Fig. 6.**
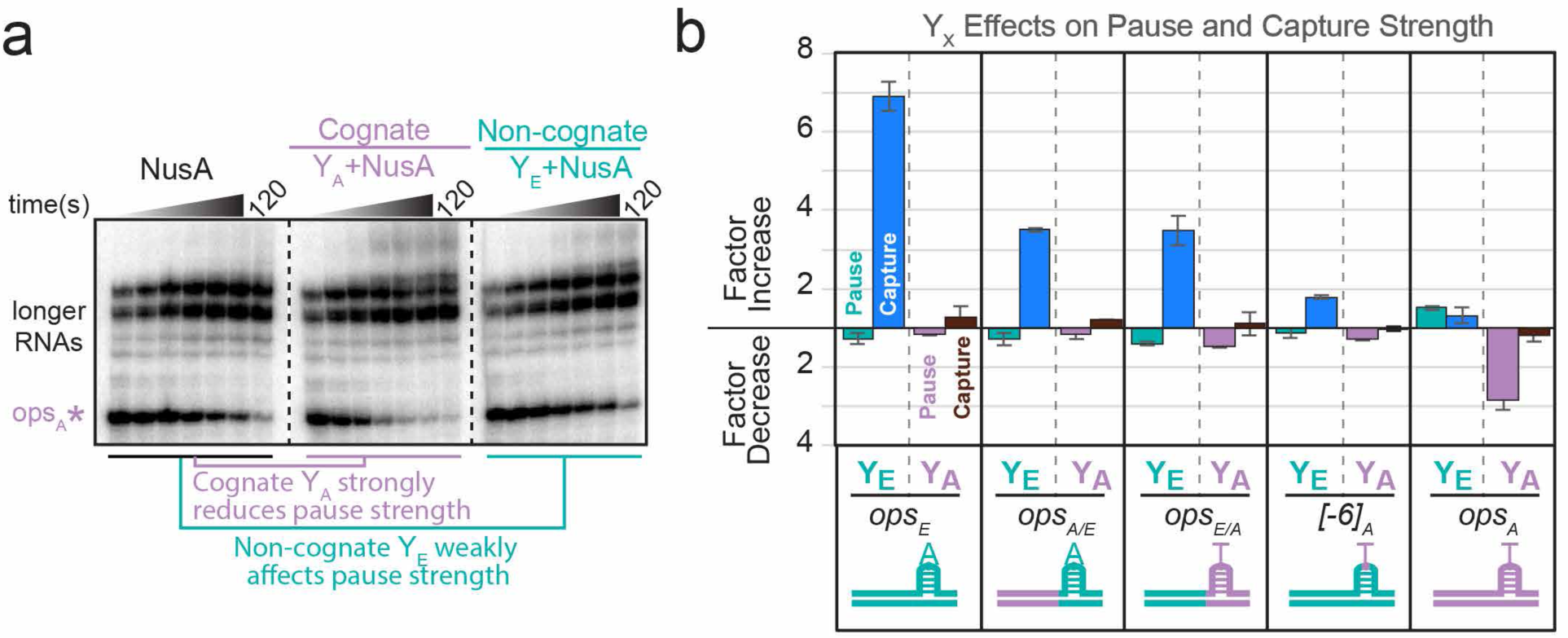
Effects of Y_E_ and Y_A_ on WT *ops*_X_ and hybrid *ops*_X_ sequences in PIVoT assays. **a)** Full time-course PIVoT assay illustrating cognate Y_A_, but not non-cognate Y_E_, modulates the strength of the *ops*_A_ pause. **b)** PIVoT assay quantitation of Y_A_ and Y_E_ activities on WT or hybrid scaffolds. Assays were performed in at least triplicate at a single timepoint (45 s) in the presence of 1 µM NusA and 100 µM NTPs, adding 150 nM Y_E_ or 150 nM Y_A_. Fold effects are relative to the NusA-only condition. Y_E_ association on WT *ops_E_* manifests primarily as capture activity, whereas Y_A_ association at *ops*_A_ manifests as anti-pausing activity (see Extended Data Fig. 3). Error bars are standard deviations from three experiments.

**Extended Data Fig. 7.**
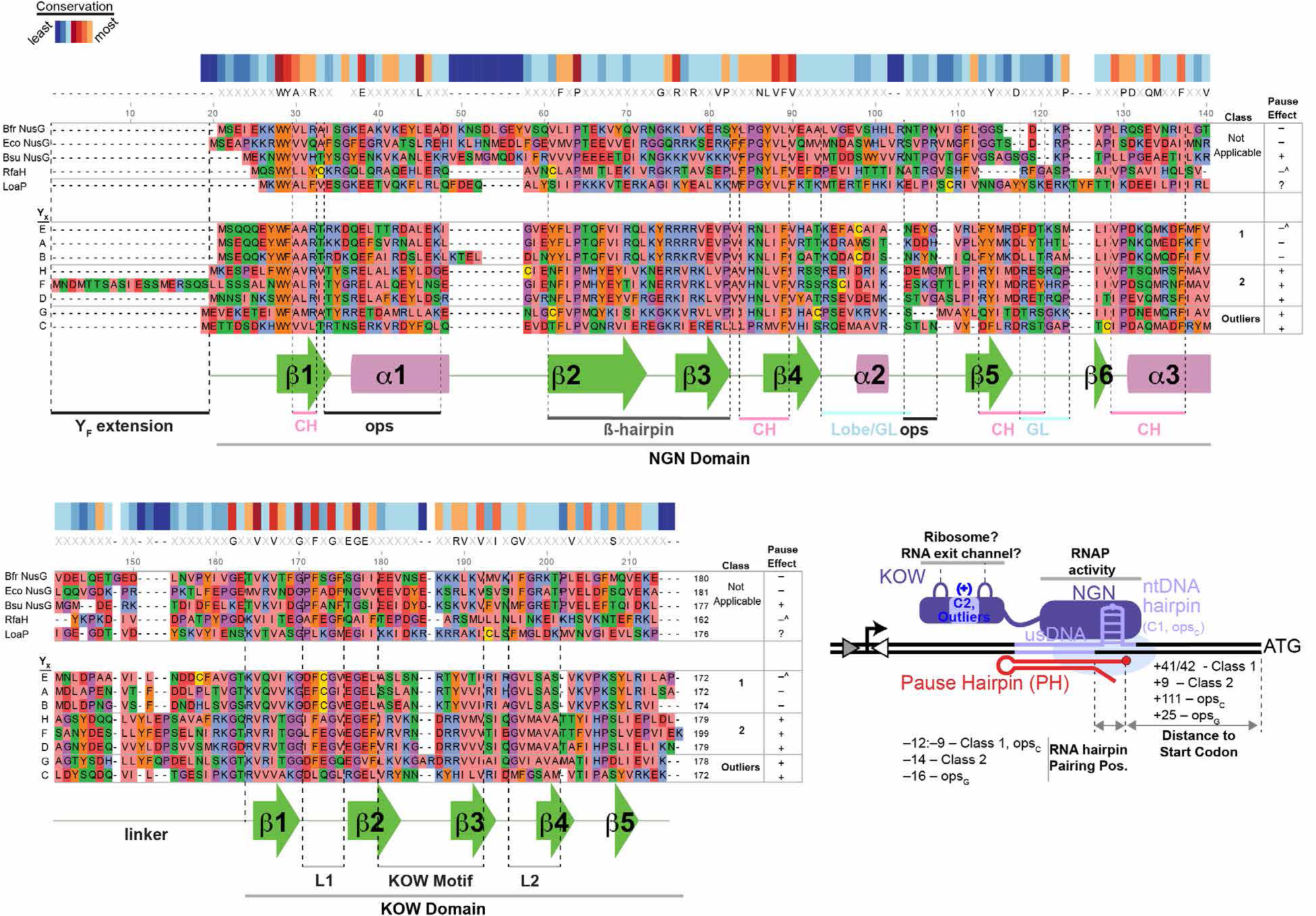
Y_X_ alignment compared to NusGs and orthologous factors. Sequences were aligned in SnapGene using the ClustalOmega algorithm. Amino acids are highlighted based on physico-chemical properties. ‘Pause effects’ indicate Y_X_ effects on pausing (‘^’ superscript indicates that RfaH^59^ or Y_E_ suppressed pausing at the pause site (*ops* or *ops*_E_), but enhanced pausing a few nucleotides downstream). *Bfr*NusG pause suppression is shown in Extended Data Fig. 4. *Eco*NusG pause suppression and *Bsu*NusG pause enhancement are documented^36,45,102,103^. Features depict some NusG_SP_-interacting modules of a PEC (lobe–gate loop [GL], clamp helices [CH]), *ops* ntDNAhp, and features of NusG_SP_ (NGN, KOW, hairpin). Bottom right: a cartoon summary of some class-specific features. Blue (+) indicates a positively KOW motif is found in Class 2 Y_X_ and Outlier Y_X_, similar to the positively charged KOW motif of LoaP^54^.

**Extended Data Fig. 8.**
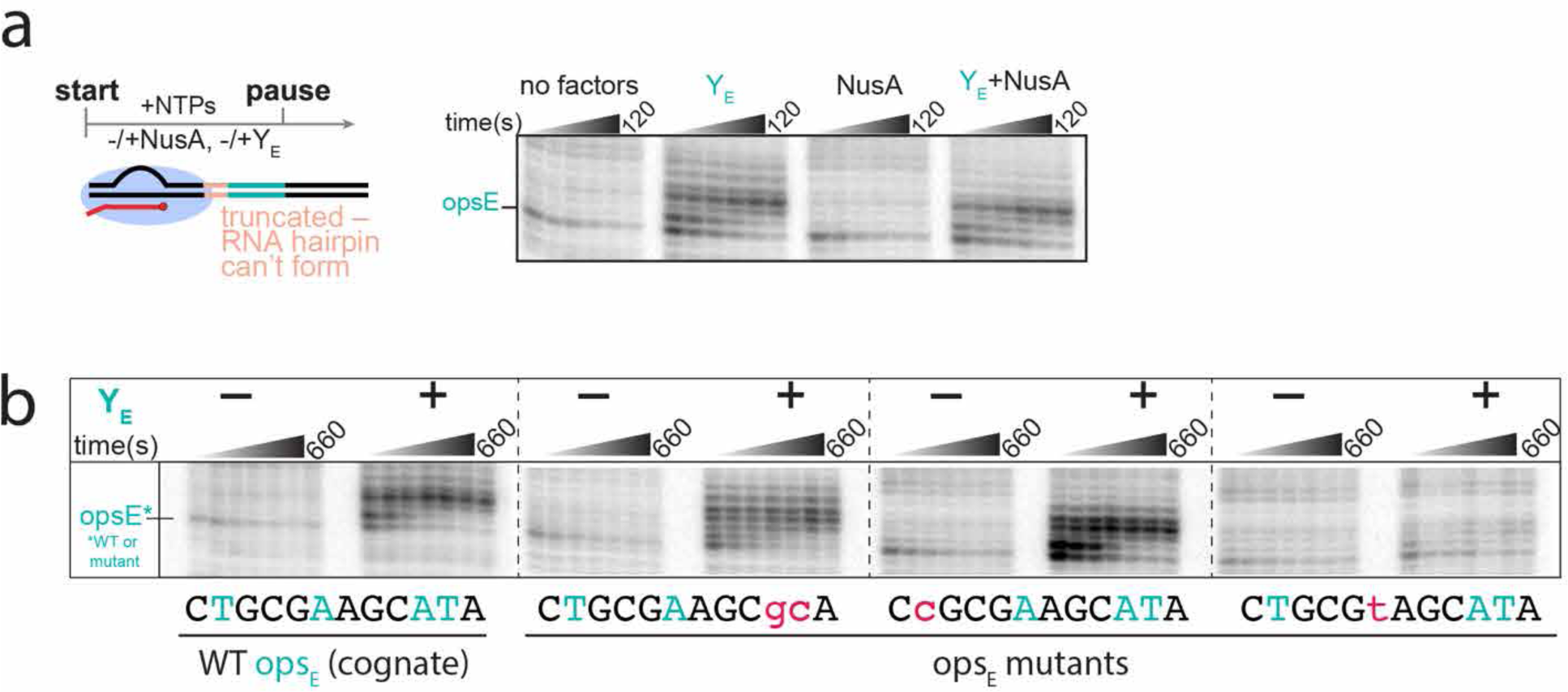
Effects of *ops*_E_ mutants on Y_E_ or NusA activity. **a)** NusA does not noticeably enhance pausing at *ops*_E_ in the absence of RNA hairpins. Experimental scheme testing the effect of NusA in the absence of upstream sequence enabling RNA hairpin formation. RNAP is reconstituted on a nucleic-acid scaffold with truncated upstream DNA. PIVoT assays performed with 150 nM Y_X_, 1 µM NusA, and 100 µM NTPs added concomitantly where indicated. **b)** Effects of substitutions within the –10:–1 *ops*_E_ window on Y_E_ activity. Results shown are representative of triplicate experiments.

**Extended Data Fig. 9.**
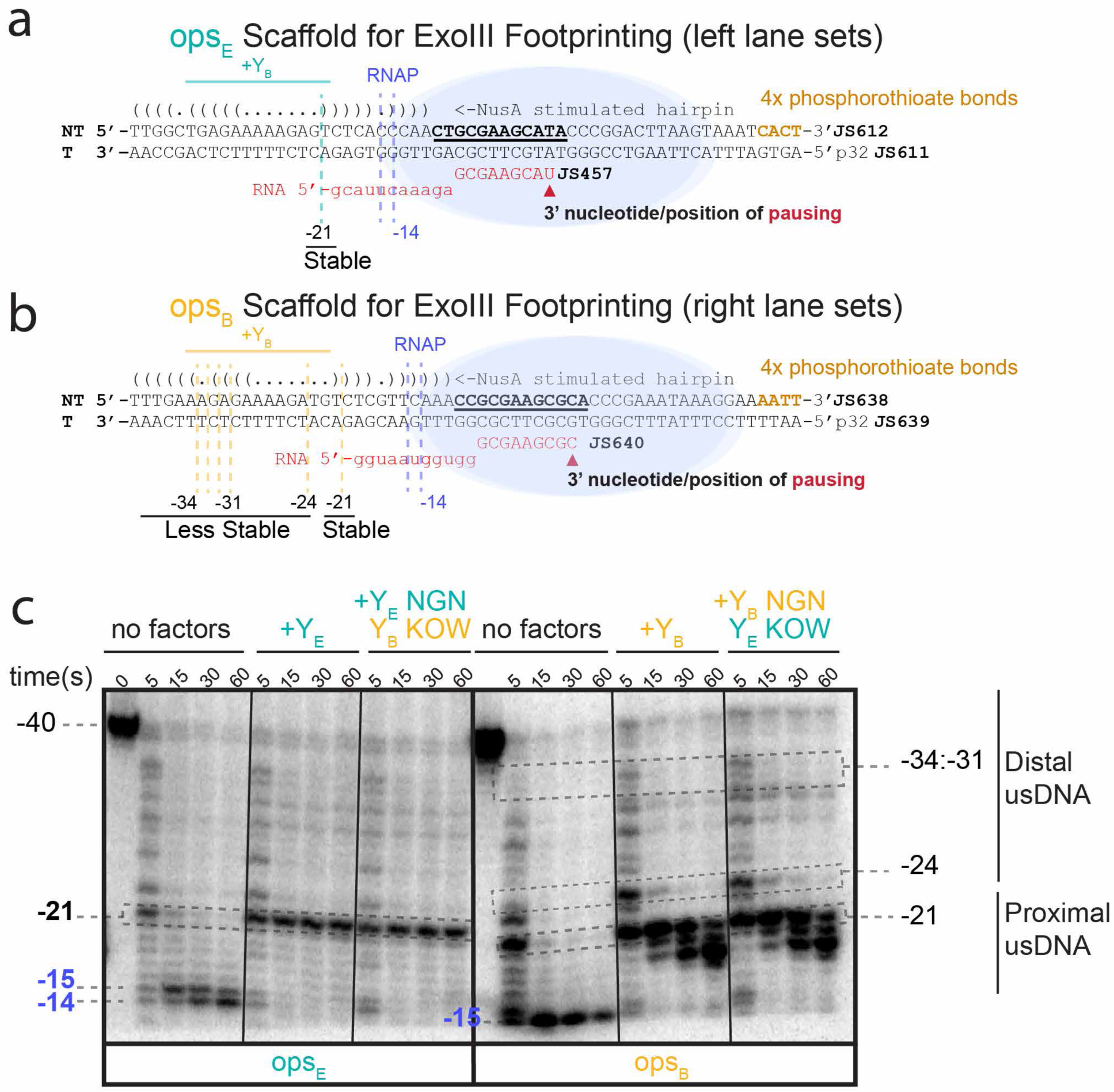
Primary data associated with Fig 6B. Representative of experiments performed in at least duplicate (see Methods). **a)** Nucleic acid scaffold used for exonuclease footprinting experiments mapping Y_E_ or hybrid Y_E_-Y_B_ NGN-KOW footprints on *ops_E_*. Footprints in the absence or presence of Y_B_ are indicated by dashed lines. 4x phosphorothioate bonds were incorporated at the 3’ end of the NT strand to prevent its cleavage and associated artifacts during the assay. **b)** Nucleic acid scaffold used for exonuclease footprinting experiments mapping Y_B_ or hybrid Y_B_-Y_E_ NGN-KOW footprints on *ops_B_*. Footprints in the absence or presence of Y_E_ are indicated by dashed lines. 4x phosphorothioate bonds were incorporated at the 3’ end of the NT strand to prevent its cleavage and associated artifacts during the assay. **c)** Representative exonuclease footprinting gel (n=2) illustrating that YB, but not YE, protects distal upstream DNA. These footprints were identical between WT and hybrid NGN-KOW proteins harboring an identical NGN domain, suggesting *ops_X_* specificity determinants are created by the NGN domain.

**Extended Data Fig. 10.**
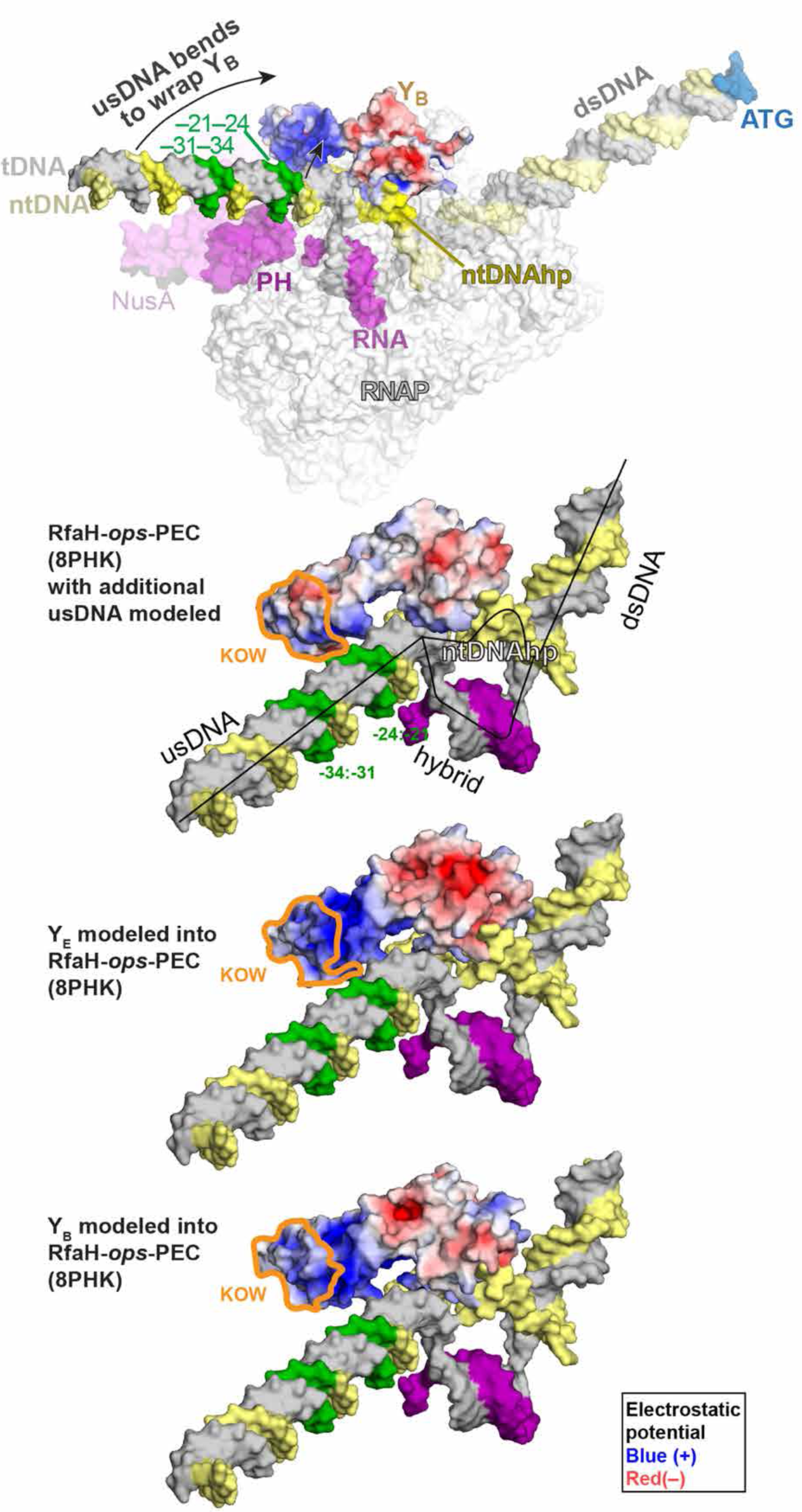
Modeling of Class 1 Y_X_ suggest that Y_X_ provides a larger positively charged surface for usDNA interaction relative to RfaH (*E. coli*). (top) Electrostatic surface potential model of Y_B_ recruited to *ops_X_* using the RfaH-*ops*-PEC (8PHK)^50^ as template and modeling additional upstream DNA. Green highlighted regions in the upstream DNA indicate Y_B_ footprints. The usDNA must distort to interact with sequence-specifically with Y_B_ (black arrows). The RfaH, Y_E_, and Y_B_ models below the full PEC model were created using the same structure but with RNAP and NusA are hidden for clarity. RfaH lacks the significant positive charge observed in models of Class 1 Y_E_ and Y_B_, suggesting this charge is an evolved feature of Y_X_ facilitating readout of upstream DNA. Most of the positive charge is localized to the NGN domain (KOW outlined in orange).

