## Supplementary Information for "*Bacteroides* expand the functional versatility of a universal transcription factor and transcribed DNA to program capsule diversity"

#### Tables, Notes, and Supplementary Figures for

Author affiliations: [1]Department of Biochemistry, University of Wisconsin–Madison, Madison, WI 53706, USA; [2]Microbiology Doctoral Training Program, University of Wisconsin–Madison, Madison, WI 53706, USA; [3]Department of Microbiology, University of Chicago, Chicago, IL, 60637, USA; [4]Duchossois Family Institute, University of Chicago, Chicago, IL 60637, USA [5]Cell and Molecular Biology Training Program, University of Wisconsin–Madison, Madison, WI 53706, USA; [6]Genetics Training Program, University of Wisconsin–Madison, Madison, WI 53706, USA; [7]Department of Bacteriology, University of Wisconsin–Madison, Madison, WI 53706, USA.

This PDF file includes:

1. Tables
   - Table S1. Oligonucleotides used in this study pS2
   - Table S2. *B. fragilis* strain construction pS11
   - Table S3. Plasmids used in this study pS13
   - Table S4. Strains used in this study pS16
2. Supplementary Notes
   - Nucleic scaffolds used in PIVoT and footprinting assays pS17
3. Supplementary Figures
   - Supplementary Fig. 1. Scaffolds used in PIVoT assays of Fig 1d,e and Extended Data Fig. 2. pS25
   - Supplementary Fig. 2. Predicted secondary structures of nascent RNA hairpins. pS26
   - Supplementary Fig. 3. WT $ops_X$ sequences and variants used for Fig. 5e pS27
   - Supplementary Fig. 4. Primary data associated with Fig 6c. pS28

**Table S1 – Oligonucleotides**

| Oligo | Sequence (5'-3') | Length (nt) | Purpose/Scaffold |
|---|---|---|---|
| JS145_T | CGTCACGAGTAGTAAGCTCCTGATCCTTACGGGTACGGGCAGCGAACCAATATTCTTGTTGTTGAGACATTTAGAAGGATATATTCCTGAGTGATTTACTTAAGTCCGGGTATGCTTCGCAGTTGGGTGAGACTCTTTTTCTCAGCCAAACTGCAAGATTTTTTATGACTTATTTTAGGTGGAAACGAACGGATTCGATC | 200 | PIVoT PSE ultramer scaffold |
| JS146_R | UUUUAACGUUUCCAC | 15 | PIVoT various scaffolds |
| JS147_NT | GATCGAATCCGTTAACCGATCTCTAAAATAAGTCATAAAAAATCTTGCAGTTTGGCTGAGAAAAAGAGTCTCACCCAACTGCGAAGCATACCCGGACTTAAGTAAATCACTCAGGAATATATCCTTCTAAATGTCTCAACAACAAGAATATTGGTTCGCTGCCCGTACCCGTAAGGATCAGGAGCTTACTACTCGTGACG | 200 | PIVoT PSE ultramer scaffold |
| JS510_NT | GCACTGAATTTTTCATTAAAATCATAGAGAAATAACTTGACCGGGAGGTATGCTTCGCTACTCCGGTGTCCCCAAAGAACATCCTGACGTGAACGACATGACAACATCCGCATCTATCGAATCTTCGATG | 130 | PIVoT PSF ultramer scaffold |
| JS511_T | CATCGAAGATTCGATAGATGCGGATGTTGTCATGTCGTTCACGTCAGGATGTTCTTTGGGGACACCGGAGTAGCGAAGCATACCTCCCGGTCAAGTTATGTAATGCGGATTTTAATGAAAAATTCAGTGC | 130 | PIVoT PSF ultramer scaffold |
| JS191_R | UUUUUUCCGCAUUAC | 15 | PIVoT PSF ultramer scaffold |
| JS708_NT | GATCGAATCCGTTAACCGATCTGTTCAAATCCGGTGCGAATACTTTTTCTGTACCCGATTATCATAAAACTTAATTTGTAAATTGCTGAAAATAAGGCATGTTTTTGAATATTCCTGTTTTAACAAATTTTCATCCTTAG | 200 | PIVoT PSC ultramer scaffold |

| | | | |
|---|---|---|---|
| | TCATTACTGAAACTTTTTCTTACGAACGTAGTCTTGGAGACAACAGATAGCGATAAAC | | |
| JS709_T | GTTTATCGCTATCTGTTGTCTCCAAGACTACGTTCGTAAGAAAAAGTTTCAGTAATGACTAAGGATGAAAATTTGTTAAAAACAGGAATATTCAAAAAACATGCCTTATTTTCAGCAATTTACAAATTAAGTTTTATGATAATCGGGTACAGAAAAAGTATTCGCACCGGATTTGAACGTGGAAACGAACGGATTCGATC | 200 | PIVoT PSC ultramer scaffold |
| JS551_NT_opsB (+escHP) | GTCCGTTCGTTTCCACGAATTTGACGGTTTGAAAGAGAAAAGATGTCTCGTTCAAACCGCGAAGCGCACCCGAAATAAAGGAAAATTTCC | 90 | opsB with pause and escape hp (reconstitute upstream of RNA hp) |
| JS552_T_opsB(+escHP) | GGAAATTTTCCTTTATTTCGGGTGCGCTTCGCGGTTTGAACGAGACATCTTTTCTCTTTCAAACCGTCAAATTCGTGGAAACGAACGGAC | 90 | opsB with pause and escape hp (reconstitute upstream of RNA hp) |
| JS553_NT_opsA (+escHP) | GTCCGTTCGTTTCCACGTCTTACGGTTTGAATGGGAAAAGATGTCTCGTCCAAACCGCGTAGCGCACCCGAAAGTAACCTCTCGACTTTG | 90 | opsA with pause and escape hp (reconstitute upstream of RNA hp) |
| JS554_T_opsA(+escHP) | CAAAGTCGAGAGGTTACTTTCGGGTGCGCTACGCGGTTTGGACGAGACATCTTTTCCCATTCAAACCGTAAGACGTGGAAACGAACGGAC | 90 | opsA with pause and escape hp (reconstitute upstream of RNA hp) |

| | | | |
|---|---|---|---|
| JS521_NT_opsE | GTCCGTTCGTTTCCACGATCTTGCAGTTTGGCTGAGAAAAAGAGTCTCACCCAACTGCGAAGCATACCCGGACTTAAGTAAATCACTCAG | 90 | opsE with pause hairpin (reconstitute upstream of RNA hp) |
| JS522_T_opsE | CTGAGTGATTTACTTAAGTCCGGGTATGCTTCGCAGTTGGGTGAGACTCTTTTTCTCAGCCAAACTGCAAGATCGTGGAAACGAACGGAC | 90 | opsE with pause hairpin (reconstitute upstream of RNA hp) |
| JS523_NT_opsF | GTCCGTTCGTTTCCACGTAACTTGACCGGGAGGTATGCTTCGCTACTCCGGTGTCCCCAAAGAACATCCTGACGTGAACGACATGACAAC | 90 | opsF with pause hairpin (reconstitute upstream of RNA hp) |
| JS524_T_opsF | GTTGTCATGTCGTTCACGTCAGGATGTTCTTTGGGGACACCGGAGTAGCGAAGCATACCTCCCGGTCAAGTTACGTGGAAACGAACGGAC | 90 | opsF with pause hairpin (reconstitute upstream of RNA hp) |
| JS583_NT_opsH | GTCCGTTCGTTTCCACGTCACCTGACCGGGAGGTACTTTCGTACTCCGGTGTCCCCAAAGAACATCCTTTTGTGAAGGAATCCCCAGAGC | 90 | opsH with pause hairpin (reconstitute upstream of RNA hp) |
| JS584_T_opsH | GCTCTGGGGATTCCTTCACAAAAGGATGTTCTTTGGGGACACCGGAGTACGAAAGTACCTCCCGGTCAGGTGACGTGGAAACGAACGGAC | 90 | opsH with pause hairpin (reconstitute |

| | | | |
|---|---|---|---|
| | | | upstream of RNA hp) |
| JS540_PSE-asDNA_6 | CTGCAAGATCGTGGA | 15 | asDNAs to probe extent of RNA hairpin in PSE |
| JS541_PSE-asDNA_7 | ACTGCAAGATCGTGG | 15 | asDNAs to probe extent of RNA hairpin in PSE |
| JS542_PSE-asDNA_8 | AACTGCAAGATCGTG | 15 | asDNAs to probe extent of RNA hairpin in PSE |
| JS543_PSE-asDNA_9 | AAACTGCAAGATCGT | 15 | asDNAs to probe extent of RNA hairpin in PSE |
| JS544_PSE-asDNA_10 | CAAACTGCAAGATCG | 15 | asDNAs to probe extent of RNA hairpin in PSE |
| JS545_PSE-asDNA_11 | CCAAACTGCAAGATC | 15 | asDNAs to probe extent of RNA hairpin in PSE |
| JS566_PSE-asDNA_17 | GTGCTAGAACGTCAAACCGACTCT | 24 | negative control asDNA |
| JS555_NT | GTCCGTTCGTTTCCACGTCTTACGGTTTGAATGGGAAAAGATGTCTCGTCCAAACTGCGAAGCATACCCGAAAGTAACCTCTCGACTTTG | 90 | $HP_A$,$[-10:-1]_E$ (Pause Cycling Observed) |

| | | | |
|---|---|---|---|
| JS556_T | CAAAGTCGAGAGGTTACTTTCGGGTATGCTTCGCAGTTTGGACGAGA CATCTTTTCCCATTCAAACCGTAAGACGTGGAAACGAACGGAC | 90 | $HP_{A}$,$[-10:-1]_{E}$ (Pause Cycling Observed) |
| JS557_NT | GTCCGTTCGTTTCCACGATCTTGCAGTTTGGCTGAGAAAAAGAGTCT CACCCAACCGCGTAGCGCACCCGGACTTAAGTAAATCACTCAG | 90 | $HP_{E}$,$[-10:-1]_{A}$ (Pause Cycling Observed) |
| JS558_T | CTGAGTGATTTACTTAAGTCCGGGTGCGCTACGCGGTTGGGTGAGAC TCTTTTTCTCAGCCAAACTGCAAGATCGTGGAAACGAACGGAC | 90 | $HP_{E}$,$[-10:-1]_{A}$ (Pause Cycling Observed) |
| JS591_NT | GTCCGTTCGTTTCCACGATCTTGCAGTTTGGCTGAGAAAAAGAGTCT CACCCAACTGCGtAGCATACCCGGACTTAAGTAAATCACTCAG | 90 | -6 opsE>opsA |
| JS592_T | CTGAGTGATTTACTTAAGTCCGGGTATGCTaCGCAGTTGGGTGAGAC TCTTTTTCTCAGCCAAACTGCAAGATCGTGGAAACGAACGGAC | 90 | -6 opsE>opsA |
| JS593_NT_mut opsE+hp -1,-2,-10 | GTCCGTTCGTTTCCACGATCTTGCAGTTTGGCTGAGAAAAAGAGTCT CACCCAACcGCGAAGCgcACCCGGACTTAAGTAAATCACTCAG | 90 | opsE+hp > -10:1 opsB |
| JS594_T_mut opsE+hp -1,-2,-10 | CTGAGTGATTTACTTAAGTCCGGGTgcGCTTCGCgGTTGGGTGAGACT CTTTTTCTCAGCCAAACTGCAAGATCGTGGAAACGAACGGAC | 90 | opsE+hp > -10:1 opsB |
| JS595_NT_mut opsE+hp -1,-2,-10,-14 | GTCCGTTCGTTTCCACGATCTTGCAGTTTtGCTGAGAAAAAGAGTCTC ACCaAACcGCGAAGCgcACCCGGACTTAAGTAAATCACTCAG | 90 | opsE+hp > -10:1, -14:-36 opsB |
| JS596_T_mut opsE+hp -1,-2,-10,-14 | CTGAGTGATTTACTTAAGTCCGGGTgcGCTTCGCgGTTtGGTGAGACTC TTTTTCTCAGCaAAACTGCAAGATCGTGGAAACGAACGGAC | 90 | opsE+hp > -10:1, -14:-36 opsB |

| | | | |
|---|---|---|---|
| JS597_NT_mut opsE+hp -1,-2,-10,insert U-A | GTCCGTTCGTTTCCACGATCTTGCAGTTTGGCTGAGAAAAAGAGTCTCACCCAAaCcGCGAAGCgcACCCGGACTTAAGTAAATCACTCA | 90 | opsE+hp > -10:1, -14 ins opsB |
| JS598_T_mut opsE+hp -1,-2,-10,insert U-A | TGAGTGATTTACTTAAGTCCGGGTgcGCTTCGCgGtTTGGGTGAGACTCTTTTTCTCAGCCAAACTGCAAGATCGTGGAAACGAACGGAC | 90 | opsE+hp > -10:1, -14 ins opsB |
| JS599_NT_mut opsE+hp -1,-2,-10,insert U-A, change var pairing region | GTCCGTTCGTTTCCACGATCTTGCAGTTTGaaaGAGAAAAAGAGTCTCgttCAAaCcGCGAAGCgcACCCGGACTTAAGTAAATCACTCA | 90 | opsE+hp > -10:1, -14 ins, -36:-34, -18:-16 opsB |
| JS600_T_mut opsE+hp -1,-2,-10,insert U-A, change var pairing region | TGAGTGATTTACTTAAGTCCGGGTgcGCTTCGCgGtTTGaacGAGACTCTTTTTCTCtttCAAACTGCAAGATCGTGGAAACGAACGGAC | 90 | opsE+hp > -10:1, -14 ins, -36:-34, -18:-16 opsB |
| JS672_NT | GTCCGTTCGTTTCCACGATCTTGCAGTTTGaaaGAGAAAAgatGTCTCgttCAAaCcGCGAAGCgcACCCGGACTTAAGTAAATCACTCA | 90 | opsE+hp > (mostly opsB now) -14 ins, -36:-34, -18:-16, -26:-24 opsB |
| JS673_T | TGAGTGATTTACTTAAGTCCGGGTgcGCTTCGCgGtTTGaacGAGACatcTTTTCTCtttCAAACTGCAAGATCGTGGAAACGAACGGAC | 90 | opsE+hp > (mostly opsB now) -14 ins, -36:-34, -18:-16, -26:-24 opsB |

| | | | |
|---|---|---|---|
| JS674_NT | GTCCGTTCGTTTCCACGATCTTGCAGTTTGaaaGAGAAAAgatGTCTCgttCAAaCcGCGAAGCgcACCCGaaataaaggaaaatttccc | 90 | opsE+hp > (opsB-$_{\text{escape hairpin}}$) -14 ins, -36:-34, -18:-16, -26:-24, dsDNA opsB |
| JS675_T | gggaaattttcctttatttCGGGTgcGCTTCGCgGtTTGaacGAGACatcTTTTCTCtttCAAACTGCAAGATCGTGGAAACGAACGGAC | 90 | opsE+hp > (opsB-$_{\text{escape hairpin}}$) -14 ins, -36:-34, -18:-16, -26:-24, dsDNA opsB |
| JS666_R_opsB-3 | UUUUUUCCGCGAAGC | 15 | opsB-hp (reconstitute -3) |
| JS654_NT_opsB | TTTGAAAGAGAAAAGATGTCTCGTTCAAACCGCGAAGCGCACCCGAAATAAAGGAAAATTTCCCCAAG | 68 | opsB-hp (reconstitute -3) |
| JS655_T_opsB | CTTGGGGAAATTTTCCTTTATTTCGGGTGCGCTTCGCGGTTTGAACGAGACATCTTTTCTCTTTCAAA | 68 | opsB-hp (reconstitute -3) |
| JS660_NT_opsB -36-34 | TTTGggcGAGAAAAGATGTCTCGTTCAAACCGCGAAGCGCACCCGAAATAAAGGAAAATTTCCCCAAG | 68 | opsB -36-34 mut to opsE |
| JS661_T_opsB-36-34 | CTTGGGGAAATTTTCCTTTATTTCGGGTGCGCTTCGCGGTTTGAACGAGACATCTTTTCTCgccCAAA | 68 | opsB -36-34 mut to opsE |
| JS662_NT_opsB -26-24 | TTTGAAAGAGAAAAaAgGTCTCGTTCAAACCGCGAAGCGCACCCGAAATAAAGGAAAATTTCCCCAAG | 68 | opsB -26-24 mut to opsE |
| JS663_T_opsB-26-24 | CTTGGGGAAATTTTCCTTTATTTCGGGTGCGCTTCGCGGTTTGAACGAGACcTtTTTTCTCTTTCAAA | 68 | opsB -26-24 mut to opsE |

| | | | |
|---|---|---|---|
| JS664_NT_opsB -18-16 | TTTGAAAGAGAAAAGATGTCTCcacCAAACCGCGAAGCGCACCCGAAATAAAGGAAAATTTCCCCAAG | 68 | opsB -18-16 mut to opsE |
| JS665_T_opsB-18-16 | CTTGGGGAAATTTTCCTTTATTTCGGGTGCGCTTCGCGGTTTGgtgGAGACATCTTTTCTCTTTCAAA | 68 | opsB -18-16 mut to opsE |
| JS467_NT-opsE-mut1 | GAGAAAAAGAGTCTCACCCAACTGCGAAGCgcACCCGGACTTAAGTAAATCACTCAGGAA | 60 | opsE_mut1(opsA/B-mimic-1(AT□GC)) |
| JS468_T-opsE-mut1 | TTCCTGAGTGATTTACTTAAGTCCGGGTgcGCTTCGCAGTTGGGTGAGACTCTTTTTCTC | 60 | opsE_mut1(opsA/B-mimic-1(AT□GC)) |
| JS469_NT-opsE-mut2 | GAGAAAAAGAGTCTCACCCAACcGCGAAGCATACCCGGACTTAAGTAAATCACTCAGGAA | 60 | opsE_mut2(opsA/B-mimic-2(T□C)) |
| JS470_T-opsE-mut2 | TTCCTGAGTGATTTACTTAAGTCCGGGTATGCTTCGCgGTTGGGTGAGACTCTTTTTCTC | 60 | opsE_mut2(opsA/B-mimic-2(T□C)) |
| JS471_NT-opsE-mut3 | GAGAAAAAGAGTCTCACCCAACTGCGtAGCATACCCGGACTTAAGTAAATCACTCAGGAA | 60 | opsE_mut3(opsA-mimic-3(A□T)) |
| JS472_T-opsE-mut3 | TTCCTGAGTGATTTACTTAAGTCCGGGTATGCTaCGCAGTTGGGTGAGACTCTTTTTCTC | 60 | opsE_mut3(opsA-mimic-3(A□T)) |
| 9563 | GGTCAGTACGTCCGGCATAGTTGCGCCCGTAAATTCAGATCTTCCAGTGG | 50 | consensus pause |
| 8334 | CCACTGGAAGATCTGAATTTACGGGCGCAACTATGCCGGACGTACTGACC | 50 | consensus pause |

| | | | |
|---|---|---|---|
| 8342 | UUUUUUGGCAUAGUU | 15 | consensus pause |
| JS563 | GGTCAGTACGTCCTT*TTGGTGGT*CGTTGTAGTGGGCAGATCTTCCAGTGG | 50 | anticonsensus pause |
| 8954 | CCACTGGAAGATCTGCCCACTACAACGACCACCAAAAGGACGTACTGACC | 50 | anticonsensus pause |
| 8952 | UUUUUUUUUUGGUGG | 15 | anticonsensus pause |
| 14047 | /5Phos/NNNNNNNNNNGCAGCTCTGTAGGCACCATCAATGATCGTCGGA/3ddC/ | 43 | NET-seq |
| 14637 | /5Phos/AGATCGGAAGAGCACACGTCTGAAC/iSp18/CACTCA/iSp18/CCTACACGACGCTCTTCCGATCTTCCGACGATCATTGATGGTGCCTACAG | 81 | NET-seq |
| 14641 | AATGATACGGCGACCACCGAGATCTACACAGCGAGCTACACTCTTTCCCTACACGACGCTCTTCCGATCTTCCGACGATC | 80 | NET-seq |
| 14645 | CAAGCAGAAGACGGCATACGAGATAAGGATGAGTGACTGGAGTTCAGACGTGTGCTCTTCCGATCT | 66 | NET-seq |

**Table S2 – *B. fragilis* Strain Construction**

| Purpose | Plasmid name | Primers | Sequence |
|---|---|---|---|
| '[-10:-1]opsA in PSE' = Clone opsA region into PSE region into pLGB13 - left flank amplified from Δmpi M44 | pKF32 | opsA-PSE-LF_F | taagattagcattatgagtgccggctgtggaggcaattatc |
| | | opsA-PSE-LF_R | acgcggttgggtgagactca |
| | | opsA-PSE-RF_F | agagtctcacccaaccgcgtagcgcacccggacttaag |
| | | opsA-PSE-RF_R | cgaattcctgcagcccggggcagtctcctgctcatctatcg |
| Replacing opsE (+PSE hairpin) with opsA (+PSA hairpin) in pLGB13 | pKF52 | HPops-LF_F | taagattagcattatgagtgcctgggaacaaaacaaag |
| | | HPops-LF_R | attcaaaccgactgcaagattttttatgac |
| | | HPopsA_F | atcttgcagtcggtttgaatgggaaaagatg |
| | | HPopsA_R | taagtccgggtgcgctacgcggtttgga |
| | | HPops-RF_F | gcgtagcgcacccggacttaagtaaatcac |
| | | HPops-RF_R | cgaattcctgcagcccggggcagtctcctgctcatctatc |
| Clone rpoC-3X-FLAG in pLGB13 - left flank amplified from Jason's construct pJS039_Bf-rpoC-3X-FLAG, right flank amplified from B. fragilis 9343 genome | pKF36 | rpoC-3X_FLAG-LF_fwd | taagattagcattatgagtgctgtaggtggacacgtag |

| | | | |
|---|---|---|---|
| | | rpoC-3X_FLAG-LF_rev | ttgaattgtatcatttgtcgtcatcgtc |
| | | rpoC_3X_FLAG-RF_fwd | cgacaaatgatacaattcaattgactatagctgaaaaag |
| | | rpoC_3X_FLAG-RF_rev | cgaattcctgcagcccggggcgcattcaaccaagacgc |

**Table S3 – Plasmids**

| Plasmid | Lab Stock# | Purpose | Properties | Construction | Parent (s) | Anti-biotic |
|---|---|---|---|---|---|---|
| pJS011 | 5903 | T7 OEP | Bf UpaY-intein-CBD | pTYB2 (Stock 182) backbone, UpaY from Bfragilis NCTC 9343 | pTYB2 (Stock 182) | Amp |
| pJS015 | 5906 | T7 OEP | BfRNAP, strep-rpoB, rpoC-his10ppx | pRM756 backbone, codon-optimized BfRNAP | pRM756 | Kan |
| pJS019 | 5910 | T7 OEP | Bf UpeY-inteinCBD (CTD Ala insertion for optimal cleavage) | pTYB2 (Stock 182) backbone, UpeY from Bfragilis NCTC 9343 | pTYB2 (Stock 182) | Amp |
| pJS021 | 5912 | T7 OEP | Bf UpaZ-inteinCBD (CTD Ala insertion for optimal cleavage) | pTYB2 (Stock 182) backbone, UpaZ from Bfragilis NCTC 9343 | pTYB2 (Stock 182) | Amp |
| pJS022 | 5913 | T7 OEP | Bf UpfY-inteinCBD (CTD Ala insertion for optimal cleavage efficiency) | pTYB2 (Stock 182) backbone, UpfY from Bfragilis NCTC 9343 | pTYB2 (Stock 182) | Amp |
| pJS028 | 5917 | T7 OEP | Bf UphY-inteinCBD (CTD Ala insertion for optimal cleavage efficiency) | pTYB2 (Stock 182) backbone, UphY from Bfragilis NCTC 9343 | pTYB2 (Stock 182) | Amp |
| pJS031 | 5920 | T7 OEP | BfGre-inteinCBD (CTD Ala insertion for optimal cleavage efficiency) | pTYB2 (Stock 182) backbone, Gre from Bfragilis NCTC 9343 | pTYB2 (Stock 182) | Amp |

| | | | | | | |
|---|---|---|---|---|---|---|
| pJS034 | 5922 | T7 OEP | Bf UpbY-inteinCBD (CTD Ala insertion for optimal cleavage efficiency) | pTYB2 (Stock 182) backbone, UpbY from Bfragilis NCTC 9343 | pTYB2 (Stock 182) | Amp |
| pJS036 | 5924 | T7 OEP | Bf UpeZ-inteinCBD (CTD Ala insertion for optimal cleavage efficiency) | pTYB2 (Stock 182) backbone, UpeZ from Bfragilis NCTC 9343 | pTYB2 (Stock 182) | Amp |
| pJS044 | 5932 | T7 OEP | NusG-inteinCBD (CTD Ala insertion for optimal cleavage efficiency) | pTYB2 (Stock 182) backbone, NusG from Bfragilis NCTC 9343 | pTYB2 (Stock 182) | Amp |
| pJS045 | 5934 | T7 OEP | NusA-inteinCBD (CTD Ala insertion for optimal cleavage efficiency) | pTYB2 (Stock 182) backbone, NusA from Bfragilis NCTC 9343 | pTYB2 (Stock 182) | Amp |
| pJS058 | 5943 | T7 OEP | Bf UpeY(NTD)-inteinCBD (CTD Ala insertion for optimal cleavage efficiency) | pTYB2 (Stock 182) backbone, fragments from Bfragilis NCTC 9343 | pTYB2 (Stock 182) | Amp |
| pJS059 | 5944 | T7 OEP | Bf UpfY(NTD)-inteinCBD (CTD Ala insertion for optimal cleavage efficiency) | pTYB2 (Stock 182) backbone, fragments from Bfragilis NCTC 9343 | pTYB2 (Stock 182) | Amp |
| pJS060 | 5945 | T7 OEP | Avi-UpeY-inteinCBD (CTD Ala insertion for optimal cleavage efficiency) | pTYB2 (Stock 182) backbone, 2 oligos for Avi tag, fragments from Bfragilis NCTC 9343 | pTYB2 (Stock 182) | Amp |
| pJS063 | 5948 | T7 OEP | Bf eY(NGN)-bY(KOW)-intein-CBD (CTD Ala insertion for optimal cleavage efficiency) | pTYB2 (Stock 182) backbone, eY NGN and bY KOW from Bfragilis NCTC 9343 | pTYB2 (Stock 182) | Amp |

| | | | | | | |
|---|---|---|---|---|---|---|
| pJS064 | 5949 | T7 OEP | Bf bY(NGN)-eY(KOW)-inteinCBD (CTD Ala insertion for optimal cleavage efficiency) | pTYB2 (Stock 182) backbone, bY NGN and eY KOW from Bfragilis NCTC 9343 | pTYB2 (Stock 182) | Amp |
| pJS073 | 5958 | T7 OEP | Bf UpcY-intein CBD (CTD Ala insertion for optimal cleavage efficiency) | pTYB2 (Stock 182) backbone, UpcY from Bfragilis NCTC 9343 | pTYB2 (Stock 182) | Amp |
| pKF32 | 5963 | Test ops replacement effects on PS expression | [-10:-1]opsA in PSE = "opsE>opsE/A" in figure | See methods and Table S2 | | |
| pKF36 | 5966 | epitope tag RNAP | rpoC-3xFLAG | See methods and Table S2 | | |
| pKF52 | | Test ops replacement effects on PS expression | [-43:-1]opsA in PSE = "opsE>opsA" in figure | See methods and Table S2 | | |

**Table S4 – Strains used in this study**

| **Strain** | **Notes** | **Source** |
|---|---|---|
| *B. fragilis* NCTC 9343 | wild type strain from which all mutants were derived | ATCC |
| *B. fr*agilis 9343 ΩPSE | insertional mutant in PSE locus abrogating PSE synthesis | Krinos *et al.* 2001 |
| *B. fragilis* 9343 Δ*mpi* M44 | Promoter PSA and PSE locked on. (PSC constitutive) | Coyne *et al.* 2003 |
| *B. fragilis* 9343 Δ*mpi* M44 Δ*upaZ* | Promoter PSA and PSE locked on. (PSC constitutive). upaZ deletion | Chatzidaki-Livanis *et al.* 2010 |
| *B. fragilis* 9343 mpiM44 [-10:-1]opsE>[-10:-1]opsA | replacing opsE [-10:-1] with opsA [-10:-1] | This study |
| *B. fragilis* 9343 mpiM44 HP-opsE>HP-opsA | replacing opsE [-38:-1] with opsA [-38:-1] | This study |
| RL3569 | E. coli B F$^{-}$ ompT gal [dcm$^{-}$] [lon$^{-}$] hsdS$^{-}$ (rB$^{-}$ mB$^{-}$) [λDE3(lacUV5::T7 RNAP) imm21 int$^{-}$-Δnin5 ΔEcoRI(21226-26104) BamHI27972$^{-}$] pRARE2<br><br>Rif-resistant expression strain (S522F) for expressing recombinant RNA polymerases. Made by transduction of RL1674 with P1 lysate of Gourse Lab strain (RLG3360). RLG3360 is stored in our database as RL3570. | This study |
| *B. fragilis* 9343 *rpoC*-FLAG | 3x FLAG tag on C-terminus of b' subunit to enable immunoprecipitation | This study |

#### Nucleic Acid Scaffolds for PIVoT

#### Extended Data Fig 2bc – Consensus vs. Anticonsensus Pause

##### Consensus Pause

5'-GGTCAGTACGTCCGGCATAGTTGCGCCCGTAAATTCAGATCTTCCAGTGG-3′ 9563
3′-CCAGTCATGCAGGCCGTATCAACGCGGGCATTTAAGTCTAGAAGGTCACC-5' 8334
GCAUAGUU 8342
5'-UUUUUUG

##### Anticonsensus Pause (fully complementary)

5′-GGTCAGTACGTCCTT*TTGGTGGT*CGTTGTAGTGGGCAGATCTTCCAGTGG-3′ JS563
3′-CCAGTCATGCAGGAAAACCACCAGCAACATCACCCGTCTAGAAGGTCACC-5′ 8954
UUUUGGUGG 8952
5′-UUUUUU

#### Extended Data Fig 2d – leader pause scanning 3 classes

##### Class 1 PSE Ultramer (bubble; NT changed)

**NT 5′**-GATCGAATCCGTTAACCGATCTCTAAAATAAGTCATAAAAAATCTTGCAGTTTGGCTGAGAAAAAGAGTCTCACCCAA**CTGCGAAGCATA**CCCGGACTTAAGTAAATCACTCAGGAATATATCCTTCTAAATGTCTCAACAACAAGAATATTGGTTCGCTGCCCGTACCCGTAAGGATCAGGAGCTTACTACTCGTGACG-3′ **JS147**
**T 3′** -CTAGCTTAGGCAAGCAAAGGTGGATTTTATTCAGTATTTTTTAGAACGTCAAACCGACTCTTTTTCTCAGAGTGGGTTGACGCTTCGTATGGGCCTGAATTCATTTAGTGAGTCCTTATATAGGAAGATTTACAGAGTTGTTGTTCTTATAACCAAGCGACGGGCATGGGCATTCCTAGTCCTCGAATGATGAGCACTGC-5′ **JS145**
**RNA** 5′- UUUUAA**CGUUUCCAC**-3′ **JS146**

##### Class 2 PSF Ultramer 2 - test putative Class II RNA recognition mechanism (bubble; NT changes in purple)

(((((..((((.((...))))))))))) <- nusA stimulated pause
5′-GCACTGAATTTTTCATTAAAATC**ATAGAGAA**ATAACTTGACCGGGAGGTATGCTTCGCTACTCCGGTGT**CCCCAAAGAACA**TCCTGACGTGAACGACATGACAACATCCGCATCTATCGAATCTTCGATG-3′**JS510**
3′-CGTGACTTAAAAAGTAATTTTAGGCGTAATGTATTGAACTGGCCCTCCATACGAAGCGATGAGGCCACAGGGGTTTCTTGTAGGACTGCACTTGCTGTACTGTTGTAGGCGTAGATAGCTTAGAAGCTAC-5′**JS511**
RNA 5′-UUUUUUC**CGCAUUAC** **JS191**

##### PSC

**Bubble Ultramer Scaffold # 1 from PSC 5′UTR (bold region not from UTR)**
**NT 5′-GATCGAATCCGTTAACCGATCTG**TTCAAATCCGGTGCGAATACTTTTTCTGTAC**CCGATTATCA**TAAAACTTAATTTGTAAATTGCTGAAAATAAGGCATGTTTTTTGAATATTCCTGTTTTTAACAAATTTTCATCCTTAGTCATTACTGAAACTTTTTCTTACGAACGTAGTCTTGGAGACAACAGATAGCGATAAAC-3′ **JS708**
**T 3′ -CTAGCTTAGGCAAGCAAAGGTGC**AAGTTTAGGCCACGCTTATGAAAAAGACATGGGCTAATAGTATTTTGAATTAAACATTTAACGACTTTTATTCCGTACAAAAAACTTATAAGGACAAAAATTGTTTAAAAGTAGGAATCAGTAATGACTTTGAAAAAGAATGCTTGCATCAGAACCTCTGTTGTCTATCGCTATTTG-5′ **JS709**
**RNA** 5′- UUUUAA**CGUUUCCAC**-3′ **JS146**

#### Fig 1d (pause and capture next to ladder in Ext Data Fig. 2e)

##### opsE+hp (Class 1)

((((.(((((.......))))).))))<-NusA stimulated pause hairpin
**NT 5′-GTCCGTTCGTTTCCACG**ATCTTGCAGTTTGGCTGAGAAAAAGAGTCTCACCCAA**CTGCGAAGCATA**CCCGGACTTAAGTAAATCACTCAG-3′ **JS521**
**T 3′-CAGGCAAGCAAAGGTGC**TAGAACGTCAAACCGACTCTTTTTCTCAGAGTGGGTTGACGCTTCGTATGGGCCTGAATTCATTTAGTGAGTC-5′ **JS522**
**RNA** 5′-UUUUAA**CGUUUCCAC** **JS146**

#### Fig 1e, Extended Data Fig. 2e

#### opsE+hp (Class 1)

```
                                    ((((.(((((.......))))).))))<-NusA stimulated pause hairpin
NT 5'-GTCCGTTCGTTTCCACGATCTTGCAGTTTGGCTGAGAAAAAGAGTCTCACCCAACTGCGAAGCATACCCGGACTTAAGTAAATCACTCAG-3'  JS521
T  3'-CAGGCAAGCAAAGGTGCTAGAACGTCAAACCGACTCTTTTTCTCAGAGTGGGTTGACGCTTCGTATGGGCCTGAATTCATTTAGTGAGTC-5'  JS522
RNA 5'-UUUUAACGUUUCCAC   JS146
```

#### opsF+hp (Class 2)

```
                                 (((((..((((.((...))))))))))) <-NusA-stimulated pause hairpin
NT 5'-GTCCGTTCGTTTCCACGTAACTTGACCGGGAGGTATGCTTCGCTACTCCGGTGTCCCCAAAGAACATCCTGACGTGAACGACATGACAAC-3'JS523
T  3'-CAGGCAAGCAAAGGTGCATTGAACTGGCCCTCCATACGAAGCGATGAGGCCACAGGGGTTTCTTGTAGGACTGCACTTGCTGTACTGTTG-5'JS524
RNA 5'-UUUUAACGUUUCCAC   JS146
```

### Extended Data Fig. 3

#### opsE+hp (Class 1)

```
                                    ((((.(((((.......))))).))))<-NusA stimulated pause hairpin
NT 5'-GTCCGTTCGTTTCCACGATCTTGCAGTTTGGCTGAGAAAAAGAGTCTCACCCAACTGCGAAGCATACCCGGACTTAAGTAAATCACTCAG-3'  JS521
T  3'-CAGGCAAGCAAAGGTGCTAGAACGTCAAACCGACTCTTTTTCTCAGAGTGGGTTGACGCTTCGTATGGGCCTGAATTCATTTAGTGAGTC-5'  JS522
RNA 5'-UUUUAACGUUUCCAC   JS146
```

#### opsB+hp (Class 1)

```
                                 (((((((((.((((.......)))).))))))))))<-Escape Hairpin (shared w/ opsA)
                                    ((((((.((((.......)))).)))))))<-NusA stimulated hairpin
NT 5'-GTCCGTTCGTTTCCACGAATTTGACGGTTTGAAAGAGAAAAGATGTCTCGTTCAAACCGCGAAGCGCACCCGAAATAAAGGAAAATTTCC-3'  JS551
T  3'-CAGGCAAGCAAAGGTGCTTAAACTGCCAAACTTTCTCTTTTCTACAGAGCAAGTTTGGCGCTTCGCGTGGGCTTTATTTCCTTTTAAAGG-5'  JS552
RNA 5'-UUUUAACGUUUCCAC   JS146
```

#### opsA+hp (Class 1)

```
                               (((((((.((((((.......)))))).))))))))<- Escape Hairpin
                                  ((((.((((((.......)))))).)))))<-NusA stimulated hairpin
NT 5'-GTCCGTTCGTTTCCACGTCTTACGGTTTGAATGGGAAAAGATGTCTCGTCCAAACCGCGTAGCGCACCCGAAAGTAACCTCTCGACTTTG-3'  JS553
T  3'-CAGGCAAGCAAAGGTGCAGAATGCCAAACTTACCCTTTTCTACAGAGCAGGTTTGGCGCATCGCGTGGGCTTTCATTGGAGAGCTGAAAC-5'  JS554
RNA 5'-UUUUAACGUUUCCAC   JS146
```

#### opsH+hp (Class 2)

```
                                 (((((..(((((....)))))))))) <-NusA-stimulated hairpin
NT 5'-GTCCGTTCGTTTCCACGTCACCTGACCGGGAGGTACTTTCGTACTCCGGTGTCCCCAAAGAACATCCTTTTGTGAAGGAATCCCCAGAGC-3'JS583
T  3'-CAGGCAAGCAAAGGTGCAGTGGACTGGCCCTCCATGAAAGCATGAGGCCACAGGGGTTTCTTGTAGGAAAACACTTCCTTAGGGGTCTCG-5'JS584
RNA 5'-UUUUAACGUUUCCAC   JS146
```

#### opsF+hp (Class 2)

```
                                 (((((..((((.((...))))))))))) <-NusA-stimulated hairpin
NT 5'-GTCCGTTCGTTTCCACGTAACTTGACCGGGAGGTATGCTTCGCTACTCCGGTGTCCCCAAAGAACATCCTGACGTGAACGACATGACAAC-3'JS523
T  3'-CAGGCAAGCAAAGGTGCATTGAACTGGCCCTCCATACGAAGCGATGAGGCCACAGGGGTTTCTTGTAGGACTGCACTTGCTGTACTGTTG-5'JS524
RNA 5'-UUUUAACGUUUCCAC   JS146
```

### PSC

**Bubble Ultramer Scaffold # 1 from PSC 5'UTR (bold region not from UTR)**

```
NT 5'-GATCGAATCCGTTAACCGATCTGTTCAAATCCGGTGCGAATACTTTTTCTGTACCCGATTATCATAAAACTTAATTTGTAAATTGCTGAAAATAAGGCATGTTTTTTGAATATTCCTGTTTTTAACAAATTTTCATCCTTAGTCATTACTGAAACTTTTTCTTACGAACGTAGTCTTGGAGACAACAGATAGCGATAAAC-3' JS708
T 3' -CTAGCTTAGGCAAGCAAAGGTGCAAGTTTAGGCCACGCTTATGAAAAAGACATGGGCTAATAGTATTTTGAATTAAACATTTAACGACTTTTATTCCGTACAAAAAACTTATAAGGACAAAAATTGTTTAAAAGTAGGAATCAGTAATGACTTTGAAAAAGAATGCTTGCATCAGAACCTCTGTTGTCTATCGCTATTTG-5' JS709
RNA     5'-  UUUUAACGUUUCCAC-3' JS146
```

#### Fig 1e, Extended Data Fig. 4

opsF+hp (Class 2)

```
                           (((((..((((.((...))))))))))) <-NusA-stimulated hairpin
NT  5'-GTCCGTTCGTTTCCACGTAACTTGACCGGGAGGTATGCTTCGCTACTCCGGTGTCCCCAAAGAACATCCTGACGTGAACGACATGACAAC-3'JS523
T   3'-CAGGCAAGCAAAGGTGCATTGAACTGGCCCTCCATACGAAGCGATGAGGCCACAGGGGTTTCTTGTAGGACTGCACTTGCTGTACTGTTG-5'JS524
RNA 5'-UUUUAACGUUUCCAC   JS146
```

#### Fig 2d (Graph Z titration effect Ye activity) – 2023-01-04 expt

opsE+hp

```
                              ((((.(((((.......))))).))))<-NusA stimulated pause hairpin
NT  5'-GTCCGTTCGTTTCCACGATCTTGCAGTTTGGCTGAGAAAAAGAGTCTCACCCAACTGCGAAGCATACCCGGACTTAAGTAAATCACTCAG-3'  JS521
T   3'-CAGGCAAGCAAAGGTGCTAGAACGTCAAACCGACTCTTTTTCTCAGAGTGGGTTGACGCTTCGTATGGGCCTGAATTCATTTAGTGAGTC-5'  JS522
RNA 5'-UUUUAACGUUUCCAC   JS146
```

#### Extended Data Fig. 6a (Cog vs Non-cog Yx on opsA) – 2022-03-23 expt

opsA+hp

```
                         (((((((.((((((.......)))))).)))))))<- Escape Hairpin
                            ((((.((((((.......)))))).))))<-NusA stimulated hairpin
NT  5'-GTCCGTTCGTTTCCACGTCTTACGGTTTGAATGGAAAAGATGTCTCGTCCAAACCGCGTAGCGCACCCGAAAGTAACCTCTCGACTTTG-3'  JS553
T   3'-CAGGCAAGCAAAGGTGCAGAATGCCAAACTTACCCTTTTCTACAGAGCAGGTTTGGCGCATCGCGTGGGCTTTCATTGGAGAGCTGAAAC-5'  JS554
RNA 5'-UUUUAACGUUUCCAC   JS146
```

#### Extended Data Fig. 6b (opsE→A mutants) (2022-05-12 and 2022-06-02 expts)

opsE+hp (Pause Cycling Observed)

```
                   ...((((((((.((.(((((.......))))))).)))))))))...... < Does not form bc structure below forms first
                              ((((.(((((.......))))).))))<-NusA stimulated hairpin
NT  5'-GTCCGTTCGTTTCCACGATCTTGCAGTTTGGCTGAGAAAAAGAGTCTCACCCAACTGCGAAGCATACCCGGACTTAAGTAAATCACTCAG-3'  JS521
T   3'-CAGGCAAGCAAAGGTGCTAGAACGTCAAACCGACTCTTTTTCTCAGAGTGGGTTGACGCTTCGTATGGGCCTGAATTCATTTAGTGAGTC-5'  JS522
RNA 5'-UUUUAACGUUUCCAC   JS146
```

$HP_A$,$[-10:-1]_E$ (Pause Cycling Observed)

```
                         (((((((.((((((.......)))))).)))))))<-Escape Hairpin Disrupted
                            ((((.((((((.......)))))).))))<-NusA stimulated hairpin
NT  5'-GTCCGTTCGTTTCCACGTCTTACGGTTTGAATGGAAAAGATGTCTCGTCCAAACTGCGAAGCATACCCGAAAGTAACCTCTCGACTTTG-3'  JS555
T   3'-CAGGCAAGCAAAGGTGCAGAATGCCAAACTTACCCTTTTCTACAGAGCAGGTTTGACGCTTCGTATGGGCTTTCATTGGAGAGCTGAAAC-5'  JS556
RNA 5'-UUUUAACGUUUCCAC   JS146
```

#### opsE+hp -6 mut (Pause Cycling Observed)

```
                       ...((((((((.((.(((((.......))))))).))))))))...... < Does not form bc structure below forms first
                                 ((((.(((((.......))))).))))<-NusA stimulated hairpin
NT 5’-GTCCGTTCGTTTCCACGATCTTGCAGTTTGGCTGAGAAAAAGAGTCTCACCCAACTGCGtAGCATACCCGGACTTAAGTAAATCACTCAG-3’ JS591
T  3’-CAGGCAAGCAAAGGTGCTAGAACGTCAAACCGACTCTTTTTCTCAGAGTGGGTTGACGCaTCGTATGGGCCTGAATTCATTTAGTGAGTC-5’ JS592
RNA 5’-UUUUAACGUUUCCAC   JS146
```

#### HP$_E$,[-10:-1]$_A$ (Pause Cycling Observed)

```
                       .....((.(((.((.(((((.......))))))).)))).))..
                                 ((((.(((((.......))))).)))) <-NusA stimulated hairpin
NT 5’-GTCCGTTCGTTTCCACGATCTTGCAGTTTGGCTGAGAAAAAGAGTCTCACCCAACCGCGTAGCGCACCCGGACTTAAGTAAATCACTCAG-3’ JS557
T  3’-CAGGCAAGCAAAGGTGCTAGAACGTCAAACCGACTCTTTTTCTCAGAGTGGGTTGGCGCATCGCGTGGGCCTGAATTCATTTAGTGAGTC-5’ JS558
RNA 5’-UUUUAACGUUUCCAC   JS146
```

#### opsA+hp (NO Pause Cycling)

```
                            (((((((.((((((.......)))))).)))))))<- Escape Hairpin
                               ((((.((((((.......)))))).))))<-NusA stimulated hairpin
NT 5’-GTCCGTTCGTTTCCACGTCTTACGGTTTGAATGGGAAAAGATGTCTCGTCCAAACCGCGTAGCGCACCCGAAAGTAACCTCTCGACTTTG-3’ JS553
T  3’-CAGGCAAGCAAAGGTGCAGAATGCCAAACTTACCCTTTTCTACAGAGCAGGTTTGGCGCATCGCGTGGGCTTTCATTGGAGAGCTGAAAC-5’ JS554
RNA 5’-UUUUAACGUUUCCAC   JS146
```

#### Fig 4ab – Antisense DNA expts opsE

```
          3’-TCTCAGCCAAACTGCAAGATCGTG-5’ JS566 neg. control asDNA

                   3’-CTAGAACGTCAAACC-5’ JS545 asDNA
                  3’-GCTAGAACGTCAAAC-5’ JS544 asDNA
                 3’-TGCTAGAACGTCAAA-5’ JS543 asDNA
                3’-GTGCTAGAACGTCAA-5’ JS542 asDNA
               3’-GGTGCTAGAACGTCA-5’ JS541 asDNA
              3’-AGGTGCTAGAACGTC-5’ JS540 asDNA
opsE+hp                          ||||
                                 ((((.(((((.......))))).))))<-NusA stimulated hairpin
NT 5’-GTCCGTTCGTTTCCACGATCTTGCAGTTTGGCTGAGAAAAAGAGTCTCACCCAACTGCGAAGCATACCCGGACTTAAGTAAATCACTCAG-3’ JS521
T  3’-CAGGCAAGCAAAGGTGCTAGAACGTCAAACCGACTCTTTTTCTCAGAGTGGGTTGACGCTTCGTATGGGCCTGAATTCATTTAGTGAGTC-5’ JS522
RNA 5’-UUUUAACGUUUCCAC   JS146
```

#### Fig 4cd – Antisense RNA expts opsB

```
                   3’-CUUAAACUGCC-5’ JS659 asRNA
                 3’-UGCUUAAACUGC-5’ JS694 asRNA
                3’-GUGCUUAAACUG-5’ JS695 asRNA
               3’-GGUGCUUAAACU-5’ JS696 asRNA
                              ((((((((((.((((.......)))).))))))))))<-Escape Hairpin (Shared w/ opsA)
                                 ((((((.((((.......)))).)))))))<-NusA stimulated hairpin
NT 5’-GTCCGTTCGTTTCCACGAATTTGACGGTTTGAAAGAGAAAAGATGTCTCGTTCAAACCGCGAAGCGCACCCGAAATAAAGGAAAATTTCC-3’ JS551
T  3’-CAGGCAAGCAAAGGTGCTTAAACTGCCAAACTTTCTCTTTTCTACAGAGCAAGTTTGGCGCTTCGCGTGGGCTTTATTTCCTTTTAAAGG-5’ JS552
RNA 5’-UUUUAACGUUUCCAC   JS146
```

#### Extended Data Fig. 8a

##### Truncated scaffold for testing opsE without pause hairpin

```
5'-GAGAAAAAGAGTCTCACCCAACTGCGAAGCATACCCGGACTTAAGTAAATCACTCAGGAA-3' JS156
3'-CTCTTTTTCTCAGAGTGGGTTGACGCTTCGTATGGGCCTGAATTCATTTAGTGAGTCCTT-5' JS157
               CUCACCCA JS473
  RNA5'-UUUUUUA
```

#### Extended Data Fig. 8b – ntDNA hp mutations

##### opsE (WT)

```
5'-GAGAAAAAGAGTCTCACCCAACTGCGAAGCATACCCGGACTTAAGTAAATCACTCAGGAA-3' JS156
3'-CTCTTTTTCTCAGAGTGGGTTGACGCTTCGTATGGGCCTGAATTCATTTAGTGAGTCCTT-5' JS157
               CUCACCCA JS473
  RNA5'-UUUUUUA
```

##### opsE_mut1(opsA/B-mimic-1(AT→GC))

```
5'-GAGAAAAAGAGTCTCACCCAACTGCGAAGCgcACCCGGACTTAAGTAAATCACTCAGGAA-3' JS467
3'-CTCTTTTTCTCAGAGTGGGTTGACGCTTCGcgTGGGCCTGAATTCATTTAGTGAGTCCTT-5' JS468
               CUCACCCA JS473
  RNA5'-UUUUUUA
```

##### opsE_mut2(opsA/B-mimic-2(T→C))

```
5'-GAGAAAAAGAGTCTCACCCAACcGCGAAGCATACCCGGACTTAAGTAAATCACTCAGGAA-3' JS469
3'-CTCTTTTTCTCAGAGTGGGTTGgCGCTTCGTATGGGCCTGAATTCATTTAGTGAGTCCTT-5' JS470
               CUCACCCA JS473
  RNA5'-UUUUUUA
```

##### opsE_mut3(opsA-mimic-3(A→T))

```
5'-GAGAAAAAGAGTCTCACCCAACTGCGtAGCATACCCGGACTTAAGTAAATCACTCAGGAA-3' JS471
3'-CTCTTTTTCTCAGAGTGGGTTGACGCaTCGTATGGGCCTGAATTCATTTAGTGAGTCCTT-5' JS472
               CUCACCCA JS473
  RNA5'-UUUUUUA
```

#### Fig 5c,d – opsB to E mutants (-PH only) 2023-04-05 expt

##### opsB-hp (reconstitute -3)

```
      ((((((.((((.......)))).)))))))<-NusA stimulated hairpin
NT 5'-TTTGAAAGAGAAAAGATGTCTCGTTCAAACCGCGAAGCGCACCCGAAATAAAGGAAAATTTCCCCAAG-3' JS654
T  3'-AAACTTTCTCTTTTCTACAGAGCAAGTTTGGCGCTTCGCGTGGGCTTTATTTCCTTTTAAAGGGGTTC-5' JS655
                                   CCGCGAAGC JS666
                      RNA5'-UUUUUU
```

##### opsB -36-34 mut to opsE

```
      ((((((.((((.......)))).)))))))<-NusA stimulated hairpin
NT 5'-TTTGggcGAGAAAAGATGTCTCGTTCAAACCGCGAAGCGCACCCGAAATAAAGGAAAATTTCCCCAAG-3' JS660
T  3'-AAACccgCTCTTTTCTACAGAGCAAGTTTGGCGCTTCGCGTGGGCTTTATTTCCTTTTAAAGGGGTTC-5' JS661
                                   CCGCGAAGC JS666
                      RNA5'-UUUUUU
```

#### opsB -26-24 mut to opsE

```
        ((((((.((((.......)))).))))))<-NusA stimulated hairpin
NT 5'-TTTGAAAGAGAAAAaAgGTCTCGTTCAAACCGCGAAGCGCACCCGAAATAAAGGAAAATTTCCCCAAG-3'  JS662
T  3'-AAACTTTCTCTTTTtTcCAGAGCAAGTTTGGCGCTTCGCGTGGGCTTTATTTCCTTTTAAAGGGGTTC-5'  JS663
                                       CCGCGAAGC  JS666
                        RNA5'-UUUUUU
```

#### opsB -18-16 mut to opsE

```
        ((((((.((((.......)))).))))))<-NusA stimulated hairpin
NT 5'-TTTGAAAGAGAAAAGATGTCTCcacCAAACCGCGAAGCGCACCCGAAATAAAGGAAAATTTCCCCAAG-3'  JS664
T  3'-AAACTTTCTCTTTTCTACAGAGgtgGTTTGGCGCTTCGCGTGGGCTTTATTTCCTTTTAAAGGGGTTC-5'  JS665
                                       CCGCGAAGC  JS666
                        RNA5'-UUUUUU
```

#### **Fig 5d (+PH only) – compares opsB-escape duplex to scaffold "-10:1, -14 ins opsB"**

##### opsE+hp > –10:1, –14 ins

```
                                             (((((.(((((.......))))).)))))<-NusA stimulated hairpin
NT  5'-GTCCGTTCGTTTCCACGATCTTGCAGTTTGGCTGAGAAAAAGAGTCTCACCCAAaCcGCGAAGCgcACCCGGACTTAAGTAAATCACTCA-3'  JS597
T   3'-CAGGCAAGCAAAGGTGCTAGAACGTCAAACCGACTCTTTTTCTCAGAGTGGGTTtGgCGCTTCGcgTGGGCCTGAATTCATTTAGTGAGT-5'  JS598
RNA 5'-UUUUAACGUUUCCAC   JS146
```

##### opsB-escape duplex

```
                                             ((((((.((((.......)))).))))))<-NusA stimulated hairpin
NT  5'-GTCCGTTCGTTTCCACGATCTTGCAGTTTGaaaGAGAAAAgatGTCTCgttCAAaCcGCGAAGCgcACCCGaaataaaggaaaatttccc-3'  JS674
T   3'-CAGGCAAGCAAAGGTGCTAGAACGTCAAACtttCTCTTTTctaCAGAGcaaGTTtGgCGCTTCGcgTGGGCtttatttccttttaaaggg-5'  JS675
RNA 5'-UUUUAACGUUUCCAC   JS146
```

#### **Fig 5e/Supplementary Fig. 3 – opsE to opsB progressive swap +PH – 2022-06-02 and 2023-04-12**

##### opsE+hp

```
                                              ((((.(((((.......))))).))))<-NusA stimulated hairpin
NT  5'-GTCCGTTCGTTTCCACGATCTTGCAGTTTGGCTGAGAAAAAGAGTCTCACCCAACTGCGAAGCATACCCGGACTTAAGTAAATCACTCAG-3'  JS521
T   3'-CAGGCAAGCAAAGGTGCTAGAACGTCAAACCGACTCTTTTTCTCAGAGTGGGTTGACGCTTCGTATGGGCCTGAATTCATTTAGTGAGTC-5'  JS522
RNA 5'-UUUUAACGUUUCCAC   JS146
```

##### opsE+hp > -10:1 opsB

```
                                              ((((.(((((.......))))).))))<-NusA stimulated hairpin
NT  5'-GTCCGTTCGTTTCCACGATCTTGCAGTTTGGCTGAGAAAAAGAGTCTCACCCAACcGCGAAGCgcACCCGGACTTAAGTAAATCACTCAG-3'  JS593
T   3'-CAGGCAAGCAAAGGTGCTAGAACGTCAAACCGACTCTTTTTCTCAGAGTGGGTTGgCGCTTCGcgTGGGCCTGAATTCATTTAGTGAGTC-5'  JS594
RNA 5'-UUUUAACGUUUCCAC   JS146
```

##### opsE+hp > -10:1, -14:-36 opsB

```
                                              ((((.(((((.......))))).))))<-NusA stimulated hairpin
NT  5'-GTCCGTTCGTTTCCACGATCTTGCAGTTTtGCTGAGAAAAAGAGTCTCACCaAACcGCGAAGCgcACCCGGACTTAAGTAAATCACTCAG-3'  JS595
T   3'-CAGGCAAGCAAAGGTGCTAGAACGTCAAAaCGACTCTTTTTCTCAGAGTGGtTTGgCGCTTCGcgTGGGCCTGAATTCATTTAGTGAGTC-5'  JS596
RNA 5'-UUUUAACGUUUCCAC   JS146
```

opsE+hp > -10:1, -14 ins opsB

```
                                 (((((.(((((.......))))).)))))<-NusA stimulated hairpin
NT 5'-GTCCGTTCGTTTCCACGATCTTGCAGTTTGGCTGAGAAAAAGAGTCTCACCCAAaCcGCGAAGCgcACCCGGACTTAAGTAAATCACTCA-3' JS597
T  3'-CAGGCAAGCAAAGGTGCTAGAACGTCAAACCGACTCTTTTTCTCAGAGTGGGTTtGgCGCTTCGcgTGGGCCTGAATTCATTTAGTGAGT-5' JS598
RNA 5'-UUUUAACGUUUCCAC  JS146
```

opsE+hp > -10:1, -14 ins, -36:-34, -18:-16 opsB

```
                                 ((((((.((((.......)))).))))))<-NusA stimulated hairpin
NT 5'-GTCCGTTCGTTTCCACGATCTTGCAGTTTGaaaGAGAAAAAGAGTCTCgttCAAaCcGCGAAGCgcACCCGGACTTAAGTAAATCACTCA-3' JS599
T  3'-CAGGCAAGCAAAGGTGCTAGAACGTCAAACtttCTCTTTTTCTCAGAGcaaGTTtGgCGCTTCGcgTGGGCCTGAATTCATTTAGTGAGT-5' JS600
RNA 5'-UUUUAACGUUUCCAC  JS146
```

opsE+hp > (mostly opsB now) -14 ins, -36:-34, -18:-16, -26:-24 opsB

```
                                 ((((((.((((.......)))).))))))<-NusA stimulated hairpin
NT 5'-GTCCGTTCGTTTCCACGATCTTGCAGTTTGaaaGAGAAAAgatGTCTCgttCAAaCcGCGAAGCgcACCCGGACTTAAGTAAATCACTCA-3' JS672
T  3'-CAGGCAAGCAAAGGTGCTAGAACGTCAAACtttCTCTTTTctaCAGAGcaaGTTtGgCGCTTCGcgTGGGCCTGAATTCATTTAGTGAGT-5' JS673
RNA 5'-UUUUAACGUUUCCAC  JS146
```

opsE+hp > (opsB-escape duplex) -14 ins, -36:-34, -18:-16, -26:-24, dsDNA opsB

```
                                 ((((((.((((.......)))).))))))<-NusA stimulated hairpin
NT 5'-GTCCGTTCGTTTCCACGATCTTGCAGTTTGaaaGAGAAAAgatGTCTCgttCAAaCcGCGAAGCgcACCCGaaataaaggaaaatttccc-3' JS674
T  3'-CAGGCAAGCAAAGGTGCTAGAACGTCAAACtttCTCTTTTctaCAGAGcaaGTTtGgCGCTTCGcgTGGGCtttatttccttttaaaggg-5' JS675
RNA 5'-UUUUAACGUUUCCAC  JS146
```

#### Fig 5f – WT vs hybrid Yx expts

opsE+hp

```
                                  ((((.(((((.......))))).))))<-NusA stimulated hairpin
NT 5'-GTCCGTTCGTTTCCACGATCTTGCAGTTTGGCTGAGAAAAAGAGTCTCACCCAACTGCGAAGCATACCCGGACTTAAGTAAATCACTCAG-3' JS521
T  3'-CAGGCAAGCAAAGGTGCTAGAACGTCAAACCGACTCTTTTTCTCAGAGTGGGTTGACGCTTCGTATGGGCCTGAATTCATTTAGTGAGTC-5' JS522
RNA 5'-UUUUAACGUUUCCAC  JS146
```

opsE+hp > -10:1, -14 ins opsB

```
                                 (((((.(((((.......))))).)))))<-NusA stimulated hairpin
NT 5'-GTCCGTTCGTTTCCACGATCTTGCAGTTTGGCTGAGAAAAAGAGTCTCACCCAAaCcGCGAAGCgcACCCGGACTTAAGTAAATCACTCA-3' JS597
T  3'-CAGGCAAGCAAAGGTGCTAGAACGTCAAACCGACTCTTTTTCTCAGAGTGGGTTtGgCGCTTCGcgTGGGCCTGAATTCATTTAGTGAGT-5' JS598
RNA 5'-UUUUAACGUUUCCAC  JS146
```

opsB+hp

```
                               (((((((((.((((.......)))).)))))))))<-Escape Hairpin
                                  ((((((.((((.......)))).))))))<-NusA stimulated hairpin
NT 5'-GTCCGTTCGTTTCCACGAATTTGACGGTTTGAAAGAGAAAAGATGTCTCGTTCAAACCGCGAAGCGCACCCGAAATAAAGGAAAATTTCC-3' JS551
T  3'-CAGGCAAGCAAAGGTGCTTAAACTGCCAAACTTTCTCTTTTCTACAGAGCAAGTTTGGCGCTTCGCGTGGGCTTTATTTCCTTTTAAAGG-5' JS552
RNA 5'-UUUUAACGUUUCCAC  JS146
```

#### Fig 6, Extended Data Fig. 9, Supplementary Fig. 4 – Exo assay scaffolds

##### opsE+hp—for exo footprinting w/ phosphorothioated NT DNA

```
      ((((.(((((.......))))).))))<-NusA stimulated hairpin
NT 5’-TTGGCTGAGAAAAAGAGTCTCACCCAACTGCGAAGCATACCCGGACTTAAGTAAAT*C*A*C*T-3’ JS612 * = phosphorothioate
T  3’-AACCGACTCTTTTTCTCAGAGTGGGTTGACGCTTCGTATGGGCCTGAATTCATTTAGTGA-5’ JS611
                                   GCGAAGCAU JS457
                 RNA 5’-gcauucaaaga
```

##### opsB+hp—for exo footprinting w/ phosphorothioated NT DNA

```
      ((((((.((((.......)))).))))))<-NusA stimulated hairpin
NT 5’-TTTGAAAGAGAAAAGATGTCTCGTTCAAACCGCGAAGCGCACCCGAAATAAAGGAA*A*A*T*T-3’ JS638 * = phosphorothioate
T  3’-AAACTTTCTCTTTTCTACAGAGCAAGTTTGGCGCTTCGCGTGGGCTTTATTTCCTTTTAA-5’ JS639
                                     GCGAAGCGC JS640


             RNA 5’-gguaauggu
```

##### opsB+hp[-36:-34]+[-26:-24] mut to opsE —for exo footprinting

```
      ((((((.((((.......)))).))))))<-NusA stimulated hairpin
NT 5’-TTTGggcGAGAAAAaAgGTCTCGTTCAAACCGCGAAGCGCACCCGAAATAAAGGAA*A*A*T*T-3’ JS710 * = phosphorothioate
T  3’-AAACccgCTCTTTTtTcCAGAGCAAGTTTGGCGCTTCGCGTGGGCTTTATTTCCTTTTAA-5’ JS711
                                     GCGAAGCGC JS640
                   RNA 5’-gguaauggugg
```

##### opsB+hp[-18:-16] mut to opsE —for exo footprinting

```
      ((((((.((((.......)))).))))))<-NusA stimulated hairpin
NT 5’-TTTGAAAGAGAAAAGATGTCTCcacCAAACCGCGAAGCGCACCCGAAATAAAGGAA*A*A*T*T-3’ JS712 * = phosphorothioate
T  3’-AAACTTTCTCTTTTCTACAGAGgtgGTTTGGCGCTTCGCGTGGGCTTTATTTCCTTTTAA-5’ JS713
                                     GCGAAGCGC JS640
                   RNA 5’-gguaauggug
```

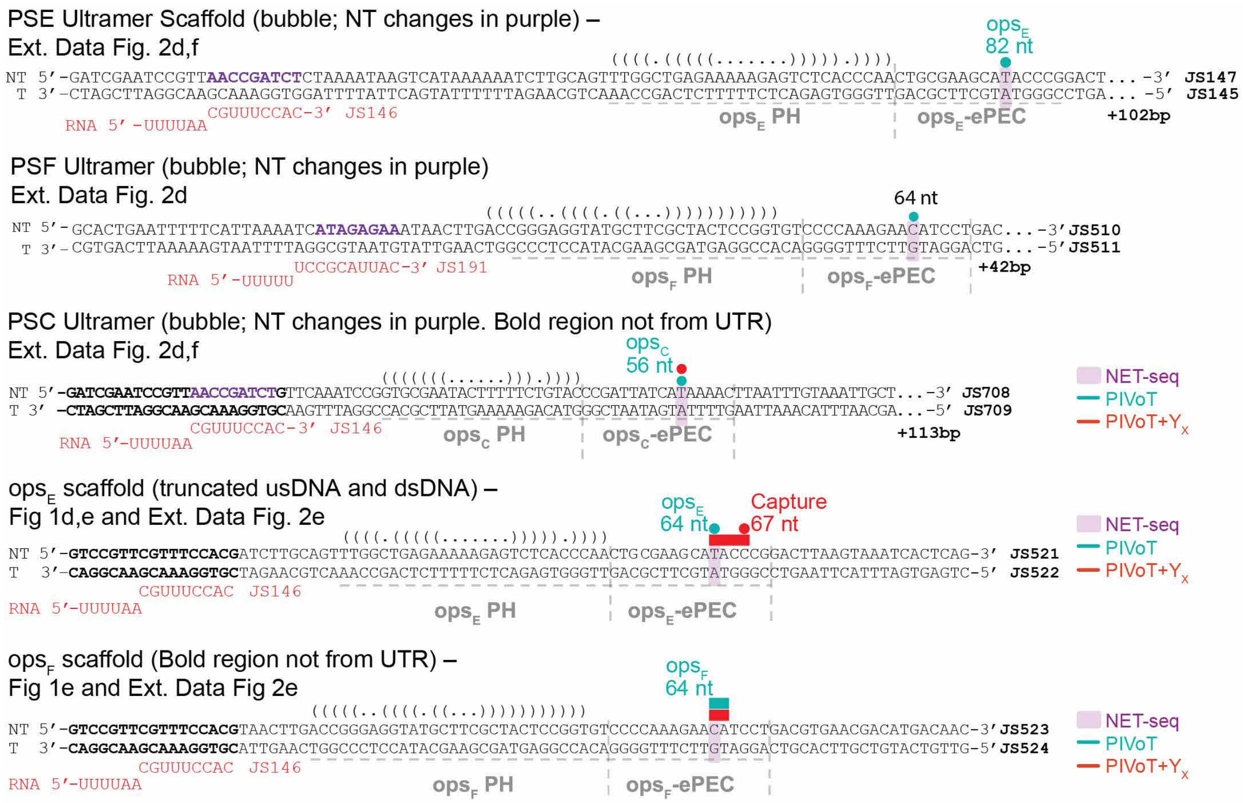


**Supplementary Fig. 1.** Scaffolds used in PIVoT assays of Fig 1d,e and Extended Data Fig. 2.

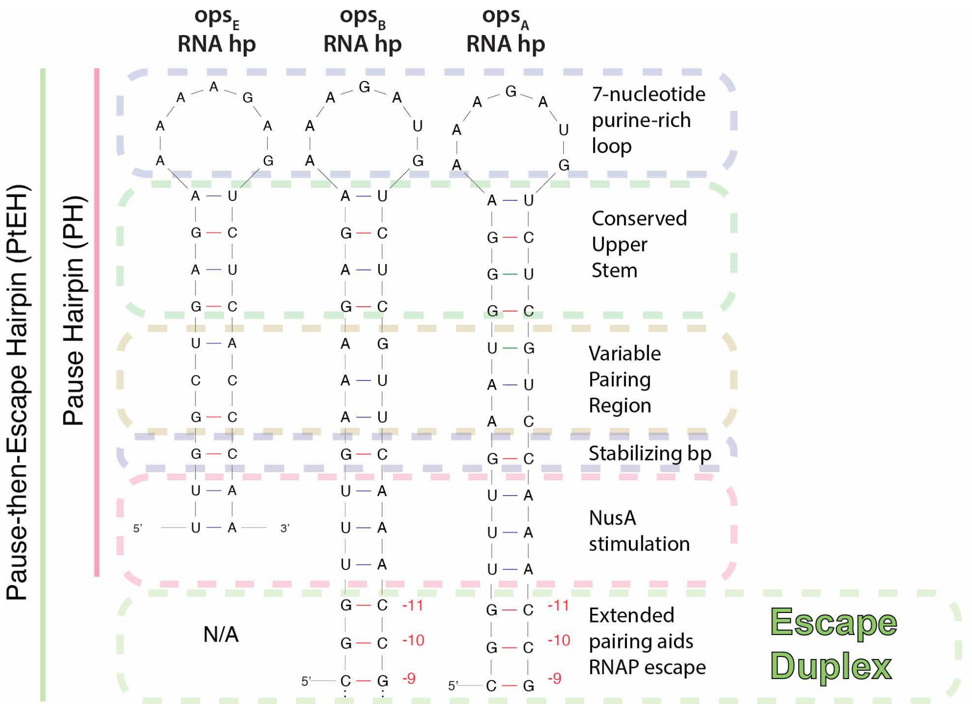


**Supplementary Fig. 2.** Predicted secondary structures of nascent RNA hairpins.

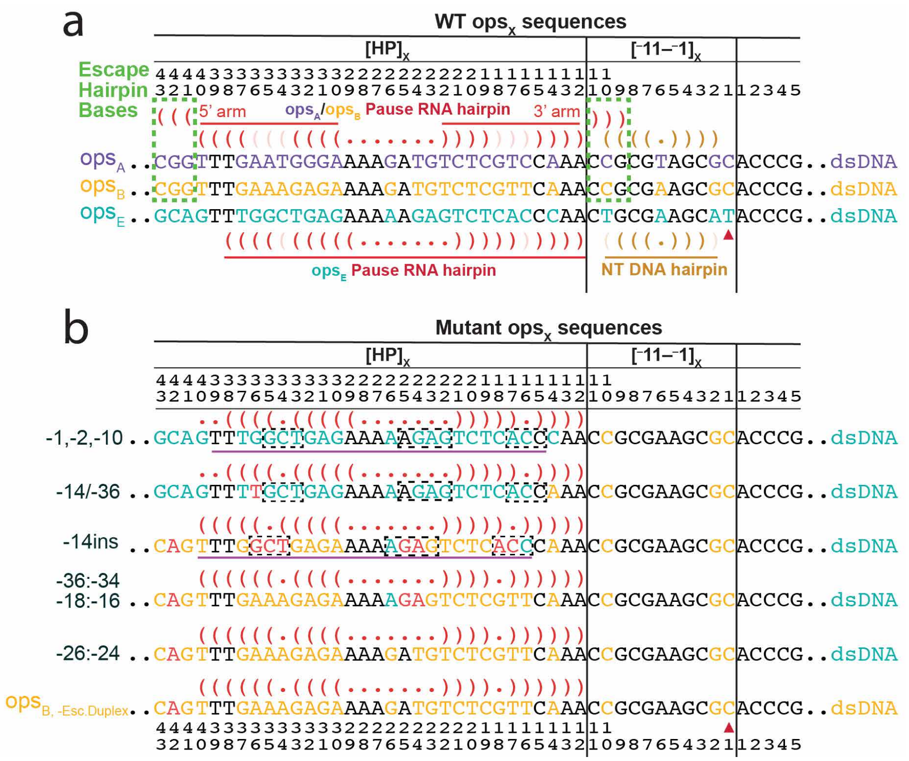


**Supplementary Fig. 3.** WT *ops*$_X$ sequences and variants used for Fig. 5e. **a)** WT *ops*$_X$ sequences highlighted from -43/-38 to +5. Sequences are generally divided into two regions: $[HP]_X$ = hairpin-coding DNA sequence; and $[-11–\,-1]_X$, corresponding to canonical *ops* region. Variable positions in *ops*$_E$ are blue. Variable positions in *ops*$_B$ are orange. Variable positions in *ops*$_A$ are purple. **b)** Regions of *ops*$_E$ progressively mutated to *ops*$_B$ to identify regions of upstream sequence critical for $Y_X$-*ops*$_X$ discrimination. Red nucleotides do not match either *ops*$_B$ or *ops*$_E$. The escape duplex *ops*$_B$ is disrupted by a mutation at –42 to simplify quantitative comparison.

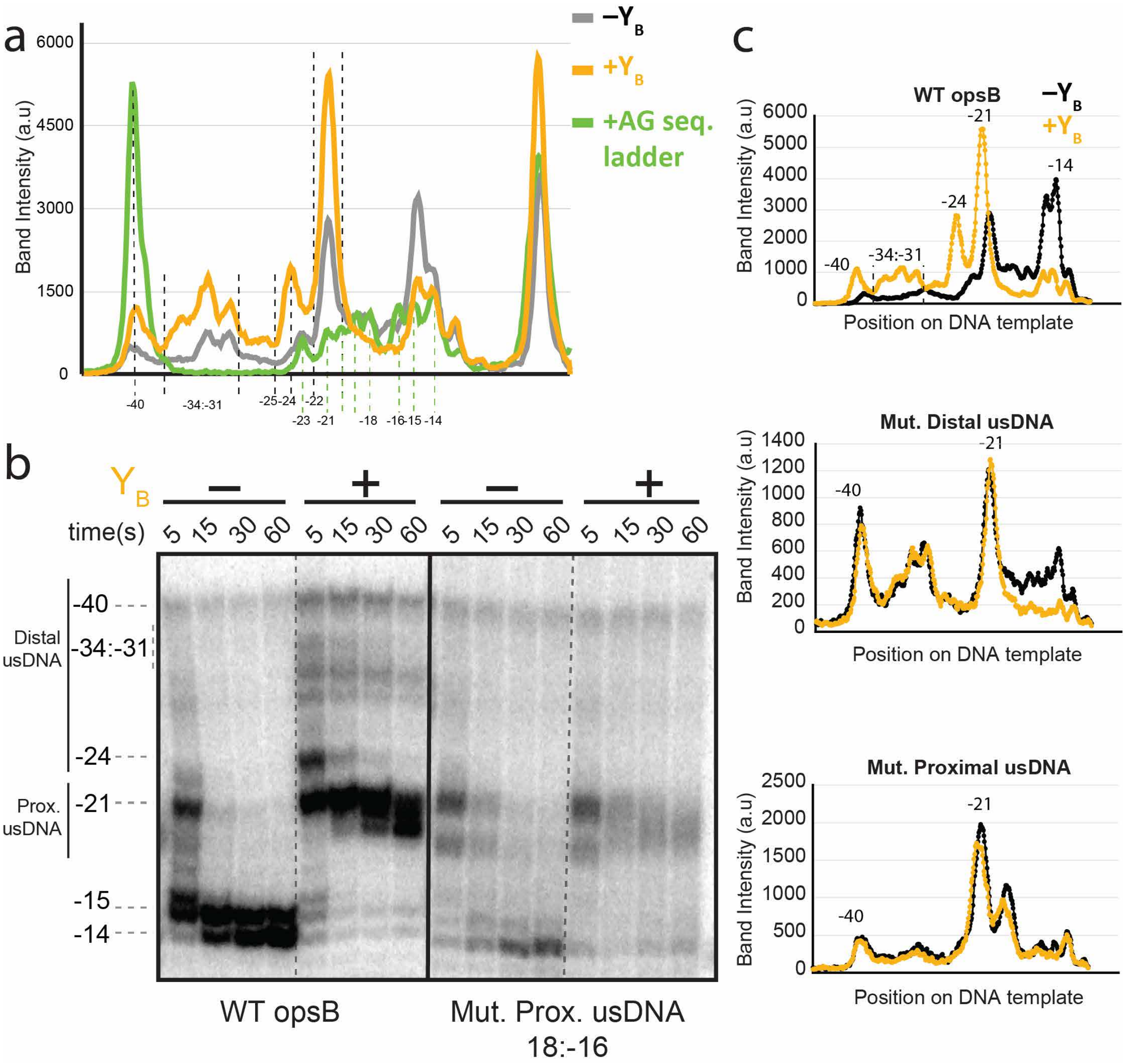


**Supplementary Fig. 4.** Primary data associated with Fig 6c. **a)** Identification of $Y_B$ protection footprints by comparison to AG sequencing ladder. **b)** Full ExoIII-footprinting time course associated with Fig 6c. Representative of experiment performed in at least duplicate. **c)** Pseudodensitometry plots of the 5-second timepoint lane associated with Fig 6c.
